# Identifying phenotype-genotype-function coupling in 3D organoid imaging using Shape, Appearance and Motion Phenotype Observation Tool (SPOT)

**DOI:** 10.1101/2025.11.28.691111

**Authors:** Felix Y. Zhou, Brittany-Amber Jacobs, Adam Norton-Steele, Xiaoyue Han, Linna Zhou, Thomas M. Carroll, Carlos Ruiz Puig, Joseph Chadwick, Xiao Qin, Richard Lisle, Lewis Marsh, Helen M. Byrne, Heather A. Harrington, Xin Lu

**Author notes:** Lyda Hill Department of Bioinformatics, University of Texas Southwestern Medical Center, Dallas, TX, USA. MRC Translational Discovery Unit (TIDU), MRC Weatherall Institute of Molecular Medicine, University of Oxford, Oxford, OX3 9DS, UK. Novo Nordisk Research Centre Oxford (NNRCO), Innovation Building, Roosevelt Dr, Headington, Oxford, OX3 7FZ, UK. Max Planck Institute of Molecular Cell Biology and Genetics, Dresden, Germany (Heather A. Harrington). These authors contributed equally (Felix Y. Zhou, Brittany-Amber Jacobs, Adam Norton-Steele, Xiaoyue Han). **Correspondence:** Correspondence to Felix Zhou, or Xin Lu.

## Abstract

Live cells in tissue are plastic, phenotypically dynamic, and modify their function in response to genetic and environmental perturbations. To unleash the power of live-cell imaging to identify phenotype-genotype-function coupling over time, we report the development of a standardized Shape-Appearance-Motion (SAM) “phenome” and SAM-Phenotype-Observation-Tool (SPOT), that act as an image-“transcriptome” and image-“transcriptome analyzer” respectively, and provide unbiased and comprehensive description of morpho-dynamic phenotypes without prior knowledge. We developed and applied SAM-SPOT to our simulated organoids database with known ground-truth and >1.6 million mouse and human organoid instances with defined genetic and chemical perturbations. SAM-SPOT can effectively and robustly characterize 3D morpho-dynamics from 2D projection videos. Combined with single-cell RNA sequencing, SAM-SPOT revealed that altered WNT signaling, but not mutant RAS or p53, predisposes intestinal organoids to irregular morphogenesis. SAM-SPOT advances biomedical discovery by empowering live-cell imaging to identify phenotype-genotype-function relationships through large-scale and cost-effective label-free live-cell imaging.

## Introduction

The shape and appearance of cells and tissues are crucial biomarkers, routinely used by pathologists to diagnose disease^1,2^. This is because distinctive cell and tissue shapes and appearances can act as proxy readouts of relevant cell functions^3^. Epithelial, neuronal, and immune cells all exhibit distinctive cell morphologies. Secretory or absorptive epithelial cells are highly polarized and adhere together with apical polarity and tight junction protein complexes to form distinctive glandular and intestinal tissue structures, respectively^4,5^. In contrast, squamous epithelial cells form stratified structures, similar to those seen in the skin^4,5^. The appearance of a cell within a particular cell shape will provide additional information about its function and fate. A mitotic nucleus is indicative of a growing, dividing cell whereas a condensed nucleus is a hallmark of a dead cell following apoptosis^6^.

It is becoming increasingly evident that morphological features of complex tissues reflect interactions between cells, their microenvironment, and genetic makeup^7^. Although interrogation of cell phenotype using single cell RNA sequencing (scRNA-seq) has yielded many new biological insights^8-11^, it only partially captures the full phenotypic state and only for a fixed timepoint. Additional phenotypic complexity is afforded by the regulation of protein translation and post-translational modifications that cannot be captured by measuring RNA levels alone. The advent of spatial multiplexing technologies^12-14^ further elucidates the role that cell-cell interactions and the microenvironment play in contributing to diverse cell behavior in tissue^15-17^. In addition to shape and appearance captured by snapshot imaging techniques, live cells are in motion, highly plastic, phenotypically dynamic, and modify their function in response to genetic and environmental perturbations. Therefore, we hypothesized that the wealth of information contained in <u>S</u>hape, <u>A</u>ppearance and <u>M</u>otion (SAM), which can be captured by live-cell imaging, has the potential to assist in the identification of phenotype-genotype-function coupling and to take phenotypic screening to a new level, advancing biomedical discovery, particularly for critical biological functions that manifest over longer-time horizons. To unleash the power of live-cell imaging, we need to 1) detect unbiased and comprehensive image features; 2) interpret these features, and 3) analyze vast and complex image datasets at high throughput.

The development of *ex vivo* organoids as “miniaturized organs” from a single stem cell or a pool of stem cells has emerged as a powerful, physiologically-relevant experimental model to discover new medicines and to study the regulation of cellular plasticity^18^. Organoids present a unique opportunity to undertake exploratory studies at a level which provides information on interactions occurring throughout the surrounding 3D tissue microenvironment. With recent legislation enabling the FDA to approve human trials of new therapeutics in the absence of animal testing, ex vivo models such as organoids will hold increasing importance for the development of new treatments^19^. High-content imaging analyses of 3D organoids have enabled successful multidimensional phenotyping^20-22^ and led to new understanding of the spatiotemporal regulation of regeneration in intestinal tissue^21^. At present, most studies focus on a limited number of *a priori* known or predicted phenotypes and biomarkers^23,24^, compare time-averaged measurements, or focus on a small number of selected timepoints^25-27^. These studies typically 1) rely on macroscale features, such as area, viability and sphericity^22,25,27^; 2) compute features available in standard packages, such as CellProfiler^21,22,28,29^, ImageJ^30^ and Harmony, that may capture only a subset of behaviours; 3) use select fluorescent molecular markers to define and interpret dynamic changes in response to treatment^21,22,31^; and 4) restrict analysis to relatively homogeneous cell cultures, isolated organoids, or developmental stages that can be individually imaged or readily segmented and analyzed^21,22,27,28^. In practice, significant phenotypic heterogeneity is both ubiquitous and a fundamental feature of live cells, organoid cultures and tissues^32^. A large proportion of living cells are not synchronized, except during development; they exhibit individualized growth rates, morphologies, packing densities and movement patterns. This phenotypic heterogeneity can then drive significantly altered and heterogeneous population responses to the same genetic and environmental perturbations^33,34^.

True 3D imaging acquisition of entire organoid cultures at near-isotropic resolution requires sophisticated and customized multi-view light-sheet imaging setups and sample holders^35,36^. These costly setups are not yet suitable to capture all organoids in a large field-of-view in a high content nor high throughput manner. Moreover, light-sheet is restricted to fluorescent markers where capturing full 3D significantly enhances phototoxicity and limits total acquisition time. Therefore, only for a particular biological hypothesis or in situations requiring sub-cellular resolution would an imaging tool that can accurately analyse true 3D time-series data be needed currently. Nevertheless, as organoids are grown in 3D and have emerged as a valuable experimental tool for high-throughput genetic and compound screens, label-free microscopy such as bright-field and phase contrast microscopy and fluorescent microscopy such as confocal modalities that can acquire time-lapse images over a large depth, with small Z-stacks, has broad appeal to both the pharmaceutical industry and research communities, enabling imaging over longer timescales and avoiding the potential side-effects of fluorescent labelling perturbing the native biology.

To respond to this unmet need, and to overcome the associated analytical challenges including (i) ability to characterize 3D dynamics through 2D projection images, (ii) ensuring robustness to artefacts in image acquisition and processing, (iii) applicability to unknown phenotypes and short and long-term acquisitions, we established a standardized SAM “phenome”, composed of 2185 SAM features and developed an associated advanced and universal quantitative image-based analytical framework, SPOT (<u>S</u>AM <u>P</u>henotype <u>O</u>bservation <u>T</u>ool) as a SAM “phenome analyzer”, both of which is applicable to label-free and fluorescent images. We validated SAM-SPOT’s robustness, and effectiveness by developing an organoid simulator and generating synthetic datasets with predefined dynamic behaviour. We then applied SPOT to analyse >1.6 million human and mouse intestine organoid instances derived from various treatment conditions and genetic alterations. We show that SPOT analysis of temporal live-cell imaging of high-content organoid cultures, synergizes with large scale genetic and compound screen to advance our ability to identify molecular switches and modulators that control phenotypic heterogeneity and cellular plasticity.

## Results

### SAM-SPOT platform for 3D live-cell imaging video analysis (Figure 1)

Inspired by single cell sequencing, to realize the potential of high-content timelapse imaging to detect unknown and complex imaging phenotypes, we established an analogous workflow for unbiased and high-throughput analysis of temporal phenotypic heterogeneity in live-cell imaging datasets (Fig 1a). Key to this was the design and construction of SAM-phenome (an image-“transcriptome”) and the development of SAM-Phenotype-Observation-Tool (SPOT) (an image-“transcriptome analyzer”), to comprehensively and quantitatively measure and analyze instantaneous shape, appearance and motion (SAM) characteristics of objects at global, regional and local spatial scales. SPOT analyzes dynamic SAM phenomes extracted from 2D videos in an unsupervised and data-driven manner, and SAM-SPOT provides a single framework for the universal analysis of 2D image and video datasets spanning computer vision and biological cell imaging.

**Figure 1.**
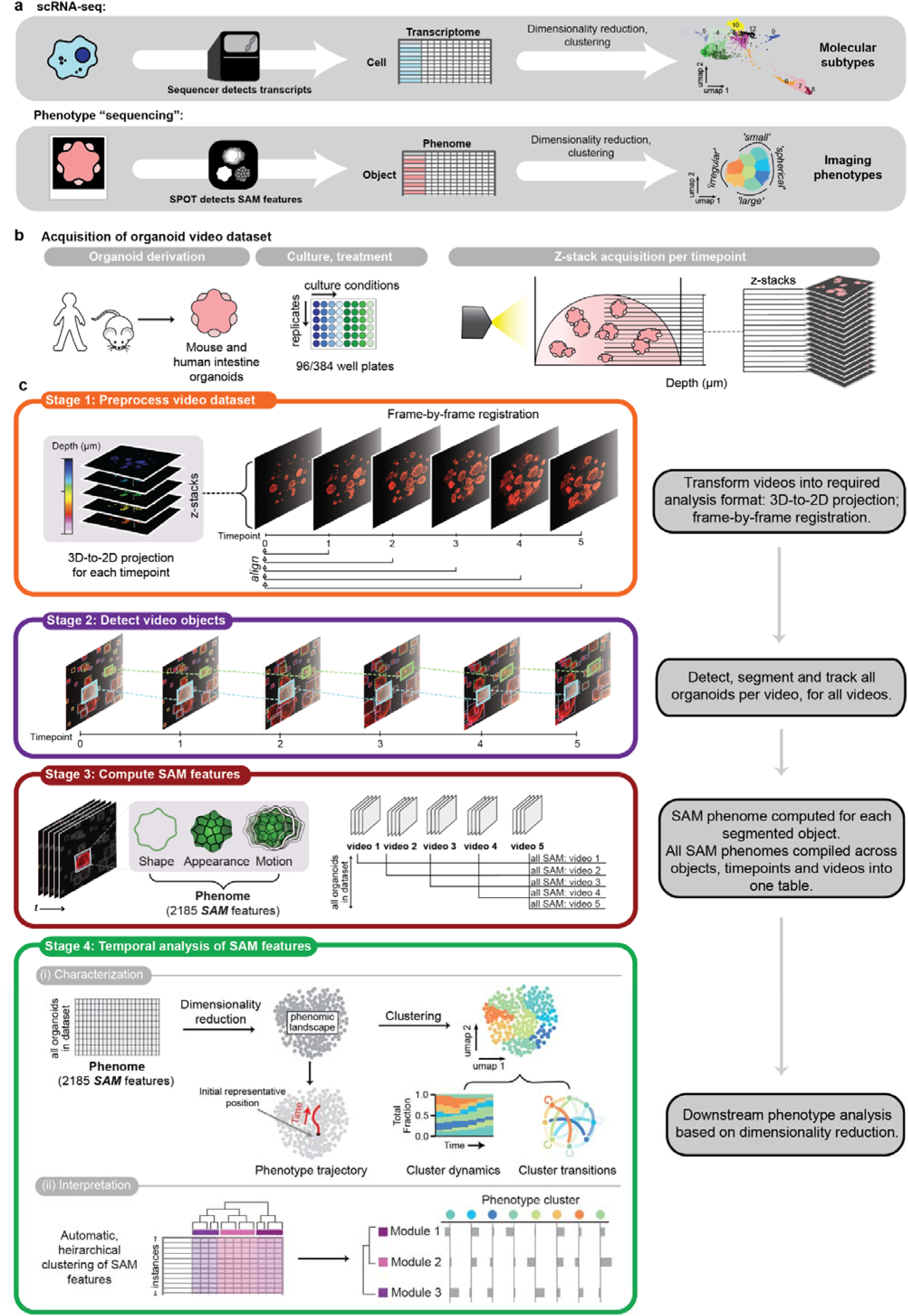
Comprehensive shape, appearance and motion (SAM) characterization enables analysis of phenotypic heterogeneity in organoid models. **a)** Measurement of a standardized high-dimensional library of gene transcripts in single-cell RNA sequencing (scRNA-seq) enables unbiased characterization of the full transcriptome. Parallel to this, our standardized high-dimensional set of SAM features enables the characterization of imaging phenotypes. **b)** Schematic overview of the process of acquiring a suitable organoid video dataset: organoids are derived from relevant tissue samples and cultured before imaging acquisition, where, for each timepoint, z-stack images of defined depth and interval are acquired for a defined period of time. **d)** Overview of the four stages in the SPOT workflow illustrating the steps in stage 1: pre-processing of video dataset (orange box), stage 2: detection, segmentation and tracking of all organoids per video for all videos (purple box), stage 3: computation of SAM phenome for each object instance (red box) and stage 4: the temporal analyses performed using the computed SAM phenomes (green box).

For organoids, <u>S</u>hape is defined as the external boundary of an object and excludes internal composition. It captures the emergent collective behaviour of the constituent cells. <u>A</u>ppearance corresponds to the pattern of image intensity internal to the organoid. It is influenced by the behaviour of each constituent cell e.g. cell death, rendering the lumen of intestinal organoid black and opaque. <u>M</u>otion is the difference in the organoid’s shape and appearance between consecutive timepoints. It captures future behaviour, serving as short-time functional readout to discriminate between organoids with identical shape and appearance. The phenome is constructed by concatenating computed diverse local and global SAM descriptors into one high-dimensional 2185-D vector. Each dimension of the vector is regarded as a feature; its value is the feature expression. This construction enables concepts such as dimensionality reduction, clustering and pseudotime analysis that are essential components of scRNAseq analysis^37,38^ to be adapted for analysing SAM phenomes (Fig. 1a, bottom row).

Herein, we describe how to apply SAM-SPOT to analyze high-content imaging videos of 3D cultures (Fig 1b,c). Organoids were cultured under varying conditions and filmed for a duration of two weeks in 96 or 384-well plates. At each timepoint, a small z-stack captured all organoids over an experiment-specific depth (Fig. 1b). Depending on experiment, our image acquisition uses 11-35 z-slice stacks to sample a total depth of 175-2000 μm. As shown in Fig 1c, Stage 1 of SPOT involves preprocessing this 3D video dataset to be suitable for 2D SPOT analysis. The z-stacks are positioned to capture all organoids within the ECM dome, and do not contain the information for full 3D analysis (Suppl. Fig. 1a, b). Consequently, at each timepoint, the 3D z-stack is first projected to produce a single, 2D image that retains the maximum numbers of organoids within the ECM dome shaped matrigel in-focus. We employed maximum intensity projection for confocal and extended focus projection for label-free microscopy (Suppl. Fig. 1a, b, Methods). The resulting 2D projection video is then registered frame-by-frame to remove drift, particularly translational stage drift in multi-part video acquisitions. In Step 2, every organoid within each video is detected, segmented and tracked (Methods). Spatiotemporal registration during preprocessing removes artefactual motion, ensuring correct organoid tracking throughout the video. Stage 3 computes the universal and comprehensive set of SAM image features that comprise the phenome for every segmented organoid at every timepoint. Stage 4 analyzes all compiled SAM phenomes in the dataset, finding and describing the phenotypic heterogeneity and its temporal evolution to group conditions that lead to the same organoid fate.

Briefly, SPOT first characterizes phenome heterogeneity across all timepoints by: pre-processing all SAM phenomes, removing zero-valued, noisy and non-temporally varying features to retain only the subset of features most relevant and informative to the timelapse experiment. Then, viewing each SAM feature as a genetic transcript equivalent, dimensionality reduction is applied to the filtered SAM phenomes to construct a reduced 2D coordinate space, the SAM phenomic landscape, which maps the salient spatiotemporal phenomic variation between organoids, and crucially enables streamlined temporal analysis. Specifically, SPOT conducts two independent but complementary analyses of temporal evolution: (1) construction of a population-level SAM phenotype trajectory used for cluster conditions, and (2) partitioning of the continuous phenomic landscape into a discrete number of SAM phenotype clusters to understand phenotype cluster dynamics (stacked barplots) and transition probability (by fitting a HMM model, Methods). Lastly, SPOT provides conceptual interpretation at a higher level than individual features by automatically grouping SAM features that co-vary in the dataset into distinct feature sets or SAM modules. The expression pattern of SAM modules can be used to interpret the identified phenotype clusters.

### Generation of an organoid simulator to confirm that SAM-SPOT can effectively and robustly characterize 3D morpho-dynamics from 2D projection videos (Figure 2)

To validate SPOT’s effectiveness and robustness, we developed an organoid simulator to generate two simulated datasets (A and B) with prescribed temporal SAM behaviour. The synthetic organoids are simulated in 3D and then projected into 2D using the same extended focus projection principles as for real acquisitions (Figure 2a). These datasets with ground truth phenotypes enabled us to objectively verify SPOT’s ability to recognise and reconcile: 1) noise and artefacts in image acquisition; 2) inaccuracies in object segmentation caused by spatially overlapping organoids; 3) the effect of fragmented organoid tracks due to incomplete tracking; and 4) the impact of movement in the z-direction on 2D projections.

**Figure 2.**
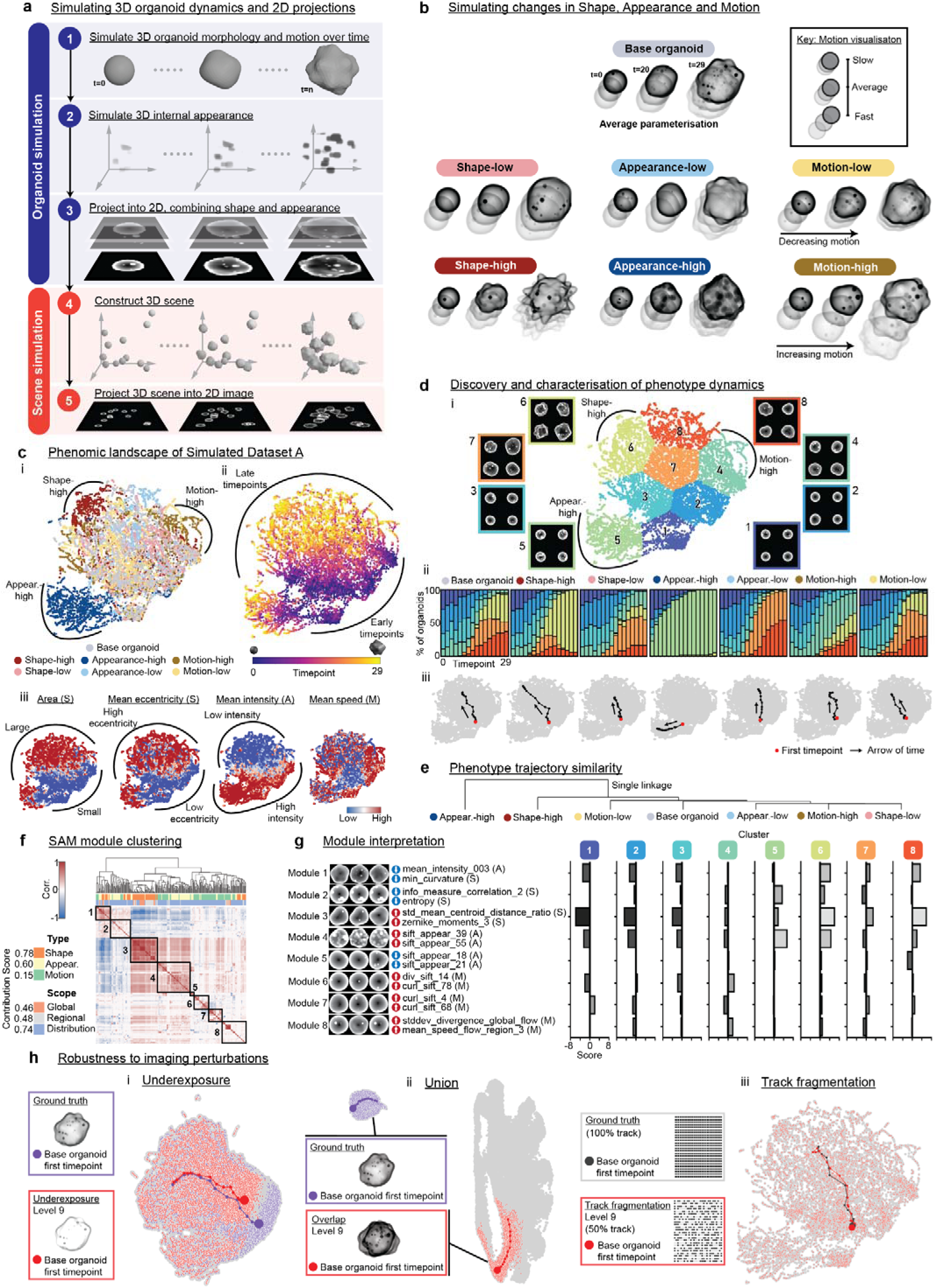
SPOT analysis of synthetic organoid video dataset with prescribed shape, appearance and motion dynamics. **a)** Schematic depicting the main steps in the creation of the synthetic 3D organoid dataset. Individual organoid dynamics were generated by evolving 3D meshes and simulating internal appearance changes across time. For each timepoint, shape and appearance were then independently projected and combined into a single 2D image. 3D scenes are built, and associated 2D projections are generated. **b)** Schematic of the 7 organoid conditions comprising Simulated Dataset A. The first, intermediate and last timepoints are displayed, with motion represented by shadows. **c)** UMAP plots (n=9744) colored to display (i) ground truth phenotypes, (ii) timepoint of each synthetic organoid instance, (iii) size-standardized representative image patches for each phenotype cluster, and (iv) relative expression of select SAM metrics. **d)** Characterization of phenotype dynamics – UMAP phenomic landscape partitioned with phenotype clusters, colored dark blue to dark orange and numbered in ascending order of mean shape eccentricity. Clusters were identified by k-means clustering and the elbow method. For each cluster, four representative images arranged as a 2x2 grid are shown, using their real size for comparison of scale across clusters. (i) Stacked barplots show the temporal changes in phenotype cluster relative frequency across discrete time bins, per phenotype. (ii) Temporal phenotype trajectories show the progression in organoid SAM in each condition across the phenomic landscape across time. Red point indicates the origin of the trajectory at t=0. **e)** Single-linkage hierarchical clustering of phenotype trajectories. **f)** Automated hierarchical clustering of the covariation between SAM features to identify SAM feature modules (left, modules outlined and numbered along the diagonal). **g)** Mean expression of each SAM module for each cluster (as defined in 2d), with bars shaded by relative expression score (right). Each module is depicted with its top three most representative images, its top two driving SAM features and whether the feature is enriched (up arrow) or depleted (down arrow). Individual SAM features are described in depth in our companion methods-focussed manuscript (*Reviewers are requested to contact the Editor for access)*. **h)** UMAP landscape of all datapoints (light grey) for the underexposure, union, and track fragmentation perturbation conditions with the ground truth (purple) and perturbation level 9 (red) emphasized. Trajectories for the ground truth and perturbation level 9 are plotted on top for the ‘base organoid’ condition only. Only perturbation level 9 is emphasized in the tracking error plot as the ground truth is all points.

We generated ‘Simulated Dataset A’ and ‘Simulated Dataset B’ to address Points (1)-(3) and point (4), respectively. ‘Simulated Dataset A’ simulates 3D organoids evolving over time starting from a small, round initial state into 7 phenotypically distinct final states (conditions). 48 organoids were simulated per condition, 30 frames per organoid. To generate the 7 conditions, we generate an initial condition as the ‘base organoid’, with ‘average’ SAM parameter values. Constructing extreme ‘high’ and ‘low’ conditions in each of shape, appearance, and motion in isolation creates the other 6 conditions (Fig. 2b).

SPOT analysis of simulated dataset A successfully distinguished the 7 ground-truth conditions and captured the divergence of each from the base organoid over time (Fig. 2c). The common initial state shared by all conditions occupies a distinct region to the bottom right of the UMAP plot (Fig. 2c i), whilst the three other corners of the phenomic landscape represent the ‘high’ conditions of shape, appearance and motion. The distinct phenotypic end states are distinguishable by a pronounced time-driven trend from bottom-right to top-left universal to all conditions (Fig. 2c ii), and determined by shared features of organoid evolution such as increasing area and eccentricity (Fig. 2c iii).

K-means clustering of the UMAP into discrete regions facilitates analysis that reveals the emergence of condition-specific temporal dynamics (Fig. 2d i). For all conditions, the first temporal bin of the stacked barplot have similar phenotype cluster compositions (Figure 2d.ii). For latter timepoints however, the composition diverges. Clusters primarily attributed to shape, appearance or motion exhibit clear expansion for ‘high’ and contraction for ‘low’. Clusters 6, 5 and 4 expand in the shape-high, appearance-high and motion-high conditions, respectively. Meanwhile the final cluster proportions of ‘low’ conditions share greater representation of the central cluster 7 (representing the average SAM organoid) and contraction of clusters 6 (shape-low), 5 (appearance-low), and 4 (motion-low). HMM cluster transition plots enable the prominent cluster transitions across all simulated organoid tracks per condition to be identified, such as the dominance of transitions to cluster 5 in the appearance-high condition (Suppl. Fig. 2a.ii).

The emergence of phenotypically distinct end states per condition is readily identified by their temporal phenotype trajectories. Starting from same region of the landscape, trajectories diverge, moving over time towards condition-specific regions (Fig. 2d iii), as confirmed by the point density heatmaps of each condition (Suppl. Fig. 2a a iii). Single-linkage clustering of phenotype trajectories reveals each condition is unique, with a general separation of ‘high’ conditions from ‘low’ conditions. All linkage methods identify appearance-high as being notably distinct, due to its unique path across the phenomic landscape (Fig. 2e, Suppl. Fig. 2a b). Applying SAM module analysis, as expected, shows contribution from shape, appearance and motion features across global, regional and distributional scopes, underscoring the importance of having a comprehensive SAM phenome (Fig. 2f). The motion contribution is lower, as it is only varied in the motion-based-conditions, whereas shape and appearance are continuously changing as organoid grow in all conditions. Automatic hierarchical clustering (Methods) identified 8 SAM modules (Fig. 2f). Plotting module expression profiles aided interpretation of phenotype cluster identities and enabled the unbiased discovery of modules that specifically quantify shape (module 3), appearance (module 5) and motion (module 6, 8), as supported by their accompanying top representative images and features (Fig. 2g). As further validation, these modules are most highly expressed in the clusters specific to shape, appearance and motion identified by the stacked barplots e.g., module 4 is highest in cluster 5. To verify SPOT’s robustness, we intentionally and systematically disrupted simulated dataset A with perturbations that mimic typical imaging artefacts and processing errors, at increasing severity levels (Suppl. Fig. 2b). We then assessed the concordance of SPOT analysis using the unperturbed dataset as ground truth using a variety of metrics (Methods). For all imaging-based errors, phenotypic clustering agreement progressively deteriorates with increasing severity level but remains high even at the most severe levels tested – only at level 5-9 of the overexposure perturbation condition is there a marked decline (Fig. 2hi, Suppl. Fig. 2b). The stacked barplot, phenotype trajectory and HMM cluster transition matrix disagreement metrics all remain consistently low (Suppl. Fig 2c). The gradual and modest deterioration of the analysis quality relative to the ground truth demonstrates robustness to even relatively high levels of blur, underexposure, overexposure and physical imaging artefacts, indicating that SPOT analysis tolerates well imaging-based errors that may arise when setting up organoid screens in practice.

In contrast, the simulated segmentation errors are clearly highlighted by the phenomic landscape. All perturbation levels clustered distinctly from the ground truth (Fig. 2h. ii, Supp. Fig. 2b,c). This allows subsetting of the dataset to exclude phenomic landscape regions affected by segmentation error, and to perform SPOT analysis on the remaining subset. Without dataset subsetting we expectedly observe low phenotypic cluster agreement, and relatively high stacked barplot, trajectory, and HMM cluster transition disagreement (Suppl. Fig 2c.v-vii).

Lastly, to address a key challenge in analysing densely growing organoids that often occlude, we simulated incomplete tracking errors arising from occlusion or mispairing of overlapped organoids in practice. By design, phenomic landscape construction of SAM-SPOT is independent of tracking and fragmentation pattern (Fig. 2h iii). Thus, ground truth trajectories for each condition also closely match the corresponding trajectories for each fragmentation level (Suppl. Fig. 2b.d), as corroborated by a low disagreement score. Stacked barplot and HMM transition matrix disagreement scores were similarly low. All these showed that fragmented tracks due to imperfections in organoid tracking has minimal effect on SPOT analysis.

‘Simulated Dataset B’ was constructed to address point (4); whether SPOT analysis is biased by the inability to specifically measure movement in the z-dimension from 2D projection images. We simulate scenes of organoids moving at the same constant speed in 3 conditions: two conditions of free movement in 3D to simulate replicates (denoted XYZ-movement replicate 1 and 2 respectively, Methods) with movement and the third condition with movement restricted to the 2D XY plane (denoted XY-movement). All three conditions overlap in the joint UMAP plot except notably for a small region of the landscope associated with the slowest-moving or static instances, and only populated by the XYZ-movement conditions, (Suppl. Fig.2d a). Visually, the lack of Z-component movement in XY-movement conditions did not generate discernible stacked barplots and phenotypic trajectories from those of XYZ-movment conditions. Nevertheless quantitatively, SAM-SPOT measures sufficient differences in the metrics of stacked barplot and trajectory pairwise disagreement to distinctly cluster the XYZ-movement replicates (Suppl. Fig. 2b ii,iv). Differences between the XY- and XYZ-movement conditions were visually and quantiatively detectable in HMM transition graphs with both XYZ-movement replicates clustering together, (Suppl. Fig.2b. iii). The ability to identify similarity between XYZ-movement replicates across all analyses demonstrate the robustness and applicability of SAM-SPOT to analyze random organoid seeding typical of organoid culture in 3D ECM domes. Notably, XYZ-movement was entirely unrestricted in this simulation. In reality, the ECM dome can spatially restrict organoid motion in the z-dimension, further diminishing the minimal differences SAM-SPOT detects between the XY- and XYZ-movement synthetic conditions. Consequently, a full 3D SAM-SPOT analysis is not required. Altogether, our organoid simulator enabled us to demonstrate the effectiveness and robustness of SAM-SPOT in comprehensively capturing and quantitatively measuring morpho-dynamics of 3D structures from projected 2D video images.

### SPOTting the spatiotemporal dynamics of complex organoids (Figure 3)

Having validated the applicability of SPOT analysis with simulated organoids, we tested its potential to analyze the spatiotemporal dynamics of non-cancerous patient-derived duodenal organoids grown in a 96-well plate as “normal” controls and imaged label-free. The entire SPOT workflow (Fig. 1) was applied to measure organoid response to eight chemotherapy drugs at three different concentrations (Fig. 3a; Methods). 60–100 organoids were seeded per well and all organoids were filmed every hour for 5 days (Suppl. Movie 1). In total, 516,503 organoid instances were segmented from the 96 videos. For each organoid instance, we computed the same 2185-D SAM phenome as for simulated organoids. The SAM phenomes of each organoid instance over all time-points were then compiled and filtered in preprocessing (Methods). UMAP was applied to construct the 2D-SAM phenomic landscape (Stage 4). As with application to 2D single-cell tracking datasets described in our companion methods-focussed manuscript (*Reviewers are requested to contact the Editor for access*), k-means clustering with the elbow method identified seven clusters that capture the phenotypic heterogeneity (Fig. 3b, Methods). The clusters group organoids with similar multidimensional SAM characteristics. Numbered 1–7 and colored from blue to red, clusters are sorted in order of increasing mean eccentricity, a measure of the deviation from the normal, spherical morphology of intestinal organoids cultured in human organoid medium (HOM), a culture condition that we adopt consistently for human organoids throughout this study (Suppl. Table 1, 2). For each cluster, we use SPOT to generate its four most representative images to depict the phenotypic variation across and within clusters (Fig. 3b, top image panels). We show both the actual images to highlight scale differences, and resized ‘standardized’ images where all images are the same size to emphasize appearance differences (Fig. 3b bottom image panels). Qualitatively, cluster 1 (dark blue) and 2 (light blue) organoids are spherical and have bright lumens, whilst cluster 7 (dark orange) organoids are dark.

**Figure 3.**
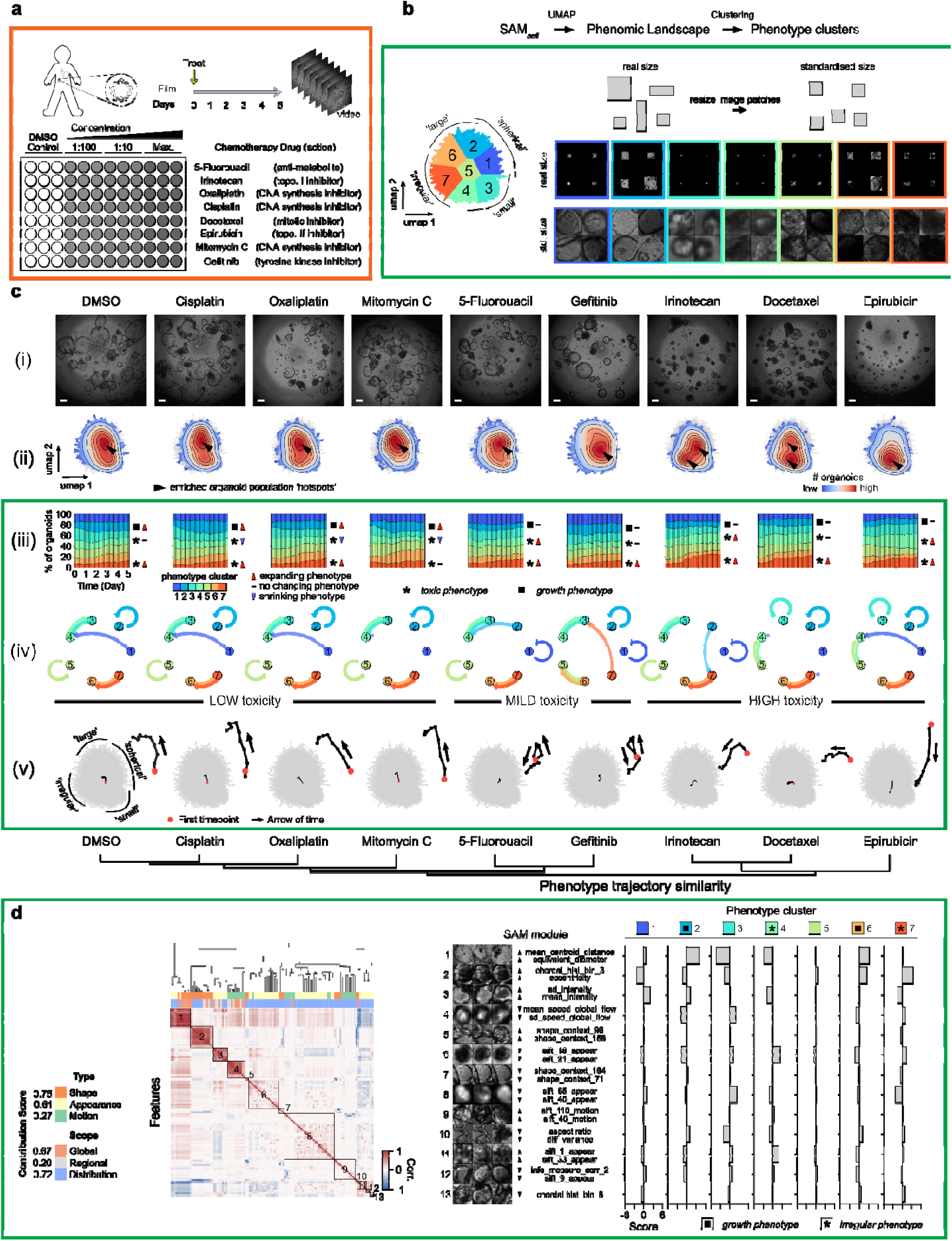
SPOT phenotype analysis of human patient-derived duodenum organoids drug response enables clustering of drugs by pharmacological action. **a)** Patient-derived D2 duodenum organoids were grown from endoscopy biopsies, in a 96-well plate with the specified drug treatment and concentration, then filmed every hour for 5 days without labeling. The whole view of each well was filmed to produce one video. See methods for maximum drug concentrations. For each drug, all videos of all concentrations were pooled together for analysis. **b)** UMAP phenomic landscape using the filtered SAM phenomes from all videos across all drugs and concentrations, n=516,503 points. Each point is a segmented organoid instance from a single timepoint. Landscape is colored dark blue to dark orange and numbered in ascending order according to the mean shape eccentricity of each phenotype cluster, identified by k-means clustering and the elbow method (left). For each cluster, four representative images arranged as a 2x2 grid are shown, using their real size for comparison of scale across clusters (right, top row) and after cropping and resizing to a standardized (std.) size to show organoid appearance (right, bottom row). Illustration of real vs std. size (top right). **c)** Hierarchically clustered chemotherapy drugs according to SAM temporal phenotype trajectories using complete linkage (bottom). (i) final frame of exemplar videos for each indicated drug treatment at maximum drug concentration, scalebar: 250 µm; (ii) UMAP point density heatmaps, same UMAP coordinates as in (b); (iii) stacked barplots showing the temporal changes in phenotype cluster (as characterized in (c)) distribution for each drug, as computed from organoids at 3-hour time intervals; (iv) graph showing the Hidden Markov Model (HMM) inferred transition probability (Methods) of an organoid transitioning to another phenotype cluster in the next timepoint given its phenotype cluster label at the current timepoint. Arrows are colored by the source cluster. The more transparent the arrow, the smaller the probability of the transition. (v) UMAP SAM phenotype trajectories for a density threshold of mean+3 standard deviations (left) and zoom of the trajectory (right) for each drug. On each trajectory, each black point denotes 12-hour increments from the starting point indicated with a red point. A black arrow indicates the directionality of time. **d)** Contribution score of shape, appearance and motion, and global, regional and distributional features (left), automated hierarchical clustering to identify SAM feature modules (middle) and SAM module expression in each phenotype cluster (right) constructed as in Fig. 2f.

Next, we applied SPOT to identify the temporal phenotypic behavior across conditions without ascribing meaning to the identified phenotype clusters *a priori*. Suppl. Movie 1 and snapshots from the final timepoints, at maximum concentration across all drugs, suggest that under the conditions used, dimethylsulfoxide (DMSO, representing control), cisplatin, oxaliplatin, mitomycin C, 5-fluorouracil and gefitinib have lower relative toxicity than irinotecan, docetaxel and epirubicin (Fig. 3c, i). This is supported by visual inspection of the density heatmaps which show the total number of organoid instances treated with the indicated drugs across all three concentrations mapped to local regions of the phenomic landscape (Fig. 3c, ii). Whilst the exact location of heatmap hotspots varies, the results for cisplatin, oxaliplatin, mitomycin C and 5-fluorouracil were similar to DMSO control, with limited impact on organoid morphology, even at the highest concentrations (6, 20, 0.03 and 20 μM, respectively; Fig. 3c, i). However, treatment with gefitinib (up to 0.5 μM) resulted in smaller organoids which clustered at the bottom of the UMAP, compared to the DMSO control. Interestingly, organoids treated with irinotecan (up to 5 μM), docetaxel (up to 0.0125 μM) or epirubicin (up to 4.5 μM) were enriched in both the lower and left areas of the UMAP (clusters 4 and 7), where the organoids appear small, with a dark lumen, and irregular morphology, consistent with accumulation of cell debris and ‘buckling-in’ of the lumen, two stereotypic phenotypic features associated with cell death. Consistent with SPOT analysis, the final frame images depict a field-of-view with more organoids exhibiting these features after treatment with these drugs (Fig. 3c i).

We next applied the SPOT temporal SAM analysis to monitor dynamic changes in phenotypic heterogeneity. The stacked barplots of phenotype cluster frequency over time show substantial expansion of the more toxic-associated phenotype clusters 4 (cyan) and 7 (dark orange) from days 2–3 onwards. In contrast, the low-toxicity drugs show expansion of clusters 1 (dark blue), 2 (light blue), and 6 (light orange) and mildly toxic 5-fluorouracil and gefitinib approximately maintain the fraction of the cluster 2 (light blue) large spherical organoid phenotype (Fig. 3c iii). The HMM cluster transition analysis shows that the dominant cluster transitions are preserved across non-toxic drugs but reveals differences in cluster transitions amongst mildly toxic 5-fluorouracil and gefitinib. Amongst the three toxic drugs, docetaxel and epirubicin appear to have a conserved cluster transition pattern, whilst irinotecan loses the autoregulatory loop of cluster 2 (light blue, Fig. 3c iv, cluster colors the same as in b). Interestingly, this transition pattern is most similar to that of 5-fluorouracil, whose mechanism of action has been reported to involve double-strand DNA breaks^39^. Lastly, the temporal phenotype trajectories were used to summarize the net phenotype behavior for each drug (Fig. 3c v). Predominant organoid growth is indicated by a trajectory from bottom to top in DMSO, cisplatin, oxaliplatin and mitomycin C treated organoids. Increasing toxicity leads to deviation of this trajectory path, with the start moving right-to-left and developing into an opposing trajectory from top to bottom, as for organoids treated with irinotecan and epirubicin. The more toxic, the faster the deviation. 5-fluorouracil and gefitinib showed deviations from the DMSO trajectory; initially going upwards and then coming back down, implying that a longer time was required to generate toxicity and thus that there is a milder toxicity that was not evident from either the final frame snapshots or the heatmap. We hierarchically clustered the trajectories to compare their similarity and the resulting dendrogram (Fig. 3c, bottom) correlates with the drugs’ principal mechanisms of action. Notably, organoids treated with chemotherapeutic drugs affecting DNA synthesis (cisplatin, oxaliplatin, mitomycin C) and inhibiting mitosis (irinotecan, docetaxel or epirubicin) cluster into two distinct groups. We further showed that 5-fluorouracil and gefitinib were functionally similar but have different mechanisms of action, as detected by the HMM transition graph analysis (Fig. 3c iv).

We validated the relative drug toxicity and cluster interpretation by repeating the screen at maximum concentrations with DMSO (control), cisplatin, 5-fluorouracil and docetaxel representing low, mild and high toxicity groupings respectively, and using multi-channel imaging to include markers of cell death (Caspase 3/7 Green) and cell apoptosis (Cytotox Green) (Suppl. Fig. 3a). Using the SAM phenomes of brightfield images only, SPOT generated a phenomic landscape colocalizing organoids with high Caspase and Cytotox expression in the same region. SPOT identified phenotype clusters with the predicted SAM properties and verified they also had the highest Caspase and Cytotox expression. Moreover, unlike biomarkers, the clusters dissect phenotypic heterogeneity associated with two different modes of cell death: i) normal death when organoids grow too big (cluster 2), which we labelled previously as growth phenotype as it indicates organoids grew, or ii) premature drug-induced death (toxic phenotype) whereby organoids are already dead or die at early timepoints without undergoing significant growth (clusters 4, 7). Lastly, clustering of phenotype trajectories showed the same relative toxicity, and was supported by the mean Caspase expression. We performed these screens with organoids cultured in optical-bottom plates (Suppl. Table 3). We also performed a like-for-like comparison with organoids cultured in standard transparent, plastic plates. SPOT analysis generates the same relative drug toxicity (Suppl. Fig. 3b) despite differences in measured SAM (Suppl. Fig. 3c).

Lastly, we used SPOT’s SAM module analysis (Stage 4, step vi) to automatically interpret the phenotypic clusters (Fig. 3d). As expected for toxicity, the contribution scores identified shape and appearance as the dominant SAM features. We found that global features (0.67) were almost equally informative as distributional SAM features (0.72), and that 13 SAM modules summarize the phenotypic variation. Impressively, modules 1–4, corresponding to the first four partitions of the dendrogram, automatically discovered the notions of area, eccentricity, brightness and mean speed without any user supervision. Specifically, SAM module 2 ranks the clusters 1–7 in increasing eccentricity. From the SAM module expression table, we identify clusters 2 (light blue) and 6 (light orange) as normal growth phenotypes based on large size (module 1), relatively bright lumen (module 3) and low moving speed (module 4) in contrast to clusters 4 (cyan) and 7 (dark orange) which are the smallest (module 1), darkest (module 3) and exhibit the highest moving speed (module 4) and also have morphodynamics of non-growing organoids.

SPOT therefore enables fully automated discovery and evaluation of the similarity of organoid phenotypes under different treatment conditions. Moreover, the phenotype trajectories can be used to cluster conditions that exhibit similar phenotype evolution. Together with the stacked barplots and HMM cluster transitions, these results demonstrate how SPOT measures, fingerprints and interprets the spatiotemporal dynamics of phenotypically heterogeneous organoid imaging datasets in a manner applicable to drug screening. We further tested SPOT on a more morphologically complex dataset: label-free live-cell imaging of mouse small intestinal organoid cultures that carry defined genetic alterations. Wild type (WT, p53^+/+^) and p53 null (p53^-/-^) organoids were grown in mouse organoid culture medium with or without inhibitors and imaged every 15 minutes over 5 days (Suppl. Fig 3d, a, Suppl. Movie 2). Valproic acid (V) inhibits histone deacetylase, an enzyme that regulates chromatin function, whereas CHIR99021 (C) inhibits GSK-3β, a serine/threonine kinase that inhibits the WNT signaling pathway. Consistent with a previous report^40^, treatment with V/C enhanced growth and branching morphology in organoids. This effect is largely due to V/C’s ability to alter key signaling pathways, including epigenetic alteration and WNT activation, to induce stem cell expansion and differentiation^40^.

SPOT identified three phenotypically distinct clusters of large area and branching morphologies, cluster 4 (cyan), 7 (light orange) and 8 (dark orange), as depicted by representative image exemplars (Suppl. Fig 3d b). The UMAP density heatmaps of all organoids (WT and p53^-/-^, with and without V/C) show a milder and stronger effect of V/C in p53^-/-^ and WT organoids, respectively, with larger and more branched organoids in both cases (Suppl. Fig 3d c i-ii). This is reflected in the stacked barplots of phenotype cluster frequency over time, which show pronounced increases in the proportion of branched organoid phenotypes, particularly clusters 7 (light orange) and 8 (dark orange) (Suppl. Fig 3d c iii). The HMM cluster transitions reveal large differences among all four conditions but also a conserved cluster 3-to-7 bi-directional transition upon V/C addition (Suppl. Fig 3d c iv). The cluster 3-to-7 transition is consistent with accelerated growth and branching induced by V/C. The cluster 7-to-3 transition is likely because, when the imaged organoid is sufficiently large, the branches ‘buckle’ or fold-in and there is slowed growth such that its state is no longer well described by any of clusters 4 (cyan), 7 (light orange), or 8 (dark orange, Suppl. Movie 2). Again, the phenotype trajectory summarizes the phenotypic heterogeneity and dynamics with respect to the phenomic landscape (Suppl. Fig 3d.c v). All four conditions show a bottom-to-top temporal trajectory, consistent with growth and increased irregularity, with changes occurring most dramatically for WT+V/C, which has the longest and most direct trajectory. Hierarchical clustering of trajectory similarity demonstrates the global effect of V/C treatment, and its stronger effect on WT organoids.

Applying SPOT’s SAM module analysis (Fig. 1, Stage 4), the contribution scores indicate that branching is more strongly associated with shape (0.73) than motion (0.57), with appearance features (0.38) contributing least, (Suppl. Fig 3d.d). Further, branching is best characterized by distributional (0.82) rather than global (0.46) or regional (0.33) features. We found that 17 SAM modules summarize the phenotypic variation. The top representative images and top SAM features, with each feature recorded as being enriched (up arrow) or depleted (down arrow) within a given module, show that module 1 discovers reduced solidity and increased curvature associated with branching clusters 4 (cyan), 7 (light orange), and 8 (dark orange). By contrast, module 2 discovers specific distributional motion pattern features (sift_motion features) that score the clusters according to the extent of growth and branching. As with our drug screen, we again automatically discovered that modules 5 (green), 6 (beige), 3 (turquoise), and 7 (light orange) correspond to area (equivalent_diameter), eccentricity (eccentricity), brightness (intensity) and mean speed (speed_global_flow), respectively. This suggests that these latter SAM characteristics can be used as a consistent reduced set of informative features to monitor intestinal organoid morphodynamics. Examining the SAM expression profiles across clusters, we verify that clusters 4 (cyan), 7 (light orange), and 8 (dark orange) indeed share similar SAM traits and that cluster 3 (turquoise) exhibits a profile associated with reduced growth and branching. These findings are consistent with the HMM predicted cluster 7-to-3 transition where organoid growth and branching significantly slow at later timepoints (Suppl. Movie 2). Henceforth, we use area, eccentricity, intensity and mean speed as targeted SAM features to profile intestinal organoid phenotype clusters, to simplify presentation.

To explore a potential phenotype-genotype connection, we performed bulk RNA-seq on organoid populations, harvested after five days of treatment. Dimensionality reduction plotting of the transcriptomes using multidimensional scaling (MDS) suggests a clear separation of treated *vs* untreated organoids (Suppl. Fig 3d.e). SPOT resolved both the global phenotypic effect of V/C treatment and its differential potency on WT and p53^-/-^organoids (Suppl. Fig 3d.c and dendrogram). This demonstrates that SPOT provides a complementary analysis to molecular sequencing. SPOT not only captures the endpoint outcome but the incorporation of time provides further insight into how the phenotypes develop.

### SPOT reveals a previously unrecognized phenotype-genotype connections (Figure 4)

To showcase the ability of SPOT to detect cellular plasticity using SAM features, we performed detailed phenotyping of mouse intestinal organoids with defined, disease-relevant genetic alterations. Mutant *APC, KRAS and TP53* are well-known drivers of colorectal cancer (CRC) tumorigenesis^41,42^. Inactivating mutations in *APC* are present in 50–83% of sporadic CRC cases^43-45^ and lead to enhanced WNT signaling and stemness of intestinal epithelial cells^46^. *APC*-deficient stem cells acquire growth advantages that facilitate their clonal expansion in intestinal crypts *in vivo*^47^. Activating mutations in *KRAS*, an essential mediator of the MAPK pathway, are present in 35–45% of sporadic CRC cases and lead to the constitutive activation of downstream signaling pathways, promoting cell survival and proliferation^48^. *TP53* is mutated in 40–50% of sporadic CRC cases, most commonly by missense mutations that lead to p53 dysfunction^49,50^. Mutant *TP53* (deletion or missense mutation) carrying cells are often resistant to cell death. Furthermore, mutations in tumor suppressors *APC* and *TP53* and the oncogene *KRAS* are known inducers of cellular plasticity in 2D cell culture^51-54^. Therefore, we hypothesized that organoids carrying mutant *APC*, *TP53* and *KRAS* either alone or in combination might exhibit distinctive morphodynamics that reflect dysregulated signaling pathways and their oncogenic potential. SAM features detected by SPOT in this set of organoids may provide new insights into phenotype-genotype connections.

**Figure 4.**
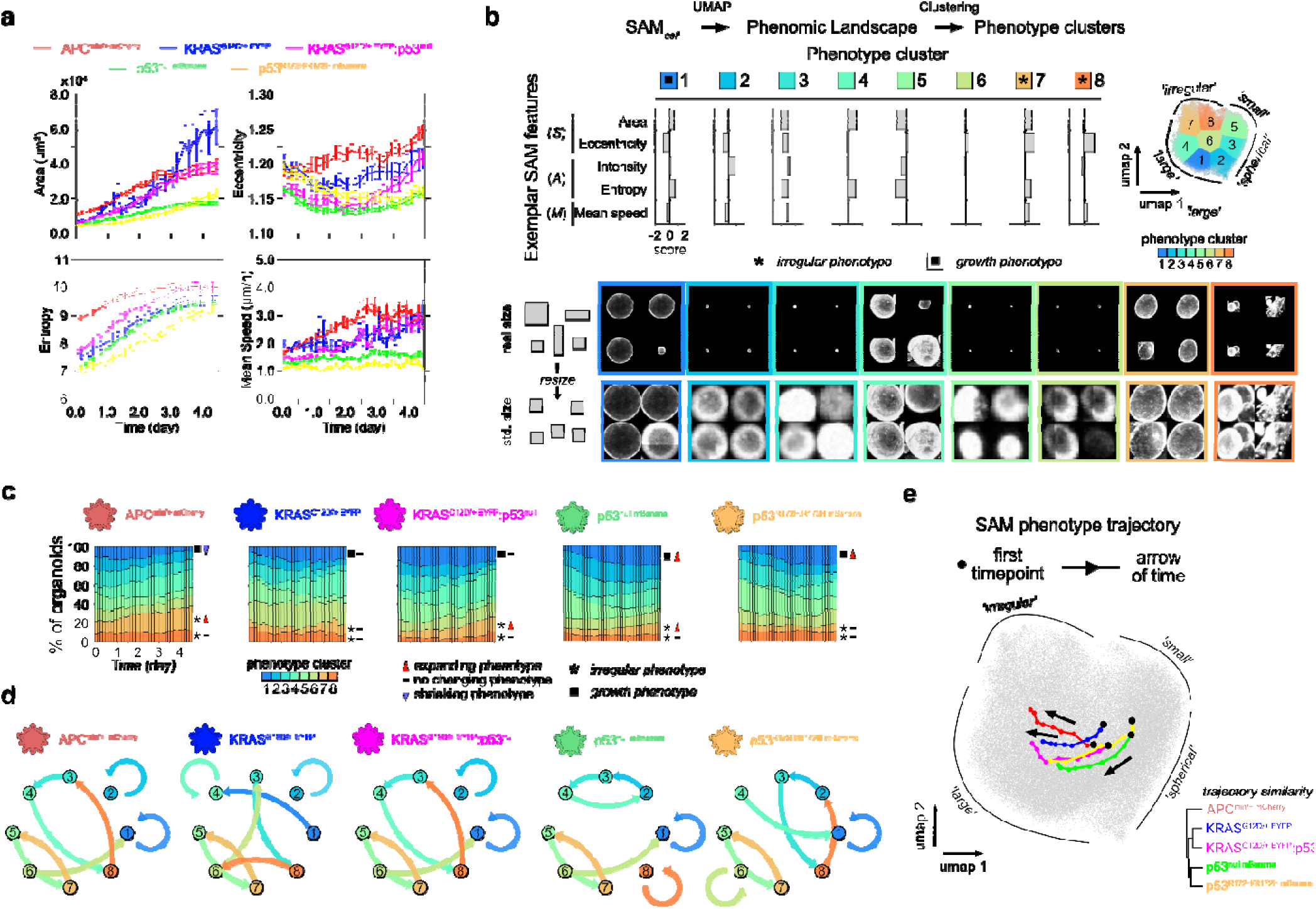
SPOT identifies unique phenotypic differences in genotype defined murine colon organoids. **a)** Temporal measurements of the mean area, eccentricity, image entropy and speed of mutant organoid instances segmented in consecutive time intervals, colored according to genotype. Error bars represent the standard error of the mean. **b)** SPOT characterization of organoids. K-means clustering of the UMAP phenomic landscape constructed using the combined filtered SAM phenomes from a total of 64 videos from five separate repeats carried out in 384-well plates with 1 μL Matrigel droplets, n=79,662 points. Each point corresponds to a segmented organoid instance from a single timepoint. The landscape is colored from dark blue to dark orange and numbered in ascending order according to the mean shape eccentricity of each phenotype cluster identified by k-means clustering and the elbow method (right). Barplots of the SAM score (Methods) of representative global features for each phenotype cluster (left). Representative exemplar images constructed as in Fig. 3b. **c)** Stacked barplot showing the fraction of organoid instances in each phenotype cluster (as characterized in (a)) in consecutive 3 hour time intervals over 0–4.5 days of timelapse acquisition. **d)** Graph showing the Hidden Markov Model (HMM) inferred transition probability (Methods) for an organoid transitioning to another phenotype cluster in the next timepoint given its phenotype cluster label in the current timepoint. Arrows are colored by the source cluster. The more transparent the arrow, the smaller the probability of the transition. **e)** UMAP SAM temporal phenotype trajectories as a function of time from 0–4.5 days, colored according to genotype. On each trajectory, each colored point denotes 12-hour increments from the starting point (black point). Black arrows show the direction of time. Hierarchical clustering of the temporal trajectories (bottom right) uses average linkage.

We derived organoids from the proximal colon of mice carrying mutations in *Apc*, *Tp53* or *Kras* either alone or in combination. Organoid genotypes were confirmed by PCR (Suppl. Fig. 4a.a), labelled with fluorescent proteins and artificially colored in images as indicated: APC^min/+^ (mCherry-red); p53^null^ (mBanana-green); p53^R172H:R172H^ (mBanana-yellow); KRAS^G12D/+^ (EYFP-blue) and KRAS^G12D/+^:p53^null^ (EYFP-purple) (Suppl. Fig. 4a.b). Organoids with the above-mentioned genotypes were first cultured for 72 hours and imaged with confocal microscopy every 2 hours for 4.5 days (Suppl. Movie 3). We segmented and measured a total 79,662 organoid instances pooled from all five independent experiments across all timepoints and all genotypes. KRAS^G12D/+^, APC^min/+^ and KRAS^G12D/+^:p53^null^ organoids grew to larger areas than p53^null^ and p53^R172H:R172H^ organoids throughout the 4 days of filming (Fig. 4a, Suppl. Movie 3). In contrast, all genotypes showed similar entropy (an appearance-based feature that measures the roughness of the image texture) at day 4, despite a higher initial value for APC^min/+^ organoids. Interestingly, APC^min/+^ organoids have higher eccentricity and mean speed during day 1 and day 2 of filming but the double mutant KRAS^G12D/+^:p53^null^ organoids developed similar levels by day 4. These results demonstrate that the choice of a single statistic or timepoint cannot adequately quantify the subtle, dynamic, multidimensional phenotypic differences between organoids of different genotypes (Suppl. Movie 3). Using SPOT analysis (Fig. 1c, Stage 3, 4) we computed the universal 2185-D SAM phenome for all organoids, from all genotypes, and all timepoints. We then applied UMAP to the compiled and filtered SAM phenomes following preprocessing to construct a 2D phenomic landscape, which identified eight phenotypic clusters, ordered by increasing eccentricity (Fig. 4b). As expected, the stacked barplot of phenotype cluster frequency over time showed that the *Kras* and *Tp53* genotypes exhibit similar patterns of phenotype cluster dynamics, whilst APC^min/+^ was different (Fig. 4c). Over time, the most shape eccentric clusters 7 (light orange) and 8 (dark orange, Fig. 4b) expanded the most in APC^min/+^ organoids compared to the other genotypes. Specifically, approximately 15–20% of APC^min/+^ organoids have features identified in the orange clusters during the early stages of filming, with the number of organoids in these two clusters expanding to 30–40% of the total organoids measured at the end of the videos (Fig. 4c). Depth colouring the 2D projections enabled visualisation of depth position of organoids in z-stacks at initial, middle and final timepoints, and revealed that the irregular morphology correspond to organoid flattening, and preferentially occur at the boundary of the ECM dome (Suppl. Fig. 4b). The flattening occurs primarily in APC^min/+^, suggesting a more invasive, migratory behaviour, consistent with APC^min/+^ being commonly the earliest observed of the three mutations in colon cancer^42^. HMM cluster transitions revealed different dynamics for each genotype, with consensus between APC^min/+^ and KRAS^G12D/+^:p53^null^ (Fig. 4d). The temporal phenotype trajectories highlight the early deviation of the APC^min/+^ trajectory leftwards from p53^null^, p53^R172H:R172H^, KRAS^G12D/+^ and KRAS^G12D/+^:p53^null^ genotypes (Fig. 4e). Moreover, hierarchical clustering of trajectories confirmed that APC^min/+^ clustered separately from all other genotypes (Fig. 4e, bottom right). Notably, and in agreement with the HMM cluster transitions, the KRAS^G12D/+^:p53^null^ trajectory deviates towards irregular morphologies at later timepoints; a similar profile to the APC^min/+^ trajectory.

To ensure our observations were not driven by phototoxicity from prolonged laser exposure, we verified that brightfield microscopy acquired timelapse videos of the genotypes show the same expansion of shape eccentric clusters in APC^min/+^ (Suppl. Fig. 4a c-f).

Together, our results demonstrate that SPOT can detect phenotypic changes caused by single-gene perturbations and produces consistent results irrespective of whether videos were acquired with fluorescent confocal microscopy or label-free under brightfield microscopy.

### WNT depletion sensitizes human duodenum organoids to morphological changes like those observed in Apc^min/+^ organoids (Figure 5)

APC deletion leads to elevated WNT signaling and this regulation is evolutionarily conserved from *Drosophila* to humans. This presents a conundrum. If WNT signaling is elevated in Apc^min/+^ mouse organoids, then we expect more spherical morphologies as previously reported^55,56^. Instead, we observed more irregular, flattened organoid morphologies. If the observed phenotype-genotype connection is evolutionarily conserved, an alteration in WNT signaling should have a similar impact on the spatiotemporal dynamics of both mouse and human organoids. We tested this hypothesis using human organoids derived from endoscopic biopsies of non-cancerous D2 duodenum from three individuals as “normal” controls (Methods). To support the expansion and maintenance of Lgr5+ Crypt Base Columnar (CBC) stem cells, these organoids were maintained in HOM, containing >12 growth factors, including WNT, as well as inhibitors and activators of various signaling pathways (Fig. 5a, Suppl. Table 1,2). All organoids cultured in HOM increased in size and retained a spherical morphology (Fig. 5b, top panel). Interestingly, when organoids were transferred from HOM to an ingredient-reduced culture medium containing only <u>E</u>GF, <u>N</u>oggin and <u>R</u>-Spondin (ENR) (Suppl. Table. 1,2), a dramatic morphological change was observed. Organoids grown in ENR medium developed eccentric, flattened morphologies and often ‘merged’ with neighboring organoids (Fig. 5b, middle panel). The flattened ‘sheet-like’ organoid morphology was consistently observed in organoids derived from all three individuals and arose 3–5 days following their transfer to ENR medium (Suppl. Fig. 5a, Suppl. Movie 4). Importantly, this phenomenon was cell density dependent and required a sufficiently high spatial organoid density.

**Figure 5.**
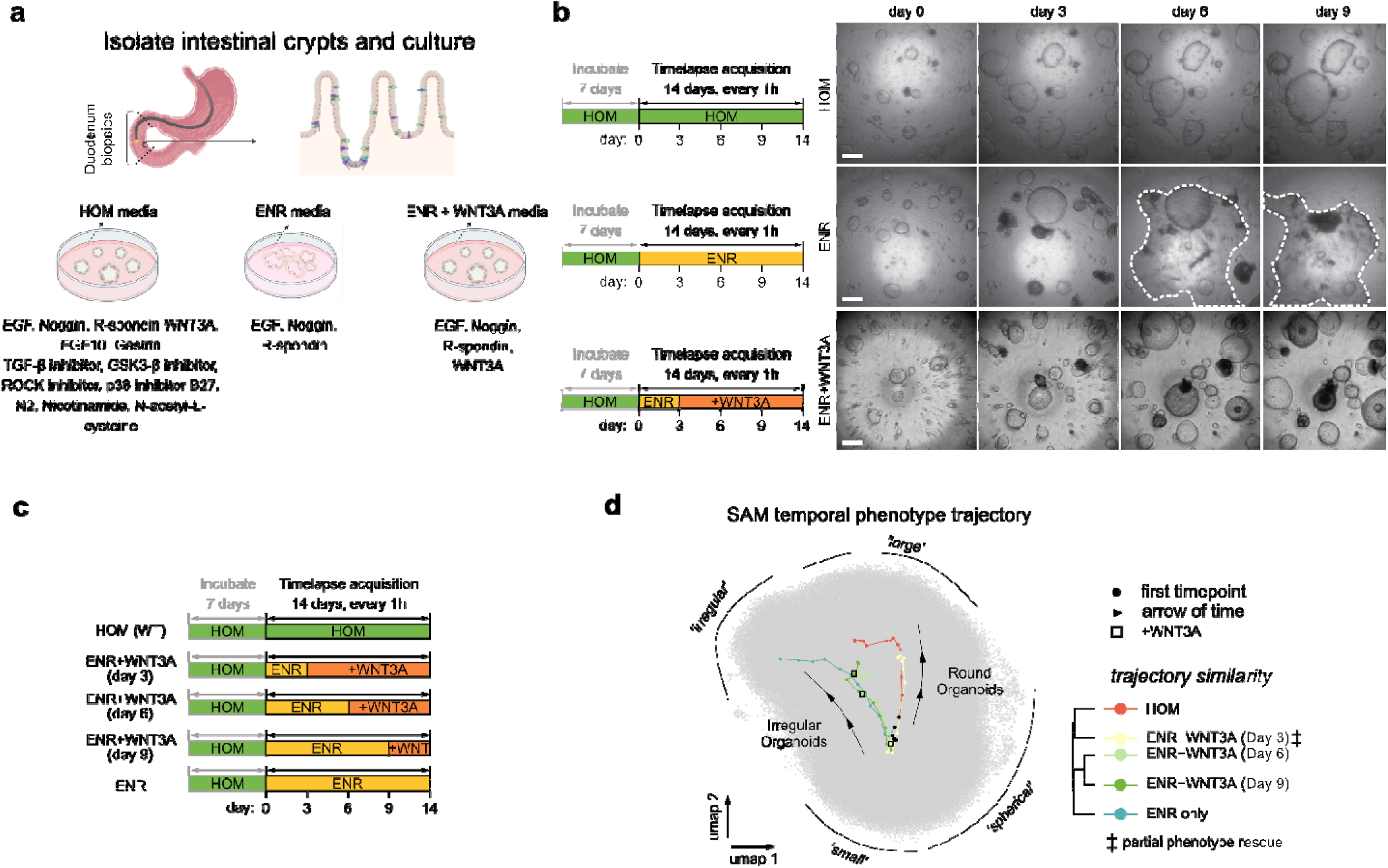
SPOT captures the timing of WNT signaling-induced phenotypic dynamics in patient-derived duodenal organoids. **a)** Patient-derived D2 duodenum organoids grown from endoscopy biopsies and cultured in different medium conditions: human organoid media (HOM), ENR and ENR + WNT3A media. **b)** Timelines of organoid culture and timelapse video acquisition using brightfield microscopy with phase contrast filter (left) and temporal snapshots of organoids cultured in HOM, ENR and ENR + WNT3A media after initial 7-day incubation in HOM media (right). Dashed white contour line from day 6 in ENR outlines the flattened organoids, merging into a sheet. Scalebars: 200 μm. **c)** Illustration of the timeline of treatment and image acquisition for progressively delayed WNT3A restoration to ENR cultured organoids. UMAP SAM temporal phenotype trajectory from 0–14 days. Each colored point denotes one day increments, with the starting timepoint colored black. Black arrows show the direction of time. Black box indicates the timepoint of WNT3A addition. Average linkage hierarchical clustering of the mean trajectories (right).

The observed flattening phenotype is visually reminiscent of Apc^min/+^ murine colon organoids. However, the role of WNT on the morphodynamic changes of human organoids was unexpected. Whereas Apc^min/+^ organoids should have high WNT signaling, ENR medium lacks WNT. To test the direct impact of WNT on the morphological changes of human duodenum organoids, we added WNT3A, a potent inducer of WNT signaling, to the ENR medium on day 3, 6 and 9 after the initial transfer of human duodenum organoids to ENR. Remarkably, restoring WNT3A to ENR medium on day 3 largely prevented the formation of morphologically flattened irregular organoids (Fig. 5b, bottom panel). To quantify the impact of WNT3A restoration, all organoids were grown in 96-well plates for 7 days and live-cell images were captured every hour for the next 14 days (Fig. 5c, Suppl. Movie 4). The spatiotemporal dynamics of a library of >500,000 segmented organoid instances over all timepoints and videos were analyzed by SPOT in the same manner as for mouse colon organoids. We subsequently found seven phenotype clusters (Suppl. Fig. 5b). After quantifying the expression of representative SAM features, organoids cultured in HOM were large and spherical as expected, shown by enriched distribution of cluster 6 (beige, marked ▪). Conversely, ENR organoids exhibited suppressed growth and were substantially enriched in the irregular-shaped cluster 7 (light orange, marked *) (Suppl. Fig. 5b, barplots).

In line with qualitative observations, restoring WNT3A on day 3 substantially reduced the percentage of irregularly shaped organoids in cluster 7 (light orange) and increased their growth in cluster 6 (beige) as observed in HOM (Suppl. Fig. 5b). Importantly, the timing of WNT3A restoration post-transfer to ENR medium was critical for phenotype rescue, with day 3 having the biggest impact, day 6 having little impact and day 9 having minimal impact on reducing the relative proportion of irregularly shaped organoids to spherical organoids, as visualized by the local point density heatmap (Suppl. Fig. 5c). This is further supported by stacked barplots of the phenotype cluster distribution over time (Suppl. Fig. 5d). Restoring WNT3A on day 3 uniquely led to enhanced proliferation (cluster 6 (beige) expansion) and suppressed the expansion of irregularly shaped organoids in cluster 7 (light orange), from day 4 onwards. On days 1–3, prior to restoration, there was little evidence of growth and irregular organoids were seen to expand, in contrast to the rapid growth in HOM conditions. Notably, in all other ENR conditions, neither expansion of cluster 6 (beige) nor regression of cluster 7 (light orange) was observed. The HMM cluster transitions reflect the gradual departure from the control HOM as the transitions become ‘rewired’ (Suppl. Fig. 5e). Intriguingly, we find that the transition graph of restoring WNT3A on day 6 (ENR+WNT3A (Day 6)) best reflects the ENR only condition. Restoring WNT3A on day 9 introduces two new transitions, cluster 5-to-3 and cluster 4-to-2. These transitions are associated with a large-to-small organoid area change which may reflect additional cell death. The ‘rescue’ potential is also succinctly captured in the temporal phenotype trajectories and their clustering, which show close overlap between day 3 WNT3A restoration and HOM conditions (Fig. 5d, right two trajectories). This overlap is distinct from the overlap of the other three conditions: WNT3A restoration on days 6 and 9, and ENR only (Fig. 5d, left three trajectories). These results demonstrate that reduced WNT signaling predisposes human intestine organoids to morphodynamic changes.

### scRNA-seq highlights the potential of SPOT to couple morphodynamics to molecular signaling (Figure 6)

Why is it that irregularly shaped, flattened organoids are enriched in two seemingly opposing conditions: high WNT containing mouse APC^min/+^ organoids and WNT deprived human duodenum organoids? To investigate this question, we performed single-cell RNA sequencing (scRNA-seq). We first performed scRNA-seq on mouse organoids carrying *Apc*, *Kras* or *Tp53* mutations, either alone or in combination, and grown under the same culture conditions as those used for SPOT analysis in Fig. 4. In total, 4,737 single-cell transcriptomes were generated. In agreement with SPOT analysis, cells derived from APC^min/+^ organoids show a unique clustering following dimensionality reduction, separated from all other indicated genotypes (Fig. 6a). PAGA^57^ analysis of all single-cell transcriptomes (n=4,737 cells) identified 12 interconnected clusters (Fig. 6b).

**Figure 6.**
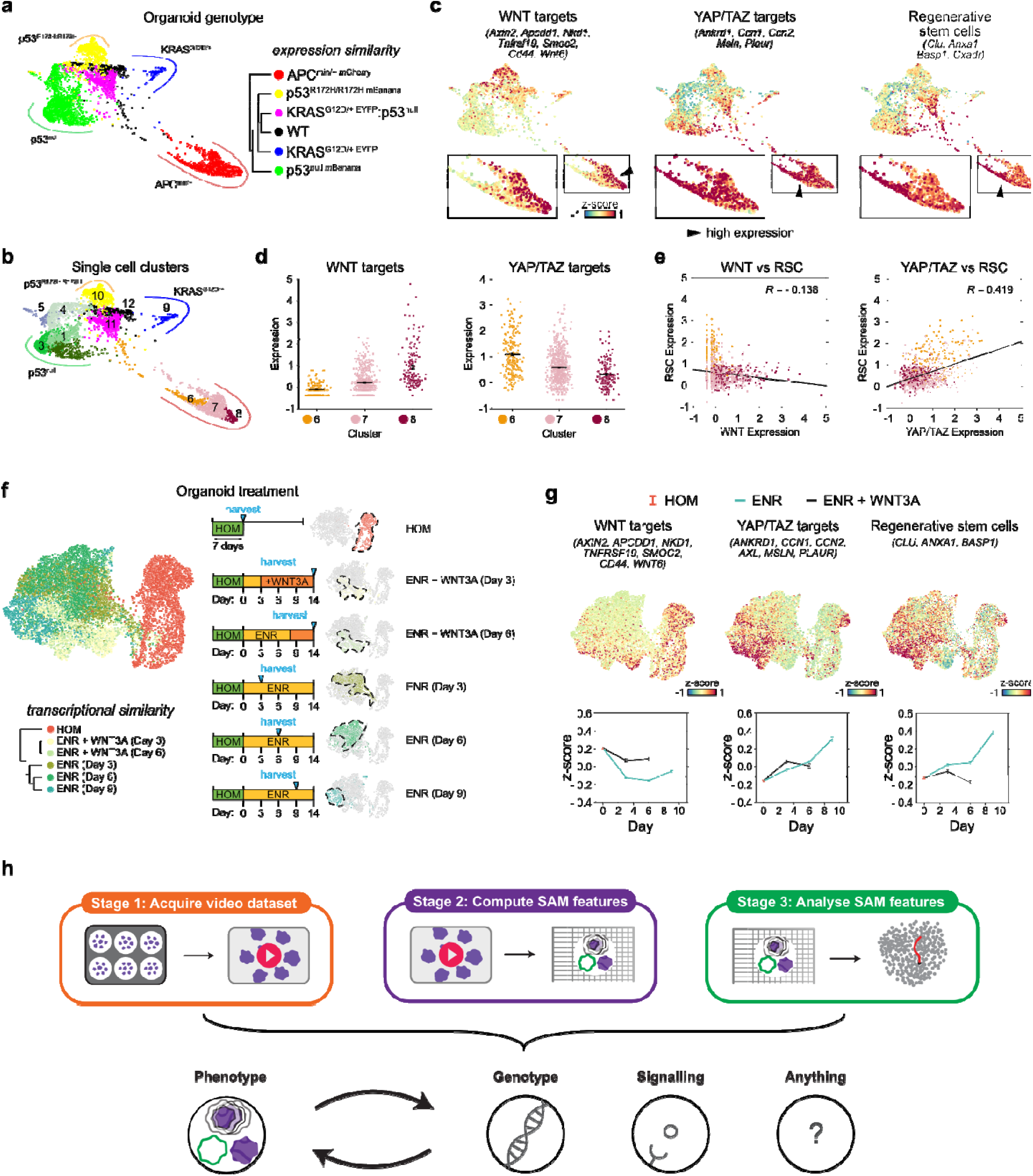
SPOT complements molecular sequencing for scientific and biomedical discovery. **a)** Partition-based graph abstraction (PAGA) ForceAtlas2 (FA) dimensionality reduction plot based on the scaled variance-stabilizing transformed counts of the top 3000 variable scRNA-seq genes (Methods) colored by genotype of mutant mouse colon organoids (left). Hierarchical clustering of individual genotypes based on expression similarity using the one-vs-all differential expression of top 25 marker genes identified by *scanpy* (right). **b)** PAGA FA plot colored by single cell clusters detected by Louvain clustering. **c)** Mean relative gene expression (z-score) of WNT (left), YAP/TAZ targets (middle) and regenerative stem cells (RSC, right) gene sets overlaid onto PAGA FA positions. Boxes show zooms of the FA embedding. z-score was computed on ln(counts+1), the log-transformed raw counts. **d)** WNT (left) and YAP/TAZ (right) associated expression (z-score) in *Apc*-associated single cell clusters 6, 7 and 8 of b). Error bars are plotted as mean ± standard deviation of expression. z-score was computed on ln(counts+1). **e)** Scatter plots of WNT (left) and YAP/TAZ (right) expression on x-axis with RSC expression on y-axis. Linear regression fit (black line) is shown with Pearson’s correlation, *R*. **f)** PAGA initialized UMAP dimensionality reduction plot based on the scaled variance-stabilizing transformed counts of the top 3000 variable scRNA-seq genes (Methods) of human patient-derived duodenum organoids colored by treatment condition (top left). Transcriptional similarity hierarchical clustering using the one-vs-all differential expression of top 25 marker genes identified by *scanpy* (bottom left). UMAP plot separated for individual treatment conditions and corresponding schematic of the organoid culture and the time of harvest for scRNA-seq (right). **g)** Mean relative gene expression (z-score) of WNT (left), YAP/TAZ targets (middle) and RSC (right). Gene sets overlaid onto UMAP positions (top panel), and graph (bottom panel) showing mean z-score ± standard error with respect to the number of days cultured in ENR (ENR) or after WNT3A restoration (ENR + WNT3A). The day of single cell RNA harvest is as indicated in the right panel of f) (Methods). z-score was computed on ln(counts+1). **h)** Utilizing a three-stage workflow and systematic characterization and analyses, SPOT can identify phenotype-genotype, phenotype-signaling and phenotype-‘anything’ couplings where ‘anything’ is determined by the initial treatment and conditions.

Gene expression profiles define cell types with known biological functions and differentiation lineages. Cells expressing high WNT targets and adult stem cell signatures are considered as CBCs whereas cells expressing high YAP targets and regenerative stem cells (RSC) signatures are considered RSCs^58,59^. Under steady-state intestinal homeostasis, WNT-dependent *Lgr5/Olfm4/Axin2-*expressing CBC stem cells dominate^60-63^, whereas RSC populations are key for epithelial repair under stress and injury^64^. These RSCs are *Lgr5* negative, YAP-dependent and express *Ly6a*, *Anxa1* and *Clu^58,65,66^.* As expected, high WNT target-expressing cells are enriched in cell clusters derived from APC^min/+^ and to some extent in p53^R172H:R172H^ organoids (Fig. 6c). Notably, however, the expression levels of WNT targets differ among APC^min/+^ PAGA subclusters (6, 7 and 8), with cluster 6 and 8 expressing the lowest and highest WNT target respectively (Fig. 6d). In contrast to WNT targets, the highest and lowest YAP target expression cells were found in cluster 6 and 8, respectively. Detailed analysis of APC^min/+^ clusters showed that WNT and YAP targets act antagonistically, with the RSC signature correlating strongly to YAP target-expression, and weak anti-correlation to WNT targets (Fig. 6e).

To understand the molecular basis of WNT depletion-induced morphodynamic changes, we also performed scRNA-seq on the human duodenum organoids cultured under conditions and timescales identical to those used for the SPOT analysis in Fig. 5 (Fig. 6f). In total, 8,763 single-cell transcriptomes were generated. UMAP analysis of transcriptional similarity showed two large global groups, corresponding to organoid-derived cells grown in HOM for 7 days before exposure to ENR, or cultured in ENR for 3–9 days (Fig. 6f). Examining the expression of WNT, YAP targets, and RSC signatures in organoid-derived cells as a function of the time of WNT3A rescue, there is an inverse association between high WNT and high YAP target-expressing cells (Fig. 6g). Importantly, the kinetics of RSC signature expression levels coincide with the extent of phenotypic changes detected by SPOT. Thus, WNT-deprived ENR medium induces or accumulates RSCs that lead to the development of morphologically irregular and flattened organoids. This effect can be prevented by restoring WNT3A, and is detected in phenotypic morphodynamic changes by SPOT.

Altogether, these results suggest that the observed ‘flattened’ irregular shaped organoids in both human duodenum and mouse Apc^min/+^ organoids are regulated by the same low WNT and high YAP signaling pathways. This is consistent with YAP as a mechanical sensor in response to environmental perturbation and a key regulator of cell fate. The fact that similar SAM features detected by SPOT are underlined by the same signaling pathways under two seemingly contradicting culture conditions highlights the potential of SPOT as a powerful tool to detect the coupling between SAM phenotype dynamics and molecular signaling (Fig. 6h).

## Discussion

Single cell sequencing and spatial transcriptomic technologies have revolutionised our ability to discover new biology and disease-causing factors at high resolution. However, major limitations to these technologies include the high cost, lack of temporal information, and incompatibility with high-throughput screening. High content label-free live-cell imaging in particular offers unique advantages in bridging the temporal gap and at scale as it is minimally phototoxic, cost-effective, compatible with high-content physiologically-relevant 3D organoid cultures. To unleash the power of label-free-live-cell imaging to advance biological discovery, here we report the development and validation of SAM-phenome and SPOT as a powerful exploratory tool for unbiased detection of genotype-phenotype-function coupling compatible with 3D organoid live-cell imaging, and high-throughput analysis. SPOT extracted >500,000 organoid instances and image features from >9,000 organoids (100 organoids per well of a 96-well plate) in less than 12 hours on a single PC without parallel processing (Intel i7-5820K CPU, 64GB RAM, NVIDIA GTX TITAN Black GPU). SPOT can be applied with fluorescent and label-free microscopy images, with organoids cultured at low density (a few) and high density (hundreds), realizing the possibility to observe more physiological behavior over increased timescales (2 weeks) at high temporal resolution (4 frames an hour).

SPOT assesses phenotype-genotype coupling effectively and efficiently on both simulated and real datasets. On simulated datasets, SPOT remarkably detected the difference between organoids constrained to move only in the XY direction from replicates of unrestricted XYZ movement, despite not being able to measure directly z-movement. On real datasets, independent fluorescent markers of cell death validated that SPOT identifies phenotype clusters and SAM modules relevant to biological processes, revealing two distinct SAM signatures and describing two modes of cell death, which would not be captured by a single marker. Using SPOT, we discovered and characterized a morphological flattening phenomenon in gastrointestinal organoids. Among all organoids examined (>1.5 million human and mouse organoid instances), only a small percentage of organoids underwent morphological changes to irregular and flattened organoids. Such phenomena occurred in APC^min/+^ mouse organoid cultures, albeit only at a low frequency following a few days in culture. The relative fraction of these irregular organoids increased over time and with organoid density, but never became the majority population. Without SPOT’s ability to measure universal image features comprehensively, unbiasedly and quantitatively over time, such time-dependent, multidimensional morphodynamic changes are likely to be missed or ignored using static snapshots or statistical testing based on single feature timeseries. Notably, the irregularly shaped, flattened organoid phenotypes were also highly enriched in WNT-deprived human duodenum organoids and the addition of WNT3A could prevent such a phenotype, in a time-dependent manner. The observed phenotypic similarity led us to hypothesize that a similar genetic/epigenetic make-up might be responsible for the same phenotype in two seemingly opposing conditions: high WNT containing APC^min/+^ mouse organoids and WNT deprived human duodenum organoids. This hypothesis was tested using scRNA-seq analysis. Reassuringly, the results in Fig. 6 illustrate that a cluster of high YAP target and low WNT target expressing cells destined to become regenerative stem cells (RSCs) are mainly detected in genotypes and conditions which produced flattened organoids (APC^min/+^ mouse colon organoids or human duodenum organoids cultured in WNT-deprived ENR for 6 or more days). In contrast, few RSCs or flattened organoids were detected in conditions where proliferation was promoted, such as culturing in HOM medium or in organoids carrying mutant *Tp53* or mutant *Kras*.

Detecting the presence of RSCs in APC^min/+^ mouse colon organoids was unexpected. APC deletion stabilizes nuclear β-catenin, leading to elevated WNT signaling and an enhanced number of CBC stem cells. This led us to ask the following question: what could cause a small percentage of APC^min/+^ cells to downregulate WNT signaling and upregulate YAP and RSC signatures to expand RSCs? Standard organoid formation assays can only measure the growth potential and growth rate of organoids, whereas SPOT can measure the temporal changes in imaging features resulting from alterations in dynamic signaling pathway interactions that cause functional consequences. As a result, SPOT revealed that some APC^min/+^ organoids developed irregular shapes, low proliferation and high motility compared to organoids of WT, mutant *Kras* or mutant *Tp53* genotypes. A possible explanation for this would be that high motility speed and morphological plasticity enable APC^min/+^ organoids to reach the edge of the Matrigel or well bottom more rapidly, where they then experience differential mechanical forces. YAP is known to translocate from the cytoplasm to the nucleus where it forms a complex with TEAD, a transcription factor, turning on YAP targets and activating YAP signaling. High YAP target expression is a signature of RSC expressing cells. The observed strong association between a flattened organoid phenotype and the presence of the RSC cell cluster thus highlights SPOT’s potential as a proxy read out for phenotype-genotype and phenotype-signaling coupling.

Altogether, we have shown that SAM-SPOT is a highly robust and versatile high-throughput and high-content image tool that is applicable to label-free live-cell imaging. SPOT will enhance our ability to use patient-derived primary cultures, including organoids, for hypothesis generation with little to no assumptions. SPOT’s ability to detect complex phenotype-genotype-function coupling will aid in identifying molecular targets of genetic and environmental perturbations, and in combination with “omics” technologies, allow researchers to better understand the spatiotemporal dependence of signaling in disease. Inspired by the ability of single-cell sequencing to detect many previously unknown cell clusters from a highly heterogeneous cell population, we envisage that SPOT will enable us to unleash the power of live-cell imaging-based studies to ultimately advance biomedical research.

## Acknowledgements

This work is mainly funded by the Ludwig Institute for Cancer Research (LICR). FYZ, XH, CRP, TMC, JC, XQ, RL, LZ, XL were funded by the LICR. FYZ was also funded by a EPSRC Life Sciences Interface Doctoral Training Centre EP/F500394/1. BAJ was funded by an Oxford-Radcliffe Scholarship and the Clarendon Fund. LZ received a Cancer Research UK Development Fund. TMC gratefully acknowledges scholarship support from the Rhodes Trust. HAH, HMB and LM were partly funded by LICR and the Centre for Topological Data Analysis, EPSRC EP/R018472/1. HAH gratefully acknowledges funding from EPSRC EP/R005125/1 and EP/T001968/1, a Royal Society University Research Fellowship (RGF\EA\201074) and UF150238. HAH and HMB gratefully acknowledge funding from the Emerson Collective. LM was funded by EPSRC grant EP/R513295/1.

We thank Simon Leedham, Francesco Boccellato, Eric O’Neil, Mary Muers, Kate Dunning, Edward Jenkins, Richard White and Colin Goding for discussion and critical reading of the manuscript. We thank Profs. Gijs van den Brink, Calvin Kuo, and Fiona Powrie for kindly providing cell lines. We also thank the John Radcliffe Hospital, Oxford, and those patients who provided duodenum biopsies.

## AUTHOR CONTRIBUTIONS

Conception: FYZ, XL; Investigation and Formal Analysis of: mutant murine colon organoid creation (BAJ), mutant murine colon organoid culture and microscopy (BAJ, LZ), mutant murine intestinal organoid culture and microscopy (XQ), human duodenum organoid culture and microscopy (XH), human duodenum organoid drug screening (CRP), RNA-seq (BAJ, XQ, TC, JC), RNA-seq analysis: (FYZ, TMC), microscopy (RL), Euler characteristic curve shape descriptor development (LM, HB, HH); SPOT and SAM development and analysis (FYZ); organoid simulation experiments and analysis (ANS). Supervision: XL; Funding acquisition: XL; Writing – Original Draft: XL, FYZ; Writing – Review and Editing: all authors.

## Resource availability

Further information and requests for resources and reagents should be directed to and will be fulfilled by the lead contact, Xin Lu.

## Methods

### Experimental model and subject details

#### Conditioned medium for organoid culture

HEK293T Rspo1-Fc cells (provided by the laboratory of Prof. Gijs van den Brink), HEK293T Nog-Fc cells (provided by the laboratory of Prof. Calvin Kuo, Stanford University) and L Wnt3A (CRL-2647^™^) cells were used to generate R-Spondin-, Noggin-, and Wnt3A conditioned medium respectively (Suppl. Table 1). To produce enough conditioned medium, cells were initially grown to confluency at 37°C, 5% CO_2_, in three T175 flasks containing the relevant selective medium and then expanded into fifteen T175 flasks containing the relevant non-selective medium. HEK293T Rspo1-Fc cells and HEK293T Nog-Fc cells were grown in 50 mL of non-selective medium for 7 days before collection. L Wnt3A cells were grown in 25 mL of non-selective medium for 5 days, followed by medium collection and the addition of another 25 mL of fresh non-selective medium for a further 2 days. The two Wnt3A conditioned medium collections were then pooled together. A HEK293T Wnt3A luciferase reporter cell line (provided by the laboratory of Prof. Fiona Powrie, Oxford University) was used to test the quality of the R-Spondin and Wnt3A conditioned medium. Cells were expanded in the relevant selective medium, plated at confluency in a 24-well plate (in non-selective medium) and allowed to adhere overnight. Cells were then transfected with 0.4 ng/mL of Renilla, using Lipofectamine 2000 (ThermoFisher) and Opti-MEM (ThermoFisher) as the transfection reagents. 5 hours later, 500 µL of conditioned medium was added to each well. The cells were left to incubate overnight, following which the Dual Luciferase® Reporter Assay System (Promega) and GloMax Multi Detection Plate Reader (Promega) were used to determine the luminescence of each well. To test the quality of the Noggin conditioned medium, 500 µL of organoid medium containing the newly made Noggin conditioned medium was placed on growing organoids and the growth monitored for 7–10 days. If the organoids began to die or grow at a slower rate, the batch was discarded. Once tested, all conditioned medium was aliquoted and stored at - 80°C.

#### Human duodenum organoid derivation and maintenance

Duodenum (D2) biopsies were collected from outpatients during endoscopy at the John Radcliffe Hospital, Oxford, with written informed consent (through the Oxford Gastrointestinal Illness Biobank, authorized by Yorkshire & The Humber - Sheffield Research Ethics Committee: 16/YH/0247; laboratory research using the samples authorized by South Central - Oxford C Research Ethics Committee: 09/H0606/78).

Half of each duodenum (D2) tissue biopsy was cut into 2–5 mm^2^ pieces and washed twice with ice-cold PBS. Tissue fragments were washed and incubated with cold 5 mM EDTA/PBS (Fisher Scientific, BP2483) and placed on a roller at 4°C for 15 minutes. Tissue fragments were washed twice more with ice-cold PBS and incubated in TrypLE (ThermoFisher Scientific, 12605010) at 37°C for 30 minutes with constant agitation^67^. Following a further two washes with ice-cold PBS, sedimented tissue fragments were vigorously resuspended in ice-cold PBS and allowed to settle under gravity. The supernatant, enriched with cells released from the tissue fragments, was collected, passed through a 35 µm cell strainer (Corning, 352235) and centrifuged at 200x *g* for 5 minutes. The pellets were embedded in growth factor reduced Matrigel (Corning, 354230) and seeded onto pre-warmed 24-well plates (∼30 µL of Matrigel per well). Matrigel was allowed to polymerize at 37°C for 5 minutes and 500 µL of the human organoid culture medium (HOM, see Suppl. Table 2) was overlaid and replaced every 2 days. To prevent anoikis, Y-27632 and CHIR99021 were added to the medium for the first 2 days. Organoids were kept at 37°C in an atmosphere of 5% CO_2_.

Organoids were passaged every 7–10 days at a ratio of 1:4. Using ice-cold PBS, the organoids were retrieved from the Matrigel and briefly centrifuged. To aid dissociation, the pellet was vigorously resuspended in TrypLE and left at room temperature for 5 minutes. The organoid fragments were then briefly centrifuged, embedded in Matrigel and seeded in a pre-warmed 24-well plate (∼ 30 µL of Matrigel/well). The Matrigel was allowed to polymerize at 37 °C for 5 minutes and 500 µL of HOM was then overlaid and replaced every two days.

The remaining half of the duodenum (D2) biopsy was placed in a cryovial and stored at -80°C.

#### Murine fluorescent mutant colon organoids

Animal work was approved by local ethical review and licensed by the UK Home Office (PPL 30/3451). Animals were bred and housed in individually ventilated cages at the Wellcome Trust Centre for Human Genetics, Oxford. All animal studies described in this study were performed in accordance with guidelines provided by the University of Oxford Institutional Animal Care and Use Committee. All procedures were performed under the Home Office Animal Scientific Procedures Act 1986 guidelines (PPL PP7833295). Unless otherwise stated, 6–10 week-old C57BL/6 mice were used in all experiments.

Organoids used in Fig. 4 were derived using published protocols^61,68^. The proximal colon was removed and was cut into small (∼5 mm) pieces and washed with ice-cold PBS until the supernatant was clear. The method stated for the derivation and maintenance of human duodenum organoids was then followed. See Suppl. Table 2 for mouse organoid medium requirements.

#### Murine mutant duodenum organoids

Organoids used in Suppl. Fig. 3d were derived using the protocol of Sato *et al*.^61^. The small intestine was dissected, opened longitudinally, and washed with ice-cold PBS. The isolated tissue was cut into 5 mm segments and vigorously washed with ice-cold PBS ∼5 times until the supernatant was clear. Tissue fragments were incubated in 2 μM EDTA/PBS and placed at 4°C for 1 hour on a roller. The EDTA/PBS was removed after the incubation and the tissue segments were washed once with ice-cold PBS. The tissue was then washed 4 times with Advanced DMEM/F12 (ADF, ThermoFisher Scientific, 12634028) by vigorous pipetting. The supernatant obtained from these washes was collected and centrifuged for 5 minutes at 200x g. The cell pellet was resuspended in 15 mL ADF and passed through a 70 μm cell strainer. The cells were collected and centrifuged for 2 minutes at 200x g. The pellet, now enriched with intestinal crypts, was embedded in Matrigel, seeded into a pre-warmed 48-well plate (∼20 μL of Matrigel per well), and incubated at 37°C for 10 minutes. 200 μL of HOM was added to each well and the plate was incubated at 37°C in the presence of 5% CO_2_. For passaging, organoids were retrieved from Matrigel using ice-cold PBS and broken up mechanically by passing through a 23G, 5/8” needle. The organoid fragments were centrifuged and washed with ice-cold PBS, and then reseeded in fresh Matrigel. The passage was performed every 7–10 days at a ratio of 1:4.

### Organoid experiments details

#### Chemotherapy drug toxicity screen with human duodenum organoids

Organoids were plated onto 96-well plates as follows. Using ice-cold PBS, the organoids were retrieved from the Matrigel and briefly microcentrifuged. To aid organoid dissociation, the pellet was vigorously resuspended in trypsin and incubated at 37°C for 10 minutes to ensure a high degree of dissociation. The organoid fragments were then briefly microcentrifuged and resuspended in 200 μL of Matrigel. The resuspended organoids were then kept on ice to prevent gelation of the Matrigel. Then, 2 μL of the organoid suspension were seeded inside each well. During this process, the organoid-Matrigel mix was resuspended every 12 wells to ensure an even mix. The Matrigel was allowed to gelate at room temperature for 5 minutes. Once the Matrigel solidified, 100 μL of HOM, pre-warmed at 37°C to prevent the detachment of the Matrigel, was then overlaid and replaced every 2 days. Once the organoids had grown for 6 days inside the 96-well plate, the medium was removed by inversion and washed with pre-warmed PBS at 37°C. After removing the PBS by inversion, the wells were filled with 100 μL of HOM with a titration of chemotherapy drugs.

##### Initial chemotherapy screen (Fig. 3)

The maximum drug concentration values used were as follows: Gefitinib, 0.5 μM; Docetaxel, 12.5 nM; Oxaliplatin, 20 μM; Irinotecan, 5 μM; Cisplatin, 6 μM; Mitomycin C, 0.03 μM; 5-Fluorouracil, 20 μM; Epirubicin 4.5 μM. These concentrations were based on the previously published screening concentrations^69^. All drugs were purchased from Sigma-Aldrich with catalogue numbers as follows: Gefitinib (SML1657-10MG); Docetaxel (01885-5MG-F); Oxaliplatin (O9512-5MG); Irinotecan (I1406-50MG); Cisplatin (479306-1G); Mitomycin C (M4287-2MG); 5-Fluorouacil (F6627-5G); Epirubicin (E9406-5MG). All were solubilized with DMSO, then diluted to working concentration with HOM. The percentage of DMSO for each drug are as follows: Gefitinib (0.010%); Docetaxel (0.025%); Oxaliplatin (0.400%); Irinotecan (0.100%); Cisplatin (0.024%); Mitomycin C (0.003%); 5-Fluorouacil (0.020%); Epirubicin (0.045%). The different conditions were filmed on a Nikon microscope for 120 hours at a frequency of 1 image per hour with the 2x objective. Z-stack images 1000 μm in-depth were acquired at 100 μm intervals (11 steps per acquisition) every hour for 5 days.

##### Chemotherapy screen with additional cell-death related fluorescent biomarkers

SPOT analysis of the initial screen suggested 3 toxicity groups (Fig. 3). We performed additional validation experiments with fluoresecent biomarkers using the subset of control DMSO, Cisplatin (6 μM and 100 μM) representing low-toxicity, 5-Fluorouacil (5FU) (20 μM) representing mild toxicity, and Docetaxel (12.5 nM) representing high-toxicity. The concentrations chosen are the maximum level concentration used in Fig. 3. The organoids were grown as described above for 6 days in a 96-well plate (Suppl. Table 3). Z-stack bright-field images and confocal images (for markers) covering 1000 μm total depth were acquired at 100 μm intervals (11 steps per acquisition) every 4 hours for 5 days. The slower acquisition was used to reduce phototoxicity and to account for the additional imaging due to increased channels. As markers, we used Hoechst 33,342 (50 nM/well, ThermoFisher) for cell viability, consistent with literature^24^; co-stained with either Cytotox Green^70^ (60 nM/well, Sartorius, #4633) for cell death; or Caspase 3/7 Green^24^ (2.5 µM/well, Sartorius, #4440) for cell apoptosis. We visually checked that Cytotox and Caspase stains were only expressed as punctate spots in organoids with positive Hoechst staining. We quantified marker expression in each segmented organoids as the number of punctate spots counted by standard difference of Gaussian blob detection. We found the number of spots was more specific and informative than measuring average intensity^24,70^ which was sensitive to nonspecific background auto fluorescence intensity, and could not discriminate between drugs.

##### Chemotherapy screen using plastic transparent and optical-bottom plates

The organoids were grown as described above for 6 days in a 96-well plate. Culture conditions were identical, only the plates used were either standard plastic transparent, or optical-bottom plates designed for microscopy (Suppl. Table 3). Z-stack bright-field images covering 1000 μm total depth were acquired at 100 μm intervals (11 steps per acquisition) every 4 hours for 5 days.

#### VPA/CHIR99021 treatment

VPA and CHIR99021 for the treatment of murine intestinal organoids (Suppl. Fig. 3d) were purchased from Cambridge Bioscience Ltd and stock solutions were prepared following the manufacturer’s protocols. The working concentrations of VPA and CHIR99021 were 1 mM and 3 μM, respectively.

#### Polymerase chain reaction (PCR)

PCR was performed to confirm the murine colon organoid genotypes using the primers outlined in Suppl. Table 4 and GoTaq Green Master Mix (New England BioLabs^®^ Inc.), according to the manufacturer’s instructions, on the ProFlex PCR System (ThermoFisher Scientific) (Suppl. Fig. 4a.a). Zymo Research Quick-DNA Miniprep Plus Kit (D4068) was used to extract organoid DNA.

#### Lentivirus production

A lentivirus expressing the mCherry and mBanana fluorescent proteins was produced to fluorescently label individual mutant murine colon organoids. Phoenix cells (CRL-3213^™^) were seeded onto a 15 cm plate pre-coated with 0.01% poly-L-lysine and allowed to reach confluency. The cells were then transfected with 15 µg of lentiviral vector (CSII pEF mCherry P2A 3F MCS and CSII pEF mBanana P2A 3F MCS), 12 µg of pCD/NL-BH*DDD (HIV-1 Gag/Pol, Tat, Rev) and 9 µg of VSV-G (Vesicular Stomatitis Virus G Glycoprotein) using Lipofectamine2000 and Opti-MEM as the transfection reagents. 24 hours later, the medium was removed and replaced with 15 mL of fresh medium. After a further 24 hours incubation, the medium was collected, passed through a 0.45 µm filter and stored at 4°C. 15 mL of fresh medium was then placed onto the cells for a further 24 hours, followed by the same collection and filtration. The two collections were then pooled, aliquoted and stored at -80°C.

#### Lentiviral transduction of organoids

Following *in vivo* tamoxifen treatment, KRAS^LSL-G12D/+ CreERT2-EYFP^:p53^Null^ murine colon organoids were confirmed to be EYFP positive by confocal microscopy. To fluorescently label the remaining two mutant organoid populations, APC^min/+^ and p53^null^ organoids were transduced with lentivirus expressing mCherry and mBanana fluorescent proteins, respectively^71^. The medium was removed from at least six confluent wells of the relevant genotype and the organoids retrieved from the Matrigel by resuspension with ice-cold PBS followed by brief microcentrifugation. To aid dissociation, the pellet was vigorously resuspended in TrypLE and left at room temperature for 5 minutes. Organoid fragments were briefly microcentrifuged and the pellet was resuspended in 5 mL of the relevant lentivirus. The organoid/lentivirus suspension was added to a 6-well plate and centrifuged at 600x *g* at room temperature for 1 hour. The plate was then incubated at 37°C for 3 hours. Labelled organoids were collected, embedded in Matrigel and seeded in pre-warmed 24-well plates (∼30 µL of Matrigel per well). The Matrigel was allowed to polymerize at 37°C for 5 minutes and 500 µL of the relevant murine organoid medium was overlaid and replaced every 2 days.

To generate a homogenous population of organoids positive for the fluorescent protein of interest, a MA900 Multi-Application Cell Sorter (Sony Biotechnology) was used. Fluorescence was detected within 24 hours of lentiviral transduction.

To generate p53^R172H:R172H^ and KRAS^G12D/+^ organoids, p53^LSL-R172H^ homozygous mBanana and KRAS^LSL-G12D/+ CreERT2-EYFP^ organoids were infected with Cre Recombinase Adenovirus (Vector Labs Catalogue No. 1045)^71^. Dissociated organoids were infected at a Multiplicity of Infection (MOI) of 50 in 1 mL of organoid culture medium and incubated at 37°C for 3 hours. The infected organoids were then briefly spun down, the pellet embedded in Matrigel and seeded in pre-warmed 24-well plates (∼30 µL of Matrigel per well). This was repeated until the mutation was homogenously expressed throughout the organoids. To note, treatment of KRAS^LSL-G12D/+ CreERT2-EYFP^ organoids with the Cre Recombinase Adenovirus rendered them EYFP positive. During the culture process, routine PCR was carried out, and it was found that a subset of KRAS^G12D/+^ mutant organoids had additionally acquired a p53^null^ mutation. We included these as an additional genotype in our SPOT analysis.

#### Phase-contrast timelapse microscopy of mouse small intestine organoids

Images were acquired using a Nikon TE 2000-E Eclipse inverted microscope, with a 10X/0.3 NA Plan Fluor Ph1 objective. The system was maintained at 37°C in the presence of 5% CO_2_. Z-stack images of 1 mm in depth were acquired at 100 μm intervals (11 steps per acquisition), one frame every 15 minutes.

#### Timelapse confocal microscopy of mouse colon mutant organoids grown in 1 µL droplets

Fluorescent mutant colon organoids, APC^min/+^ mCherry, KRAS^G12D/+^ EYFP, KRAS^G12D/+^:p53^null^ EYFP, p53^null^ mBanana and p53^R172H:R172H^ mBanana were seeded as 1 µL droplets into a pre-warmed 384-well plate (VWR Microplates, Cat No. 736-0149) and overlaid with 40 µL of the appropriate mouse organoid medium. Organoids were grown for 72 hours before being imaged on a Zeiss 710MP confocal microscope using the 10X objective. During the timelapse experiment, z-stack images 1225 μm in depth were acquired at 35 μm intervals (35 steps per acquisition) every 2 hours for 5 days at each of the chosen positions.

#### Label-free timelapse microscopy of organoids

##### Murine colon organoids

Organoids were individually seeded into a pre-warmed 96-well plate (Nunc MicroWell Optical-Bottom Plates, Suppl. Table 3) (∼10 µL of Matrigel per well), overlaid with the appropriate mouse organoid medium and grown for 72 hours.

Each well was then imaged on a Nikon TE 2000-E Eclipse inverted microscope, with a 4X/0.3 NA Plan Fluor Ph1 objective. Phase contrast and brightfield filter were used. The system was maintained at 37°C in the presence of 5% CO_2_. Z-stack images 1000 μm in depth were acquired at 100 μm intervals (11 steps per acquisition) approximately every hour for 5 days.

##### Human duodenum organoids

Human duodenum organoids were harvested and dissociated into single cells following the passaging procedure described above. Cell pellets were resuspended in 500 µL of PBS. Cells were filtered through the cell strainer (70 μm) and counted with the automated cell counter (BIO-RAD). The appropriate cell dilutions were made in Matrigel. Then, the cells were seeded in a pre-warmed 96-well black tissue culture-treated (TC-treated) sterile microplate (PerkinElmer), and plates were incubated for 10 minutes in a cell culture incubator at 37°C and 5% CO_2_ to solidify the Matrigel. HOM was added to each well (200 µL) and refreshed every other day. For WNT experiments (Fig. 5) before performing timelapse video collection, the HOM media was changed to ENR (EGF, Noggin, R-Spondin only) for the ENR imaging conditions. Specifically, group HOM organoids were cultured in HOM culture medium for 14 days. Group ENR+WNT3A (day 3) organoids were under ENR treatment for 3 days and then ENR+WNT3A for 11 days. Group ENR+WNT3A (day 6) organoids were under ENR treatment for 6 days and then ENR+WNT3a for the next 8 days. Group ENR+WNT3A (day 9) organoids were under ENR treatment for 9 days and then under ENR+WNT3A treatment for 5 days. Group ENR organoids were under ENR treatment for 14 days. All images were acquired using a Nikon TE 2000-E Eclipse inverted microscope, with a 10X/0.3 NA Plan Fluor Ph1 objective. Phase contrast and brightfield filter were used. The system was maintained at 37°C in the presence of 5% CO_2_. Z-stack images 1000 μm in depth were acquired at 100 μm intervals (11 steps per acquisition) approximately every 60 min for 5 days (for the chemotherapy drug screen, Fig. 3) and 14 days (for the WNT addition experiments, Fig. 5).

#### RNA isolation, bulk RNA sequencing and analysis

For bulk RNA sequencing, mouse small intestinal organoids were retrieved from Matrigel using ice-cold PBS, washed 2 to 3 times to remove dead cells, and collected using a benchtop centrifuge. Total RNA was extracted with the RNeasy Plus Micro Kit (Qiagen). First-strand cDNA was generated following the poly(A) enrichment protocol, and the resulting cDNA libraries were sequenced as 100 bp paired-end reads on the HiSeq 2500 System (Illumina). Library prep and sequencing were performed by the Oxford Genomics Centre based at the Wellcome Centre for Human Genetics. Sequence alignment used STAR and counts generated by Samtools. Counts per million (cpm) were computed from the raw counts using edgeR (version 3.16.5) and R (version 3.3.3) using default parameters to remove any transcripts which did not have at least two samples with cpm > 2. Default parameters of *calcNormFactors* were used to compute the normalisation library size factors for the reduced count matrix and was used with a prior.count=3 to compute the log_2_(CPM) values. Multidimensional scaling (MDS, using *cmdscale*) was then applied to the sample pairwise Euclidean distance matrix of log_2_(CPM) to generate the two-dimensional MDS plotting coordinates in Extended Fig. 3e.

#### Single-cell RNA sequencing

scRNA-seq was performed on the murine wild type, mutant fluorescent colon organoids and human duodenum organoids to determine the differences in their gene expression profiles.

##### Murine organoids

Organoids were individually seeded in a 384-well plate (as described above) and cultured for 7 days before processing. On day 7, the organoids were retrieved from the Matrigel using ice-cold PBS, centrifuged at 300x *g* for 5 minutes and incubated in TrypLE for 15 minutes. The dissociated organoids were then passed through a 35 µm cell strainer and the flow-through centrifuged at 300x *g* for 5 minutes. The pellet was resuspended in 1 mL Cell Recovery Solution (Sigma Aldrich, DLW354253) and incubated on a roller at 4°C for 2h. Samples were then centrifuged at 300x *g* for 5 minutes and the pellet resuspended in 1 mL 0.04% BSA (in PBS). The pellet was resuspended in 100 µL 0.04% BSA and dissociated organoids were passed through a 35 µm cell strainer. Filtered cell suspensions were then counted using an automated cell counter, diluted to a concentration of 1 million total cells /mL in 0.04% BSA and kept on ice until encapsulation.

scRNA-seq was conducted using a 5’ scRNA-seq gene expression workflow (Chromium Single Cell Immune Profiling, Solution v1.1, 10x Genomics). Following encapsulation of cells using the Chromium Controller, GEM-RT, cDNA amplification, and construction of final libraries was conducted following manufacturer’s instructions. Size profile and concentration of final libraries were assessed by the Agilent 2100 Bioanalyzer (High Sensitivity DNA Kit, Agilent, 5067-4626) and Qubit (BR DNA Assay, ThermoFisher, Q32853), respectively. Sequencing of final libraries was conducted on the Illumina NextSeq 550 10x Genomics recommendations (26 cycles read 1, 8 cycles i7 index, 98 cycles read 2, targeting a minimum depth of 20,000 reads/cell).

##### Human organoids

Organoids were individually seeded into a 24-well plate and cultured for 7 days before treatment (as described above).

For scRNA-seq, Group HOM were collected after the initial 7 days’ culture before further treatment, which is day 0 of the timelapse acquisition. Group ENR+WNT3A (day 3) organoids were under ENR treatment for 3 days, ENR+Wnt3A for 11 days, then collected. Group ENR+WNT3A (day 6) organoids were under ENR treatment for 6 days, ENR+Wnt3a for the next 8 days, then collected. Group ENR organoids were under ENR treatment for a total of 3, 6 and 9 days, respectively, then collected. After being harvested, the organoids were processed in the same manner as for murine organoids. Fig. 6f illustrates the treatment and harvesting times.

### Shape, appearance and motion Phenotype Observation Tool (SPOT) to analyze organoid dynamics

SPOT analysis of 2D organoid projection videos was performed using the SAM phenome in the same manner as described for 2D computer vision and live-cell imaging in our companion methods-focussed manuscript (*Reviewers are requested to contact the Editor for access*). We describe only the specific additional processes introduced at each SPOT stage necessary to analyze an organoid timelapse video with 3D z-stack timepoints.

### Stage 1: Preprocessing

SPOT operates on 2D videos. For each timepoint, the 3D organoid z-stack must be projected to a 2D image. Then, any global translational motion artefact should be removed by temporal image registration to avoid introducing unnecessary errors in motion feature computation.

#### Maximum intensity projection of fluorescent 3D organoid z-stacks to a single 2D image

A maximum intensity projection forms a 2D image whereby each pixel is derived is assigned the maximum intensity value across individual z-slices. For each timepoint, we performed maximum intensity projection of the z-stack using the ImageJ *Z project* menu function. Maximum intensity projection is the default method for fluorescent acquisitions with a cytoplasmic marker where a higher pixel intensity translates to a higher likelihood of sampling an organoid.

#### Extended focus projection of 3D organoid z-stacks to a single 2D image

An extended focus projection forms a 2D image whereby each pixel is derived is assigned the intensity value from its most in-focus z-slice. For each timepoint, we performed extended focus projection of the z-stack using the ImageJ *Stack_Focuser* plugin which assigns an in-focus score based on the image gradient computed by Sobel filters. Extended focus projection is the default method for label-free acquisitions where a higher pixel intensity does not correspond to organoid foreground, and instead we should project the organoid edges.

#### Frame-by-frame registration of 2D projection images to correct translational motion artefacts

Extraneous translational motion artefacts shift all organoids in the next frame, causing erroneous tracking and preventing accurate computation of motion features. As the artefact is global, affecting all organoids in the field-of-view, it can generate many incorrect organoid instances to negatively impact SPOT analysis. We therefore recommend correction through frame-to-frame translation registration. This correction is essential for extended long-time acquisitions whereby regular medium change necessitates multiple imaging sessions (Fig. 3,5, Suppl. Fig. 3d, 5). This commonly results in a discrepancy in stage position before and after the medium change, which manifests as a translational movement. We applied Fourier transform cross-correlation methods^72^ for correction, registering and stitching together the extended focus 2D images frame-by-frame across time periods. Blurry frames were automatically detected as those with a mean edge magnitude one standard deviation less than the mean over all video frames and were removed prior to the registration. For more robust registration, we used the absolute magnitude of difference of Gaussian filtered (img – gaussian(img, sigma=15)) frames as ‘edge enhanced’ images to obtain the correct translational correction. We then apply the movement correction to the raw video frames. As registration has no effect on artefact-free consecutive frames, in practice we always run the frame-by-frame registration.

### Stage 2: detection of video objects

#### Detecting organoids frame-by-frame

We trained a YOLOv3^73^ (from http://github.com/AlexeyAB/darknet) model to detect every individual organoid in an image as a bounding box. Our training used a total of 43,199 organoids extracted from 1534 images comprising label-free and fluorescent confocal microscopy. Images contained both well-separated and cluttered overlapping organoids ranging from a few organoids per image up to ≈100. The model operates on 512 x 512 pixel image input. Input images are resized to 512 x 512 to generate bounding box predictions, whose coordinates are resized to that of the original image. We measure a mean average precision with an intersection-over-union (IoU) cutoff of 0.25, 〖mAP_25_=0.91, and 0.50, mAP_5O_=0.67 on a test dataset of 43,145 organoids extracted from 1437 images similarly comprised of label-free and fluorescent confocal microscopy images with separated and overlapping organoids from a few per image to ≈100.

For an input video, the trained YOLOv3 detector is applied frame-by-frame to detect all organoid in each frame. For multichannel imaging, each colour channel is processed independently. YOLOv3 records each detected bounding box with its (x,y) centroid coordinate, width, height and a detection confidence score (0-1). The detection output is a compilation of individual files for each frame and channel of the list of individual YOLOv3 boxes.

#### Tracking detected organoids across video frames

A custom optical flow based predictive tracker links the individual detected bounding box files across frames into tracklets. Linking is performed for each channel independently. Tracklets are built starting with each detected box in frame 0. The tracker then predicts the coordinates of the bounding box in the subsequent frame, by estimating a rigid motion model (scale, translation, rotation) based on the computed local optical flow^74^ within the box. The predicted box is matched to the actual YOLOv3 detected box in the next frame, i.e. frame 1, via bipartite matching using linear assignment (c.f. Scipy *linear_sum_assignment*). We use the pairwise cost matrix computed from 1-intersection-over-union (IoU) between individual boxes from current and next frame (i.e. frame 0 and 1) for matching. This IoU cost simultaneously accounts for shape, and spatial proximity. An IoU cut-off determines a successful pairing. We use IoU>0.25 i.e. 1-IoU<0.75. The tracklets of successfully matched boxes are extended.

Tracklets with boxes that were unsuccessfully matched in frame 0 are extended opportunistically with the predicted coordinates. Boxes unsuccessfully matched in frame 1 start new tracklets, adding to the pool of tracklets. The process is repeated until the end of the video and all predicted boxes are a member of a tracklet. Running tracklets that have not been successfully matched to a ‘real’ organoid box detected by the trained YOLOv3 detector after a specified number of frames are ‘terminated’ and no longer considered. This tracking process was designed to ensure coverage of all detected organoids throughout the length of the video. Importantly, the bipartite matching between consecutive frames enforces the unique 1-to-1 pairing between an organoid in the current to the next frame. Consequently, more than one organoid cannot be paired to one in the next frame, nor can one organoid be paired with multiple in the next frame.

This means if two organoids fuse, only one of the tracklets is extended and the other will be terminated. Similarly, if two organoids split, the original tracklet terminates and two new tracklets are born. This design aims to ensure that within the duration of the tracklet the same organoid is consistently tracked. We further postprocessed tracklets to remove transient detections. The tracklets were filtered by setting threshold values for 1) minimal lifetime coverage (measured as a fraction of tracklet length) and 2) mean YOLOv3 confidence detection score of boxes within the tracklet. An attention UNet^75^ operating at 64x64 pixels was separately trained on a custom in-house label-free and confocal microscopy organoid image patches cropped from randomly sampled video frames to segment and produce individual organoid boundaries from the detected organoid tracklets. Extracted organoid boundaries at 64x64 pixels were resampled to the desired number of points using periodic spline interpolation (Scipy *splprep*) and transformed back to real image size coordinates for analysis.

#### Segmenting organoids from tracked bounding box image crops

We trained a small binary segmentation attention-UNet^76^ neural network model of 4 layers to segment the predominant organoid in image crops. The model has 8 features in layer 1 with the number of features doubling in the next layer as the image size is downsized by a factor of 2 by average pooling. The model is trained on 542 unique cropped organoids from fluorescent confocal and label-free images resized to 64 x 64 pixels and random image augmentation. To be robust to real scenarios, the training set comprises crops of single isolated organoids and those where the organoid partially overlaps with other neighbouring organoids. Our trained segmentation model on a test set of 555 images obtained a test accuracy measured by IoU of 0.87.

To reduce storage on hard disk, we extracted the contour of the infilled binary segmentation using marching squares (scikit-image, skimage.measure.find_contours). To reduce computational time, the boundaries are computed from the binary segmentation after resizing input images to 64x64 pixels. Boundaries were then resampled to the desired number of points (here, 200 points, which offers a good compromise between number of points and reconstruction error) using periodic spline interpolation (Scipy *splprep*). The coordinates of the boundary is lastly transformed to correspond to the actual input image size for SPOT analysis.

#### Postprocessing organoid segmentation boundaries for analysis

Organoid segmentations represented by boundaries were post-processed to remove potential duplicate detections from bleed-through across channels in dense, overlapping organoid areas. This was done on a tracklet basis, jointly considering all tracklets from all image channels. For each pair of tracklets, the mean intersection over union (IoU) overlap of the organoid bounding boxes were computed over the time window when both organoids were present. Tracklets were deemed to overlap the same tracked organoid if the computed mean IoU score was greater than a user specified threshold (set at IoU=0.25 in this study). Overlapped pairs of tracklets were grouped into unique cliques. For each clique, only the longest tracklet with the highest consistency score (a composite score accounting for objectness, number of timepoints predicted by YOLOv3 vs imputed, and temporal smoothness of contours (assumes gradual, incremental shape changes)) was retained, and all other tracklets were removed. Next, individual organoids in tracklets lying on the border of the field-of-view and only partially captured (defined by measuring the percentage of border points that are located from the image border by a number of pixels at least greater than a pre-specified cutoff distance) were removed from the analysis. Tracklets were subsequently temporally filtered using central moving averaging. This precautionary step serves as imputation to minimize the effect of cutout segmentation error between consecutive frames, and improve the accuracy of motion feature computation. In a final step, we check each tracklet, pre-terminating and creating new tracklets to ensure each tracklet satisfies a minimum frame-to-frame IoU (computed using the boundaries’ bounding boxes) score. On top of using bipartite matching, this step further minimises the chance our tracker ‘identity switches’ within a tracklet to track a nearby, but different organoid.

**Stage 3: Computation of shape, appearance, and motion (SAM) features** and **Stage 4: Temporal analysis of SAM features** were performed as described in our companion methods-focussed manuscript (*Reviewers are requested to contact the Editor for access*). For Stage 4, step ii, the following default parameters were used for 2D UMAP projection of organoids: n_neighbors=100, random_state=0, spread=0.5, min_dist=0.5, metric=’euclidean. Depending on the total number of organoid instances relative to phenotypic heterogeneity, n_neighbors was adjusted to obtain a landscape that ‘spread’ the phenotypes. For example, we use reduced n_neighbors=35 with simulated organoids, to better display the 7 conditions in simulated dataset A.

### Simulating organoids with prescribed 3D shape, appearance and motion dynamics

We generated simulated organoid datasets with full control of the 3D phenotypic heterogeneity within and between treatment conditions. By design, this dataset has ground-truth for objectively assessing SPOT analysis performance in high-content organoid screens. Custom shape, appearance and motion parametrization of each organoid within a condition establishes that all ground truth phenotype states are distinct, while the uniqueness of each individual organoid is maintained by sampling its parameters from Gaussian distributions. For each condition, all parameter values are made available in the associated metadata files at DOI: 10.5281/zenodo.14265519 and in Suppl. Table 5.

To simulate shape evolution, we generated an initial 3D organoid shape as a surface mesh, and deformed the mesh in successive timepoints. To generate the initial shape we start with a spherical icosphere mesh; sample surface coordinates to locate organoid branches; and then creating the organoid branches by deforming the local icosphere mesh vertices around to each coordinate with the same effect as placing a 3D ellipsoid with major axis oriented perpendicular to the surface. Organoid growth is simulated by isotropic scaling of mesh vertices from the organoid centroid. Then, branch growth is simulated by displacing branch-associated vertices further from the organoid centroid, with the same effect as stretching along its major axis, a 3D ellipsoid with major axis oriented perpendicular to the surface. To simulate a gradual shape evolution, we create a 3D final organoid shape similarly retaining the branching position and number of branches. We then linearly interpolate organoid and branch growth-associated parameters between initial and final shapes for intermediate timepoints.

To simulate appearance evolution, voxel-based intensity patterns were generated and intersected with the inner volume of the evolving 3D organoid meshes at each timepoint. 2D projections of the composite shape and appearance 3D simulations were produced by sampling the 3D organoid at multiple 2D planes and generating a single 2D image in an analogous way to extended depth-of-focus algorithms. The synthetic organoid boundary sampled in each 2D plane was smoothed by fitting a Bezier curve, with mesh-plane intersections serving as control points.

To simulate motion we translated and rotated organoids. For translation, at each timepoint, we sample two random vectors with their magnitude and direction (represented by a unit vector) independently sampled from Gaussian distributions, representing ‘migration’ and ‘jitter’, to displace the organoid centroid. The ‘migration’ vector represents longer-time persistent directional movement. At each timepoint, we determine with probability 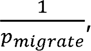, where *p_migrate_* can be interpreted as number of frames motion persists unidirectionally, whether to change its direction by sampling a new unit Gaussian vector. The ‘jitter’ vector represents stochastic movement between timepoints with no net directionality. A new direction unit vector is sampled at every timepoint. For rotation, at each timepoint we rotate the organoid by 3^0^ about a rotation axis passing through the organoid centroid and represented by a unit Gaussian random vector. The rotation axis is modelled with longer-time directional persistence as with the migration vector. At each timepoint, we determine with probability, 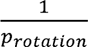 where *^p^_rotation_* can be interpreted as the number of frames rotation persists in the same direction, whether to change rotation axes by sampling a new unit Gaussian vector.

To simulate a gradual motion evolution we linearly interpolated magnitude parameters of initial and final states for intermediate timepoints. The same probability of migration and rotation changing direction between timepoints were used for all simulated conditions.

We used the described SAM simulation strategy to generate two datasets of organoids for validating SPOT analysis where each organoid is unique with a different shape, appearance and motion parameterization and categorized by a distinct SAM phenotype.

#### Simulated Dataset A for assessing the impact of imaging artefacts and processing errors

Dataset A comprises 7 conditions, 48 organoids per condition, and 30 frames per organoid. Each of the conditions share a common starting state with simulation parameter values sampled from Gaussian distributions (see above), before gradually evolving over the 30-frame time course into their uniquely-parameterised, and condition-specific end SAM state. However, ‘appearance-high’ and ‘appearance-low’ were simulated with a higher-density appearance pattern to exacerbate final state differences. We define one condition-dependent end state as the ‘base organoid’, with average parameter values of shape, appearance and motion. Using this base organoid parameterisation as template, and modifying the parameters influencing each of shape, appearance, and motion in isolation, we created the other 6 further ‘high’ or ‘low’ extremal SAM phenotypes. Simulated dataset A is made available with individual 3D organoid meshes and associated 2D projection videos at DOI: 10.5281/zenodo.14265519.

#### Perturbing Dataset A to mimic imaging artefacts and errors in segmentation and tracking

We perturbed Dataset A to investigate the impact of three primary categories of errors on SPOT analysis (Suppl. Fig. 2c): 1. imaging-based artefacts including: (i) blur, (ii) overexposure, (iii) underexposure, and (iv) foreign-object artefacts; 2. segmentation errors including: (v) overlap between organoids, (vi) partial segmentation of an organoid (cutout), and (vii) multiple organoids segmented as one (union); and 3. tracking error, namely (viii) track fragmentation as a result of incomplete tracking due to occlusion.

These perturbations were simulated with across nine increasing severity levels as follows:

##### (i) Blurring

2D projection images of individual organoids were Gaussian blurred at increasing sigma (level 1: σ = 2, level 9: σ = 14) using the GaussianBlur function from the OpenCV library.

##### (ii) Overexposure

8-bit 2D projection images of individual organoids were overexposed by increasing the image brightness, *I* by a fixed amount, γ.

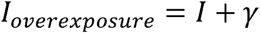

Any pixels with resulting brightness greater than 255 is clipped to 255, the maximum intensity of 8-bit images and represents camera saturation. The added amount increases linearly with severity level (level 1: γ = 20, level 9: γ = 180)

##### (iii) Underexposure

8-bit 2D projection images of individual organoids were underexposed by decreasing the image brightness, *I* by a fixed amount, γ.

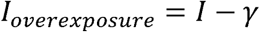

Any pixels with resulting brightness lower than 0 is clipped to 0, the minimum intensity of 8-bit images. The decreased amount increases linearly with severity level (level 1:γ = 10, level 9: γ = 90).

##### (iv) Foreign-object artefacts

We simulated two types of common object artefacts that may be imaged but unlike organoids do not temporally evolve. The first artefact is small-sized pixel-scale objects, such as fine dust. We simulated these as ‘salt’ noise in the 8-bit 2D projection images, where we replace the value of a randomly selected pixel with 255, the maximum brightness. We increase the fraction of image pixels to replace with noise using the random_noise function from scikit-image linearly for increasing severity (level 1: amount = 1/6000, level 9: amount = 9/6000). The upper amount was qualitatively determined to represent high noise at a realistic density. The second artefact is larger-sized fibre-like debris such as hair or clothes fibres. We simulate these with Bezier curves and placed them in random spatial locations in the image. Bezier curve thickness and length were chosen to resemble clothes fibres at organoid-scale. Increasing severity was simulated by increasing the number of fibres proportionately (level 1: *n* = 15, level 9: *n* = 135 fibres). The simulated dust and fibre artefacts are randomly generated for an organoid and remains identical in every timepoint.

##### (v) Overlapping organoids

We simulate the effect of correctly segmenting individual organoid boundaries but with internal appearance differences due to intensity interference from overlapping with a neighbour organoid. Intensities in the overlapped area were simulated by pairing organoids from the same condition and timepoint, aligning them horizontally with a degree of overlap, then summing pixel intensity values and clipping the result to be within the 8-bit range. Severity level corresponds to increased area of overlap; we fix a baseline level of overlap at 40 pixels, beyond which additional overlap is added as a percentage of the remaining non-overlapping area (level 1: 0%, level 9: 100%).

##### (f) Cutout

When organoids overlap, the segmentation may erroneously segment only the non-overlapped area, resulting in a smaller ‘cutout’ of the actual shape. We simulate this by pairing organoids from the same condition and timepoint, aligning them horizontally with a degree of overlap, then subtracting the overlapping area from each of the organoids. Severity level corresponds to increased area of overlap; we fix a baseline level of overlap at 40 pixels, beyond which additional overlap is added as a percentage of the remaining non-overlapping area (level 1: 0%, level 9: 100% overlap).

##### (g) Union

When organoids overlap, the segmentation may erroneously regard two organoids as one. We simulate this effect by pairing organoids from the same condition and timepoint, aligning them horizontally with a degree of overlap, and taking their union to be a single instance. Severity level corresponds to increased area of overlap; we fix a baseline level of overlap at 40 pixels, beyond which additional overlap is added as a percentage of the remaining non-overlapping area (level 1: 0%, level 9: 100% overlap).

##### (h) Fragmented tracks

We define the perfect baseline ground truth as 100% coverage whereby each individual organoid is 100% tracked for all 30 frames. To generate fragmented tracks, we fixed coverage at 50% i.e. a total of 15 frames is tracked per organoid, and at each increase severity level we increase the number of synthetic breaks by one. For example at level 1, the organoid has one tracklet, covering 50% of frames, but at level 2 has two tracklets, each covering ≈25% of frames, such that the total is still 50%, at level 3 three tracklets, etc.

#### Measuring the impact of perturbing Dataset A to SPOT analysis

Taking the unperturbed Dataset A as severity level 0, for each perturbation type (a)-(h) above, we applied SPOT, preprocessing all SAM phenomes together to derive a joint UMAP phenomic landscape pooling all organoids across levels 0-9. We then assessed the impact of each perturbation type on SPOT analysis by computing metrics of agreement and disagreement in the four primary SPOT outputs: (i) phenotype clustering agreement; (ii) stacked barplot disagreement, (iii) phenotype trajectory disagreement, and (iv) HMM transition matrix disagreement, between each severity level to the unperturbed dataset A (ground-truth).

##### (i) Phenotype clustering agreement

K-means with automatic cluster number selection using the elbow method was applied to partition the joint UMAP into phenotype clusters. Two cluster metrics: adjusted random index (ARI) and adjusted mutual information (AMI) were used to assess the agreement in the phenotype cluster assignment at each severity level compared to the unperturbed. ARI and AMI scores agreement on scale from 0 (perfect disagreement) to 1 (perfect agreement).

##### (ii) Stacked barplot disagreement

We model the stacked barplot as a 2D probability distribution, and measure the disagreement to ground truth using the Wasserstein optimal transport metric. Specifically we compute the sliced Wasserstein metric (max_sliced_wasserstein_distance implemented in the Python POT library) using uniform weights and the matrix representations of the ground truth and perturbation stacked barplot data as the samples, stabilising the estimate by setting n_projections = 50. This metric enables us to score disagreement based on intuitive ‘pattern similarity’ of the cluster frequency. Thus, the same cluster pattern but temporally lagged will have lower disagreement than a cluster pattern with expansion of one cluster and contraction of another cluster.

##### (iii) Phenotype trajectory disagreement

We measure whether the similarity between phenotype trajectories of each condition in dataset A given by their computed trajectory affinity matrix with the joint UMAP is preserved at each severity level. The disagreement at severity level i, is the mean of the absolute elementwise difference between the trajectory affinity matrix of the unperturbed dataset and that at level i.

##### (iv) HMM transition matrix disagreement

We measure the mean of the absolute elementwise difference between the HMM cluster transition matrix for the unperturbed dataset and that at each severity level 1 to 9.

These metrics were assessed and plotted in Suppl. Fig 3c for all perturbation types. Tracking error represented by fragmented tracks used the same phenomic landscape constructed by the unperturbed dataset A. Phenotype cluster agreement was therefore not assessed. To compute the stacked barplot, trajectory and HMM transition disagreement metrics, the subset of organoid UMAP coordinates corresponding to the fragmentation were used at each level.

#### Simulated Dataset B for assessing the impact of motion in the z-direction

Dataset B was generated to assess whether z-movement is detectable in SPOT analysis of 2D projection videos. Dataset B simulates 3 conditions, 96 organoids per condition, and 30 frames per organoid. The individual organoids were used to compose 6 3D scenes of 12 organoids in the field-of-view, a total 96 organoids per condition. 3D scenes were projected as 2D videos and analysed by SPOT (Fig. 2). All organoids were simulated with random directional movement (i.e. ‘jitter’) of the same speed and random rotation between consecutive frames, as described above without migration. In the first condition, organoids can move unrestricted in any 3D XYZ direction (denoted XYZ movement). The second condition replicates the first condition using a modifed construction, with the same condition-specific parameters but individual organoids are unique with different individually sampled parameters for shape, and appearance.,. Each organoid moves freely in 3D but its z-component movement is not recorded. Its position is updated with the full XYZ, and therefore is indistinguishable from the first condition under SPOT analysis. We denote the first and second condition as XYZ replicate 1 and 2 respectively. The third condition constrains organoids to only move in the 2D XY direction (denoted XY). Otherwise, organoid shape and appearance were parameterised identically. The three conditions were used to assess whether SPOT could cluster XY movement distinctly from the two replicates of XYZ movement, which represent stochastic biological variation.

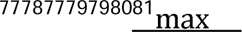

### scRNA-seq analysis

#### scRNA-seq pre-processing and quality control filtering

Pre-processing of scRNA-seq data was conducted using Cell Ranger (v3.1.0). Briefly, FASTQ files were demultiplexed from raw BCL format using cellranger *mkfastq*. A custom Cell Ranger reference was then prepared (cellranger *mkref*) using a concatenation of the default Cell Ranger mouse genome sequence and annotation files (mm10-2020-A: GENCODE vM23/Ensembl 98) and custom sequence and annotation files for the fluorescent markers used in this study. Alignment to this custom reference, identification of true cells, and tabulation of UMI counts for each sample was performed using cellranger *count*. The UMI counts for each sample were then combined into a single matrix using cellranger *aggr* without normalization.

Initial downstream processing was conducted using Seurat (v3.9.9.9038). As an initial filter for low-quality cells, cellular barcodes with less than 2000 genes detected or more than 10% mitochondrial reads were removed. Normalization, variance-stabilizing transformation (VST), and scaling were conducted on the remaining cells using Seurat’s implementation of sctransform (v0.3.2.9002), and the scaled VST counts for the 3000 most variable features were extracted from the Seurat object (scale.data slot).

#### scRNA-seq selection of variable genes and clustering

The scaled variance-stabilised counts of the 3000 most variable genes were used to cluster single cells using the graph-based Louvain algorithm (scanpy *Louvain* function) applied to the *k* nearest neighbor adjacency matrix (*scanpy.pp.neighbors*) using 20 principal components and *k*=15 for murine colorectal organoid scRNA-seq.

#### scRNA-seq cluster connectivity and dimensionality reduction

To identify relationships between clusters (‘cluster connectivity’), Partition-based Graph Abstraction (PAGA) with connectivity model v1.2 was used, with Louvain clustering at resolution=1. Based on the PAGA connectivity, murine mutant colon organoid scRNA-seq data were visualized in 2D using the ForceAtlas2, ‘fa’ layout positions through *scanpy.pl.draw_graph* (Fig. 6). For patient-derived duodenum scRNA-seq, 2D-UMAP coordinates were computed using the ForceAtlas2 PAGA as initial embedding positions (Fig. 6). Results of the ForceAtlas2 and UMAP gave the same interpretative results. We chose UMAP for figure visualization as it produces a better spread of points.

#### Combined visualization of multiple genes

Combined expression of signatures was plotted on the PAGA graph or UMAP coordinates as the mean z-score expression of the selected marker genes. The zscore of each gene was computed after log transformation, ln(raw count + 1) using scanpy.

## Data and code availability

Sequencing datasets used in this study are available from GEO: mutant mouse colon organoid scRNA-seq, accession number GSE218339, mouse small intestinal bulk RNA-seq, accession number GSE218337 and human small intestinal scRNA-seq, accession number pending.

Simulated organoid datasets A and B used in this study are available at DOI: 10.5281/zenodo.14265519.

The developed SPOT Python library used to detect, track and segment individual organoids and to compute and analyse the SAM phenome in this paper is available at GitHub, https://github.com/fyz11/SPOT.

Any additional information required to reanalyse the data reported in this paper is available from the authors upon suitable request.

## SUPPLEMENTARY FIGURES

**Supplementary Figure 1a.**
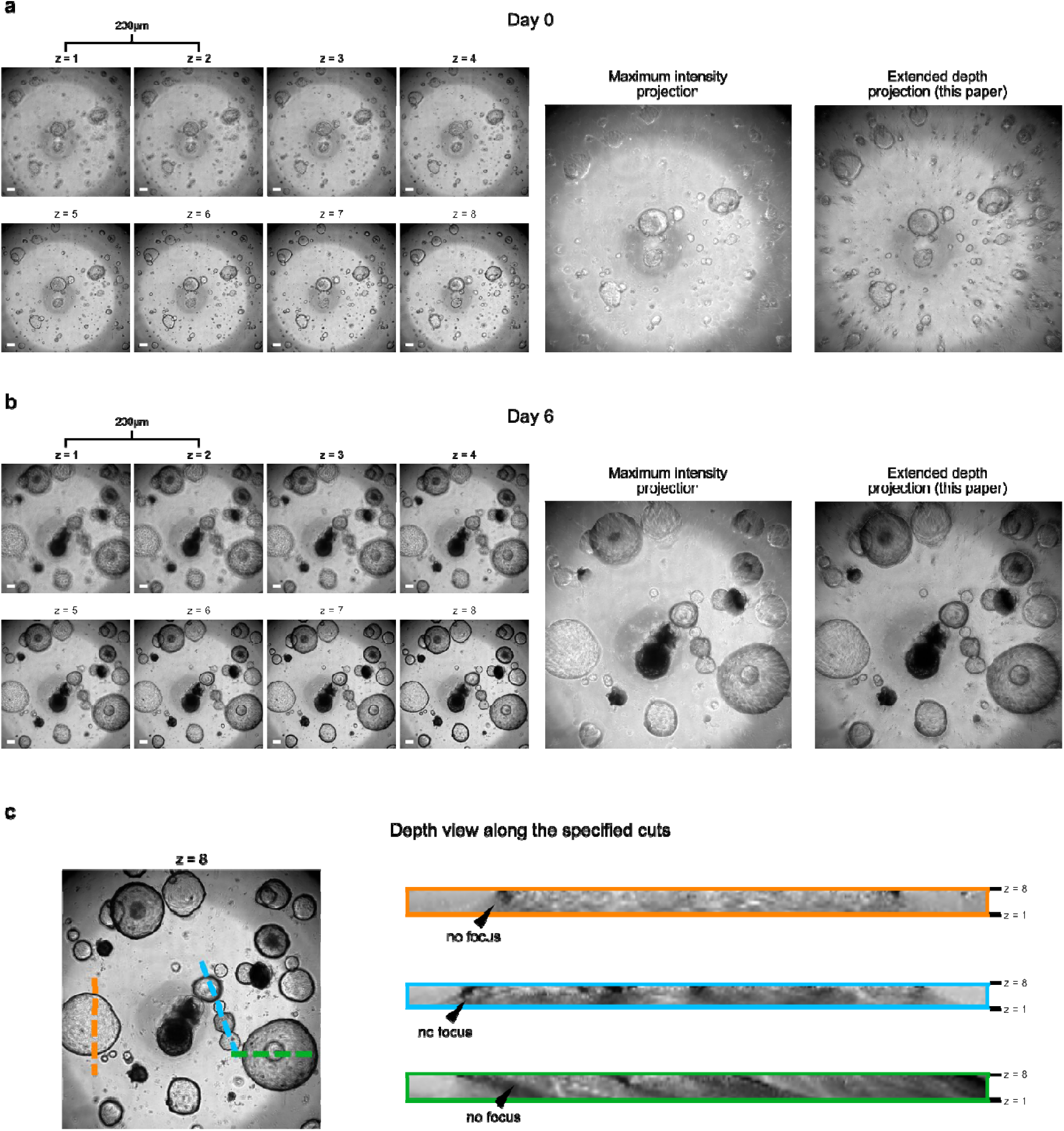
Brightfield phase contrast imaging has limited depth resolution and extended focus projection of a z-stack to a 2D image. **a)** Individual z-slices of the acquired 8-slice z-stack at day 0 (left), its 2D maximum intensity projection (middle), and 2D extended focus projection used in this paper (right). Each z-slice is separated by a depth of 200 µm. **b)** Same as a) of the same organoids acquired at day 6. **c)** Reslice of the z-stack image along the indicated color line indicated in the corresponding z=8 xy-slice on the left. Black triangle mark the edge of the organoid. All scale bars: 250 µm.

**Supplementary Figure 1b.**
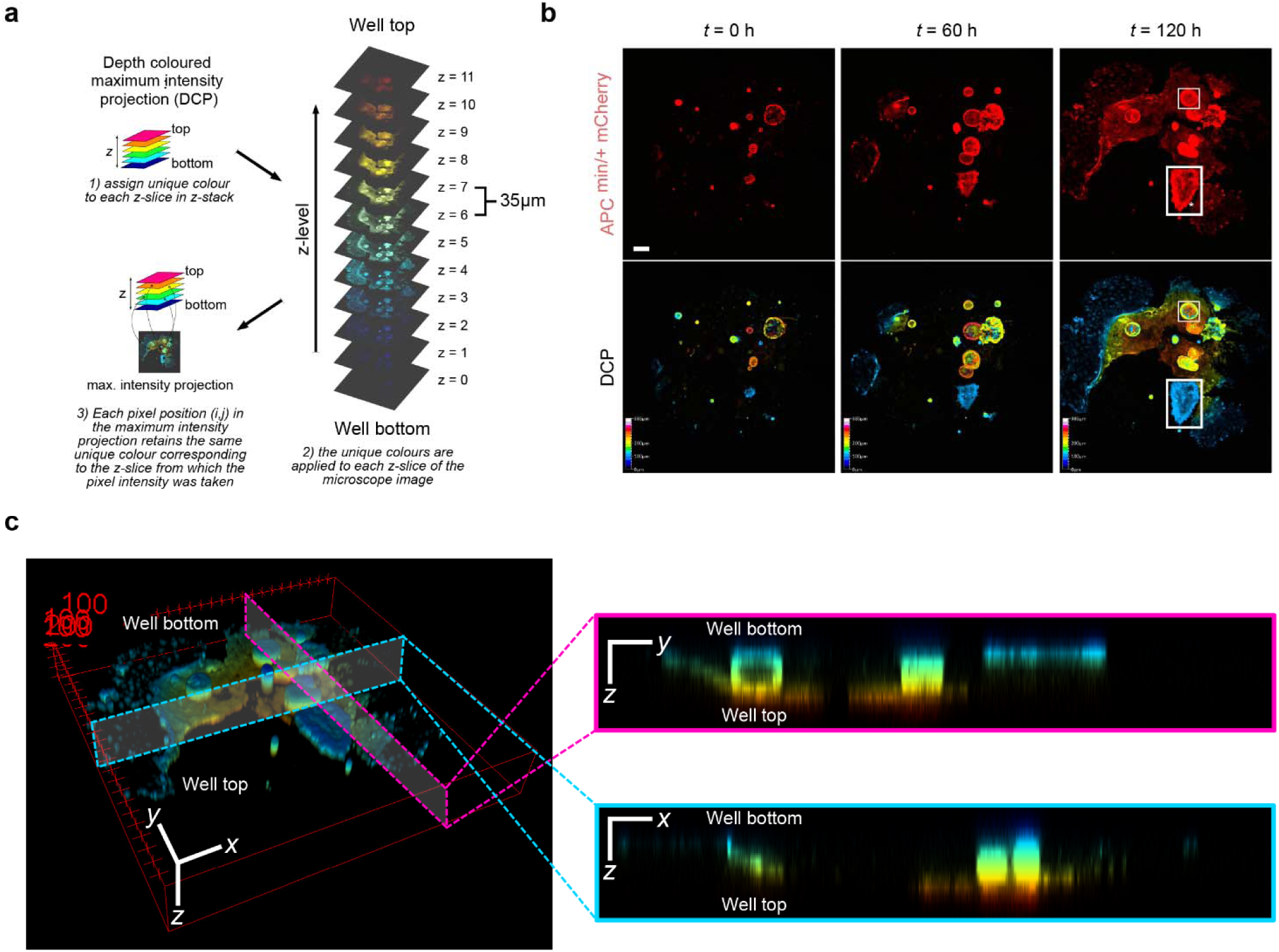
Acquired confocal z-stacks have depth-limited resolution. **a)** Schematic of computing a depth colored z-stack and its maximum intensity projection 2D image (DCP). **b)** DCP of initial, intermediate and final timepoint of APC mutant colon organoid undergoing large morphological transformation illustrating the effective capture of in-focussed organoids from different z-slices using 2D projection, here by maximum intensity. Scale bar: 200 μm. **c)** 3D render of the z-stack of the final timepoint 120 of b) (left) and its reslice in the two orthogonal colored planes (right). The coarse depth resolution offers limited 3D shape, and appearance information, whilst the available space in the ECM dome constrains organoid movement to be largely planar in xy.

**Supplementary Figure 2a.**
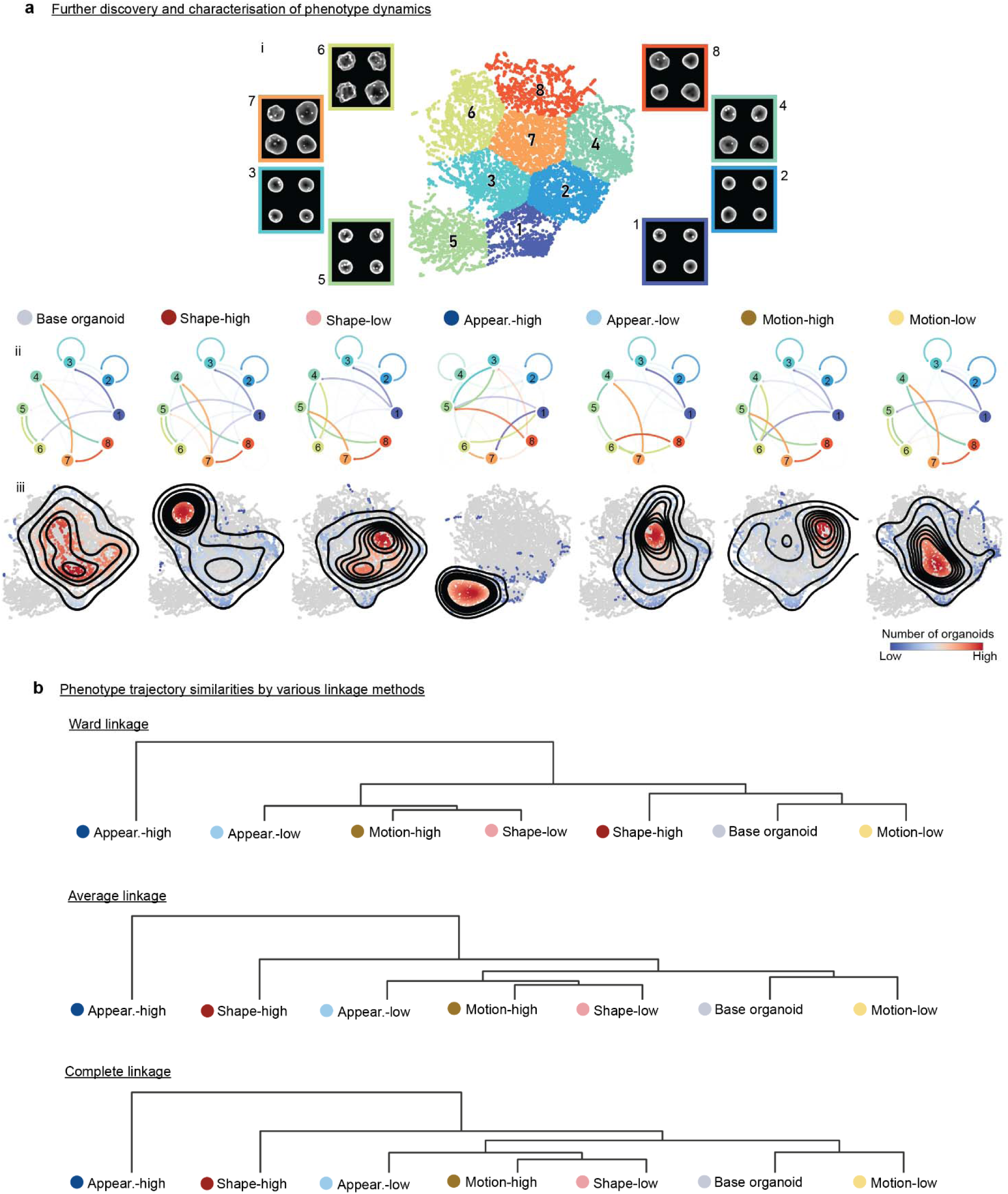
SPOT Phenotype cluster transition and density maps of the simulated organoid dynamics dataset A. **a)** (i) UMAP phenomic landscape of all organoids and conditions, same as Fig. 2d. (ii) Hidden Markov Model (HMM) inferred transition probabilities display the likelihood of a transition occurring at the subsequent timepoint given a phenotype cluster label at the current timepoint, where arrow color represents the cluster of origin and its transparency represents the probability of the transition. (iii) UMAP point density heatmaps displaying how high and low perturbations of organoid behaviour in shape, appearance and motion map within the phenomic landscape. **b)** Hierarchical clustering of phenotype trajectories per condition, using Ward, average and complete linkage, top-to-bottom.

**Supplementary Fig. 2b.**
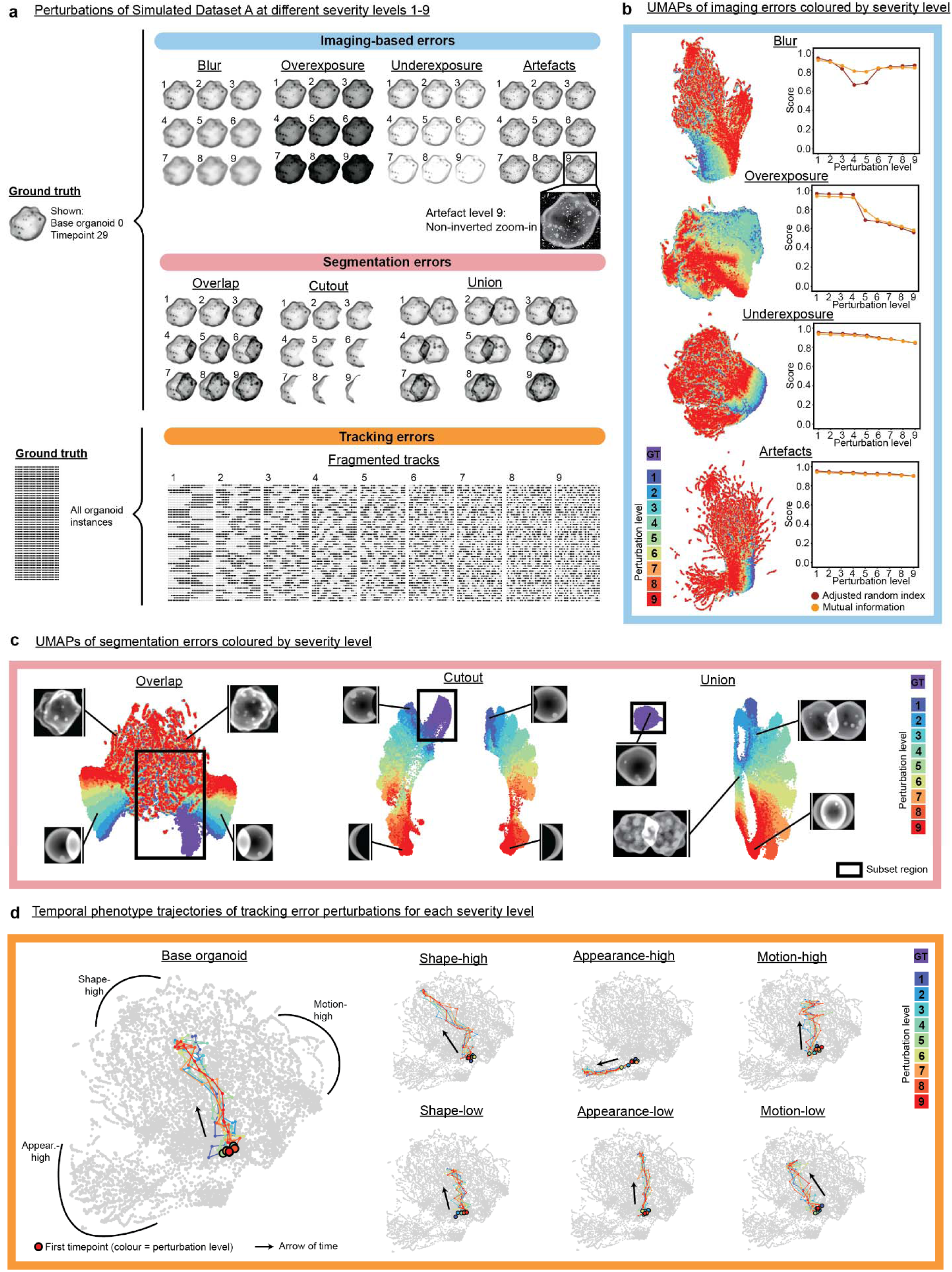
Robustness of SPOT analysis to imaging artefact, segmentation and tracking errors. **a)** Depiction of simulated perturbations grouped into imaging, segmentation and tracking error categories across 9 severity levels (Methods). Imaging and segmentation errors are illustrated using frame 29 of ‘base organoid’ 1 as a reference image. Fragmented tracks are represented by consecutive black squares in the horizontal direction. The ground truth dataset has 100% complete organoid tracks, 100% black squares. Each increasing severity level includes an additional track breaking but overall temporal coverage (total black squares) is fixed at 50% (Methods). Organoid exemplar images are depicted with inverted intensity such that underexposure appears bright and overexposure appears dark. A non-inverted artefact level 9 exemplar image is shown for clarity. **b)** For each of blur, overexposure, underexposure and artefact, a single UMAP plot (n=97440) was generated to jointly analyze ground-truth and its perturbations, colored by perturbation level. After K-means clustering as in Fig.2c, adjusted random index and mutual information was computed to assess phenotype clustering agreement between the ground truth and each perturbation level (Methods). **c)** UMAP for each segmentation perturbation are colored by perturbation severity level, showing that the ground truth can easily be isolated (black crop boxes), thereby enabling exclusion of a both severe and subtle segmentation error manifestations. Reference images are depicted alongside each UMAP, with black lines locating each instance on the plot. **d)** Temporal phenotype trajectories of each condition, for ground-truth and all track fragmentation levels.

**Supplementary Fig. 2c.**
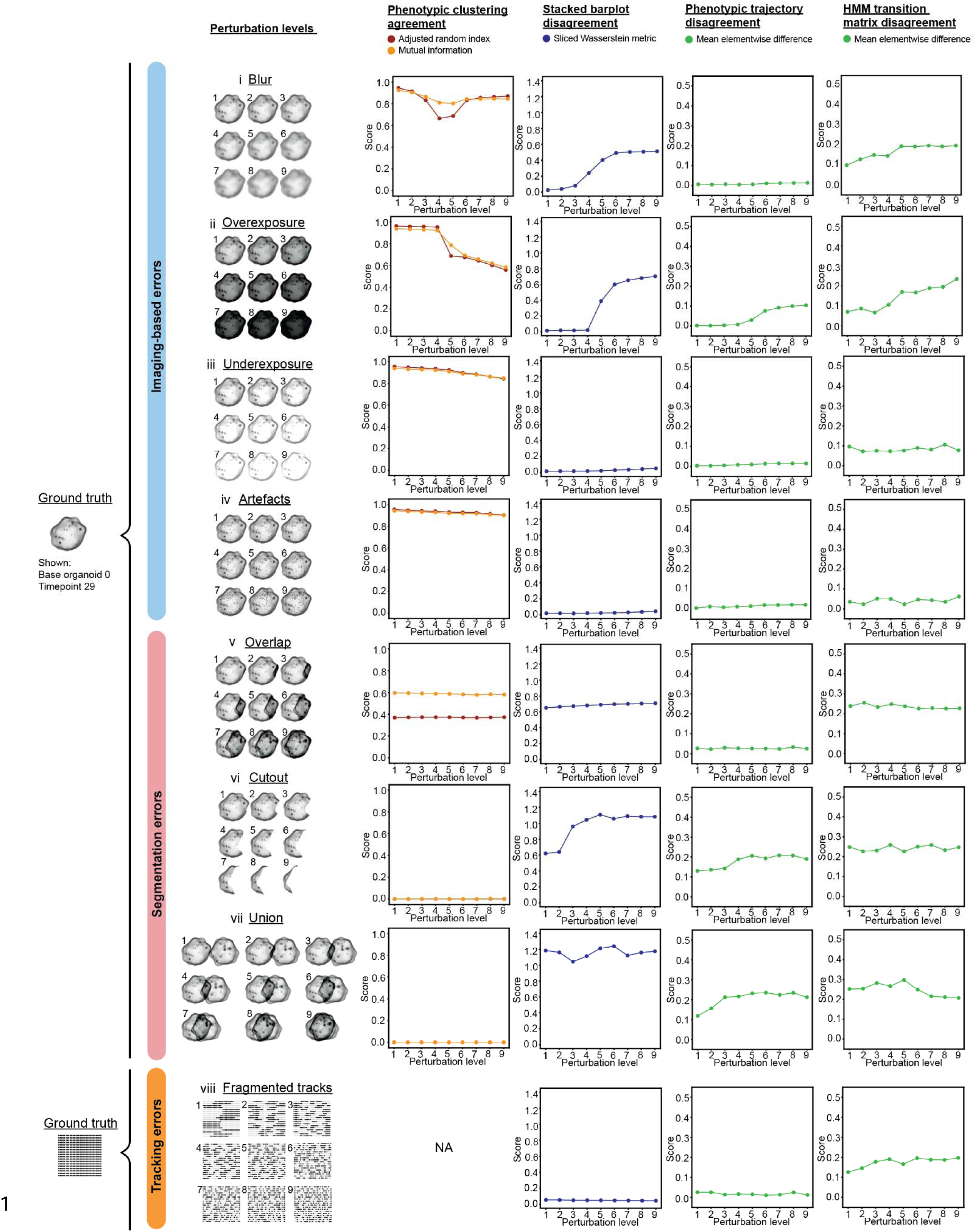
Quantitative assessment of SPOT analysis robustness to imaging artefact, segmentation and tracking errors. Schematic of increasing severity level of perturbation, and i) phenotype clustering agreement, ii) stacked barplot, iii) phenotype trajectory and iv) HMM cluster transition matrix disagreement metrics of ground-truth with each perturbation level (left to right). For agreement metrics, the higher the value, the greater the concordance with ground-truth. For disagreement metrics, the lower the value, the greater the concordance with ground-truth. Multiple independent evaluation measures were used to evaluate phenotype clustering agreement and stacked barplot disagreement. Mean elementwise difference was used to compute phenotype trajectory and HMM cluster transition matrix disagreements, (Methods).

**Supplementary Fig. 2d.**
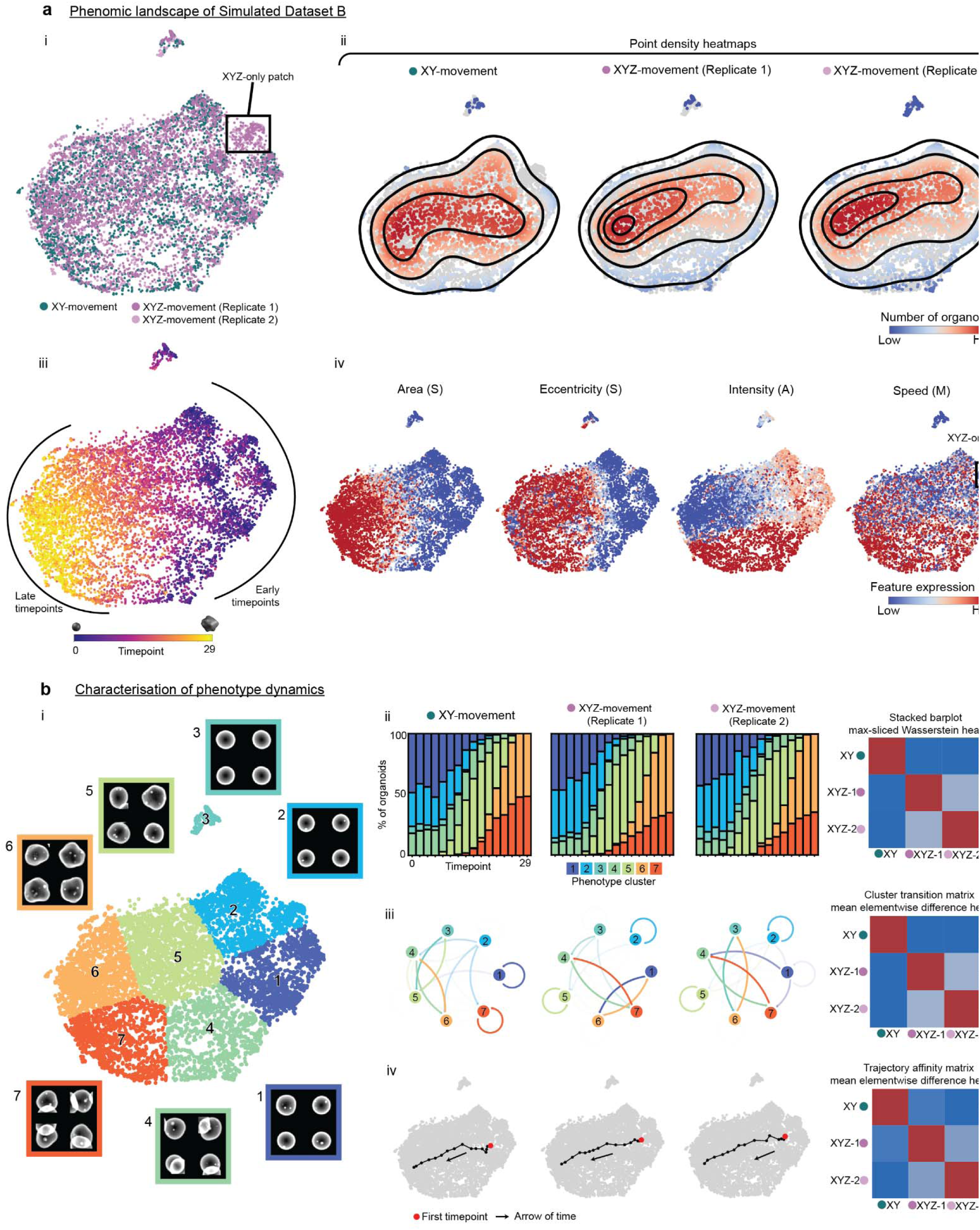
SPOT analysis of 2D projections discriminates z-component motion. **a)** Phenomic landscape of Simulated Dataset B, colored by (i) ground truth conditions, (ii) density heatmaps, (ii) timepoint in synthetic organoid evolution and (iv) selected individual features (area, eccentricity, entropy and mean speed). The region identified by a black box in (i) and (iv) denotes a UMAP region with no instances from the XY-movement condition.**b),** (i) Partitioning of phenomic landscape into 7 clusters using K-means clustering, along with representative images for each cluster. (ii) Stacked barplots show temporal changes in phenotype cluster distribution for each simulated organoid 3D movement condition and pairwise computation of the max-sliced Wasserstein distance allow average-linkage hierarchical clustering to quantify stacked barplot similarity between conditions. (iii) Hidden Markov Model (HMM) inferred transition probability (Methods) showing cluster transition likelihood for each condition, where arrow color represents the cluster of origin and its transparency represents the probability of the transition. The mean elementwise difference between pairs of HMM transition matrices is computed for all condition permutation, and hierarchical clustering is performed using average linkage. (iv) Temporal phenotype trajectories show the progression of each simulated organoid 3D movement condition across the phenomic landscape over time. Red point indicates the origin of the trajectory at t=0. Trajectories are clustered via average-linkage hierarchical.

**Supplementary Figure 3a.**
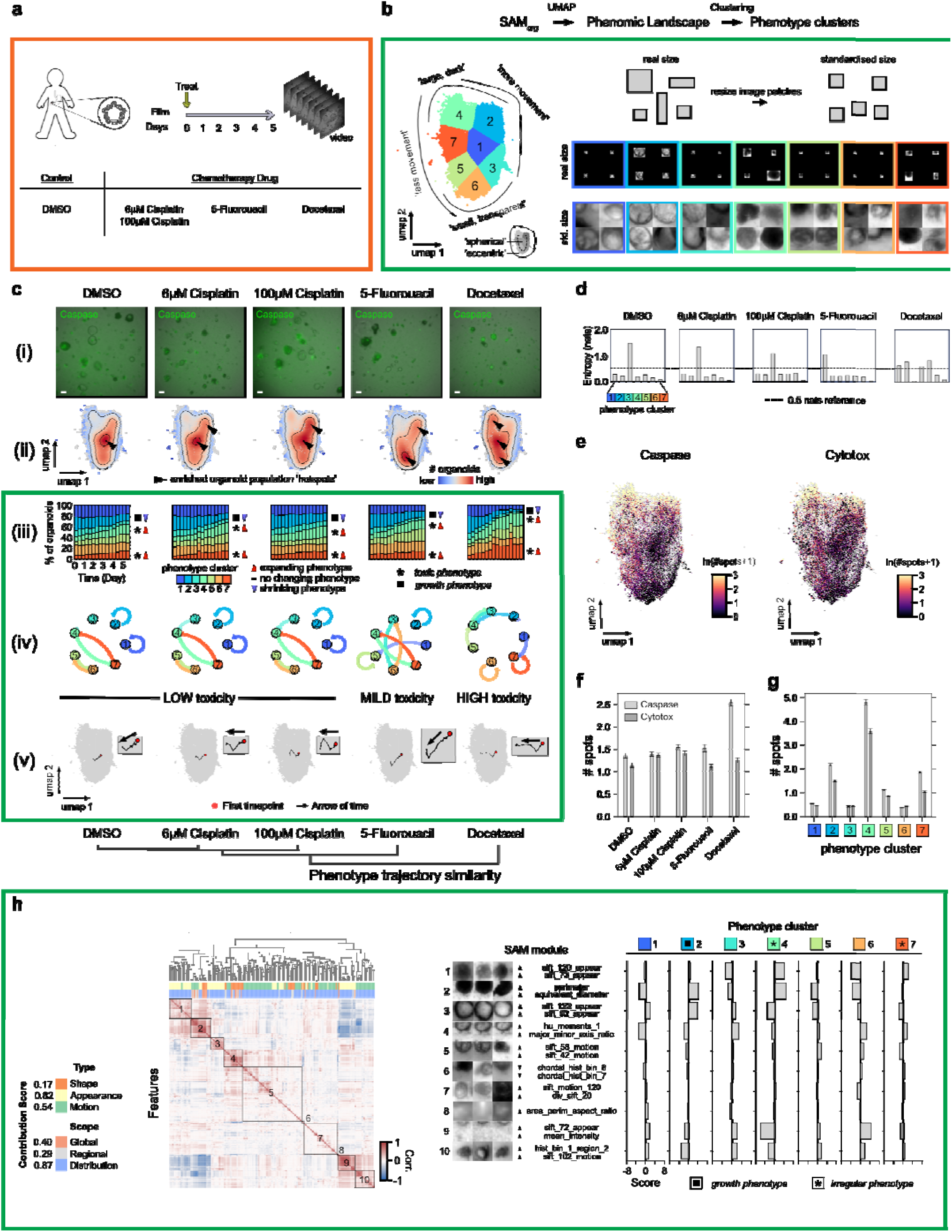
SPOT phenotype clusters capture different SAM characteristics of cell death after drug treatment. **a)** Patient-derived D2 duodenum organoids labeled with Hoescht and additional Caspase 3/7 Green or Cytotox Green were grown from endoscopy biopsies, in a 96-well plate with the specified drug treatment and concentration, then filmed every 4 hours for 5 days. Drugs without concentration specified used the corresponding maximum concentration in Fig. 3. (Methods). **b)** UMAP phenomic landscape, phenotype clusters and representative cluster images as in Fig. 3 using the filtered SAM phenomes computed from only the brightfield channel from all videos across all drugs and concentrations, n=70,757 points. **c)** Hierarchically clustered chemotherapy drugs according to SAM temporal phenotype trajectories using average linkage (bottom). (i) final frame combined with Caspase 3/7 Green channel of exemplar videos for each indicated drug treatment and drug concentration, scalebar: 250 µm; (ii) UMAP point density heatmaps, same UMAP coordinates as in (b); (iii) stacked barplots showing the temporal changes in phenotype cluster (as characterized in (c)) distribution for each drug, as computed from organoids at 8-hour time intervals; (iv) graph showing the Hidden Markov Model (HMM) inferred transition probability (Methods) of an organoid transitioning to another phenotype cluster in the next timepoint given its phenotype cluster label at the current timepoint. Arrows are colored by the source cluster. The more transparent the arrow, the smaller the probability of the transition. (v) UMAP SAM phenotype trajectories for a density threshold of mean+3 standard deviations (left) and zoom of the trajectory (right) for each drug. On each trajectory, each black point denotes 8.5-hour increments from the origin red point. A black arrow indicates the directionality of time. **d)** Shannon entropy of the HMM transition probability for each phenotype cluster, in each drug condition. **e)** UMAP with points coloured by its number of counted punctate Caspase 3/7 (left) and Cytotox (right) spots. **f)** Mean±s.e.m number of punctate Caspase 3/7 and Cytotox spots of organoids in each drug condition. **g)** Mean±s.e.m number of punctate Caspase 3/7 and Cytotox spots of organoids in each phenotype cluster. **h)** Contribution score of shape, appearance and motion, and global, regional and distributional features (left), automated hierarchical clustering to identify SAM feature modules (middle) and SAM module expression in each phenotype cluster (right) constructed as in Fig. 2f.

**Supplementary Figure 3b.**
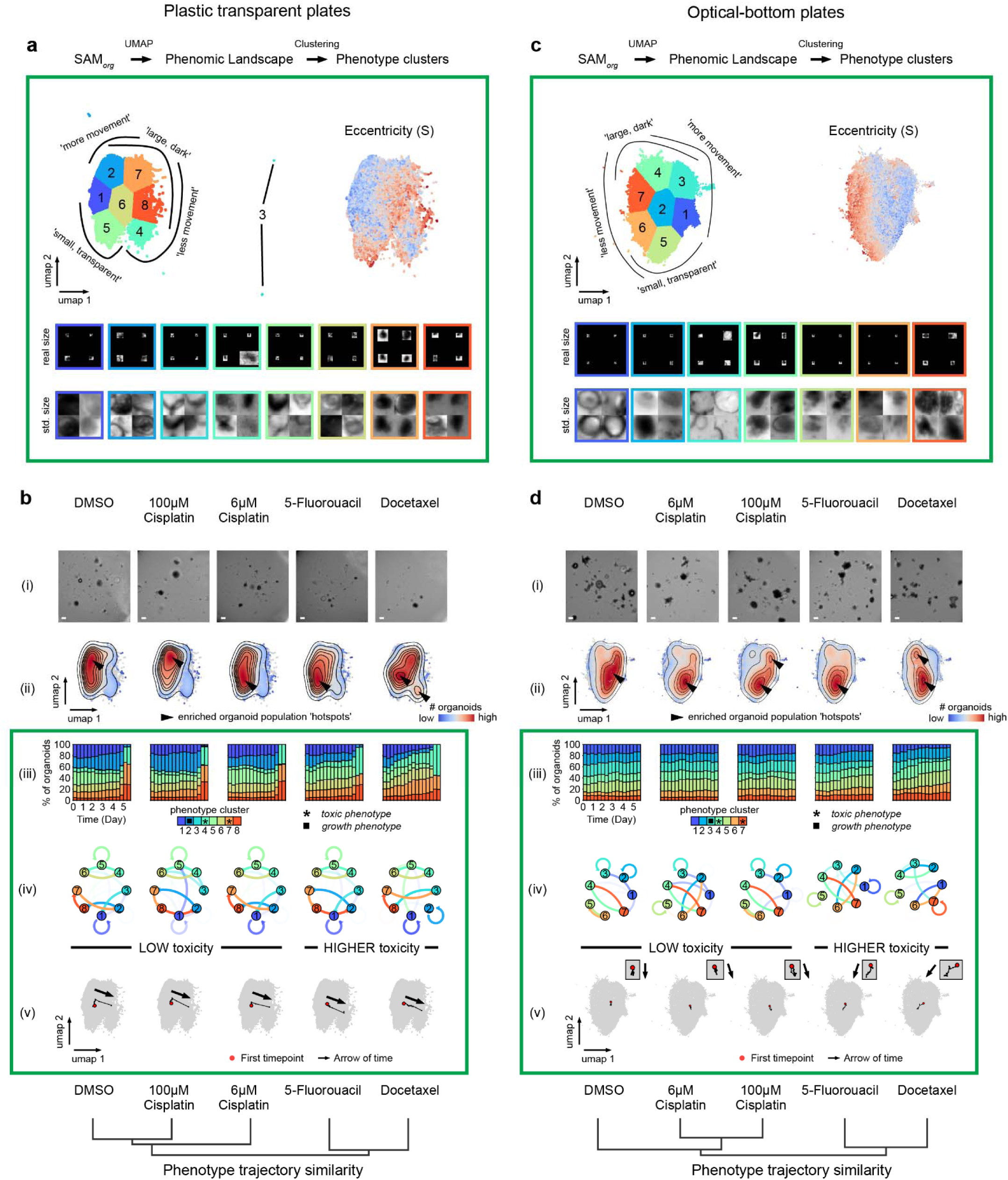
Independent SPOT analysis of organoid culture in plastic transparent and optical-bottom plates. **a)** UMAP phenomic landscape of organoids cultured in plastic transparent plates. Top row: landscape colored by phenotypic clusters, dark blue to orange and labeled by increasing mean shape eccentricity of each cluster (left). Same landscape colored by shape eccentricity (right). Bottom row: four representative images of organoids in each cluster arranged as a 2x2 grid, in real and standardized (std.) size. n=55,581 points. **b)** Same as a) for optical-bottom plates. n=53,078 points**. c)** Hierarchical clustering of drug conditions by phenotype trajectory similarity for plastic transparent plates. For each drug condition, (i) example final video frame, scalebar: 250µm; (ii) point density heatmap, (iii) stacked barplot of temporal phenotype cluster composition, (iv) HMM cluster transition probabilities, and (v) temporal phenotype trajectories. d) Same as c) for optical-bottom plates.

**Supplementary Figure 3c.**
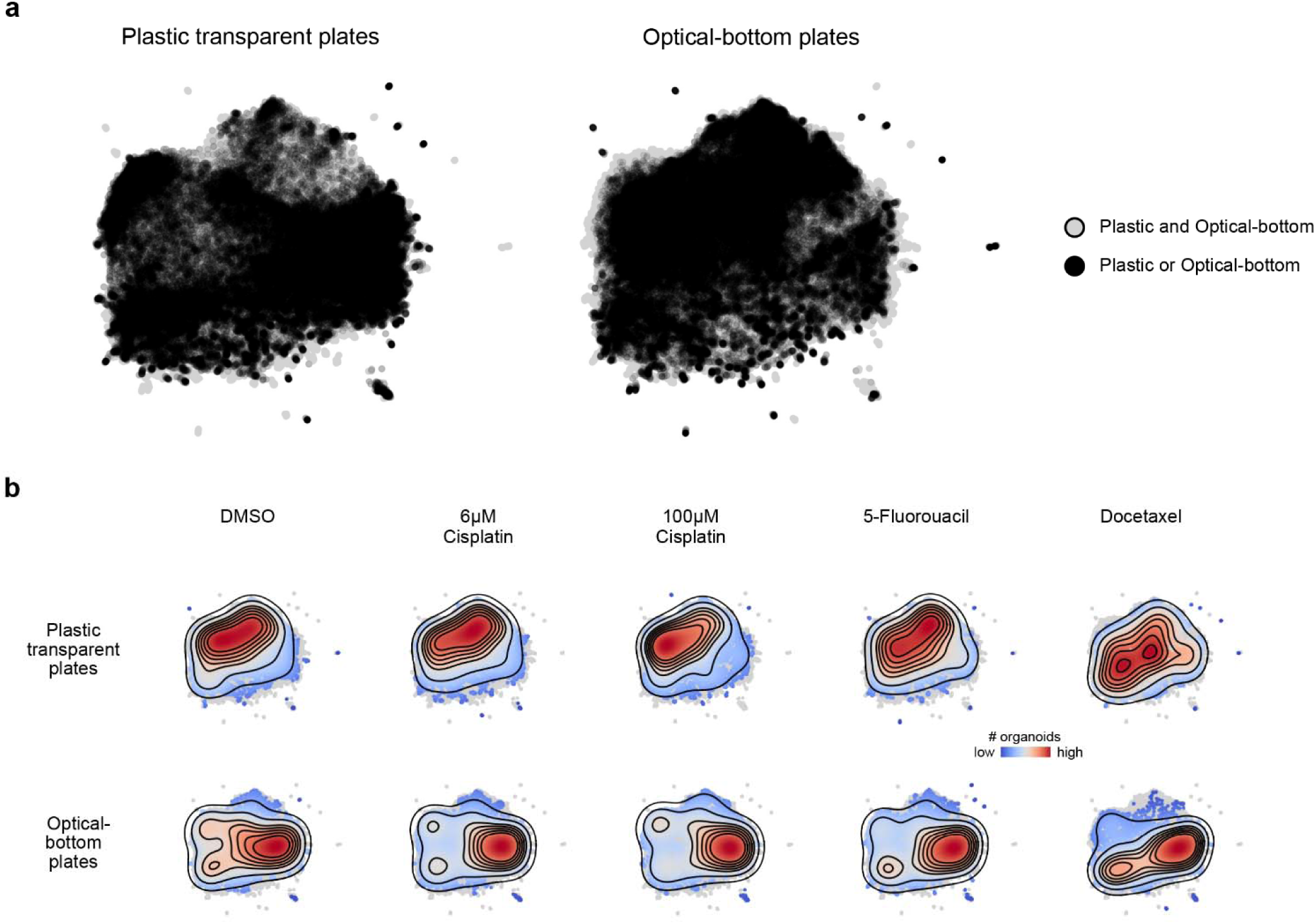
Joint UMAP analysis of chemotherapy treated patient-derived organoids cultured in plastic and glass-bottom plates. **a)** Joint UMAP phenomic landscape (light grey points), with organoids cultured in plastic transparent (left) and optical-bottom plates (right) highlighted with black points. **b)** Point density heatmaps showing how organoids cultured in the different plates and drug condition distribute in the UMAP phenomic landscape.

**Supplementary Figure 3d.**
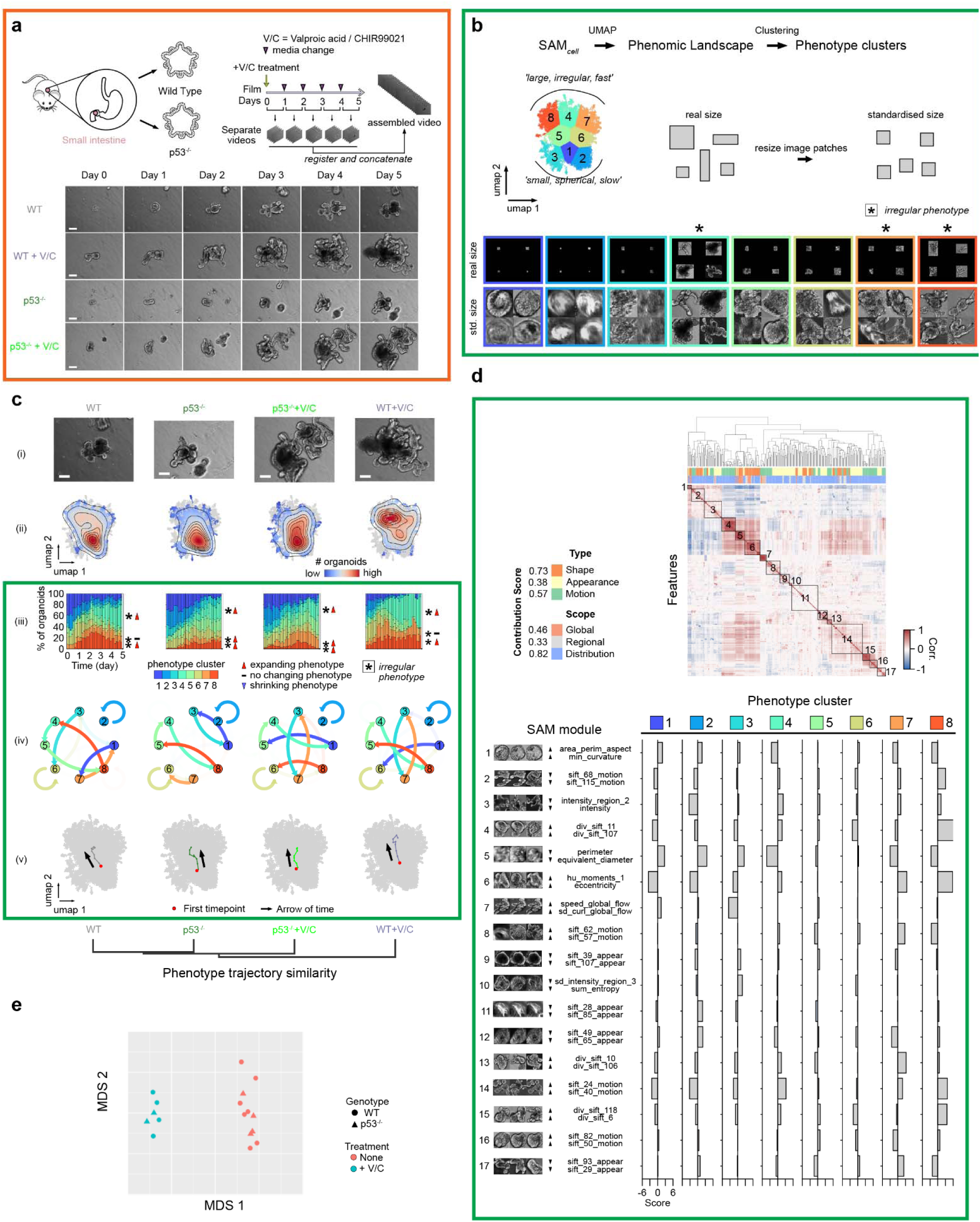
SPOT characterizes changes in murine small intestine organoid branching dynamics when treated with HDAC inhibitors. **a)** Small intestine organoids derived from WT p53^+/+^ and knockout p53^-/-^ mice were grown in 24-well plates. The organoids were left untreated or treated with a combination of valproic acid (V) and CHIR99021 (C) (+V/C), and then filmed every 15 minutes for 5 days. Media was changed every day, resulting in five separate video stacks per well, which were then registered frame-by-frame and concatenated into one video for analysis (top right). Snapshots of an example video from each condition on days 0–5 (bottom). Scalebar: 50 µm. **b)** UMAP phenomic landscape of filtered SAM phenomes from a total of 80 videos (20 per condition), 1-5 organoids in the field-of-view and carried out in 24-well plates, n=34,121 points. Each point corresponds to a segmented organoid instance from a single timepoint. Landscape is colored dark blue to dark orange and numbered in ascending order according to the mean shape eccentricity of each phenotype cluster identified by k-means clustering and the elbow method (top, left). For each cluster, four representative images arranged as a 2x2 grid is visualized, using their real size for comparison of scale across clusters (bottom, top row) and after cropping and resizing to a standardized (std.) size to show organoid appearance (bottom, bottom row). Illustration of real vs std. size (top right). An asterisk indicates the ‘irregular’ branching-relevant clusters. **c)** Hierarchically clustered treatment conditions according to UMAP SAM temporal phenotype trajectories with complete linkage (bottom). Row (i): final frame of example videos in **a)** for each condition. Scalebar: 50 µm. Row (ii): smoothed UMAP local point density heatmaps. Row (iii): stacked barplots showing the temporal changes in the phenotype cluster distribution of segmented organoid instances in consecutive 3 hour time intervals for each indicated condition. An asterisk indicates the ‘irregular’, branching-relevant clusters. Row (iv): graph showing the Hidden Markov Model (HMM) inferred transition probability (Methods) of an organoid transitioning to another phenotype cluster in the next timepoint given its phenotype cluster label at the current timepoint. Arrows are colored by the source cluster; the more transparent the arrow, the lower the probability of the transition. Row (v): UMAP SAM temporal phenotype trajectories for a density threshold of mean+3 standard deviations. Each colored point on a trajectory denotes half-day increments from the starting point (red point) on each trajectory. Black arrows indicate the directionality of time. **d)** Contribution score of shape, appearance and motion, and global, regional and distributional features (left), automated hierarchical clustering to identify SAM feature modules (right) and SAM module expression (labeled ‘score’ in barplot) in each phenotype cluster (bottom) constructed as in Fig. 2g. **e)** Multidimensional scaling (MDS) plot of the log_2_(CPM) expression of bulk RNA-seq counts from WT p53^+/+^ (circles) and knockout p53^-/-^(triangles) organoids with no treatment harvested at day 0 or harvested at day 5 after V/C treatment (+V/C), where CPM stands for transcript counts per million.

**Supplementary Figure 4a.**
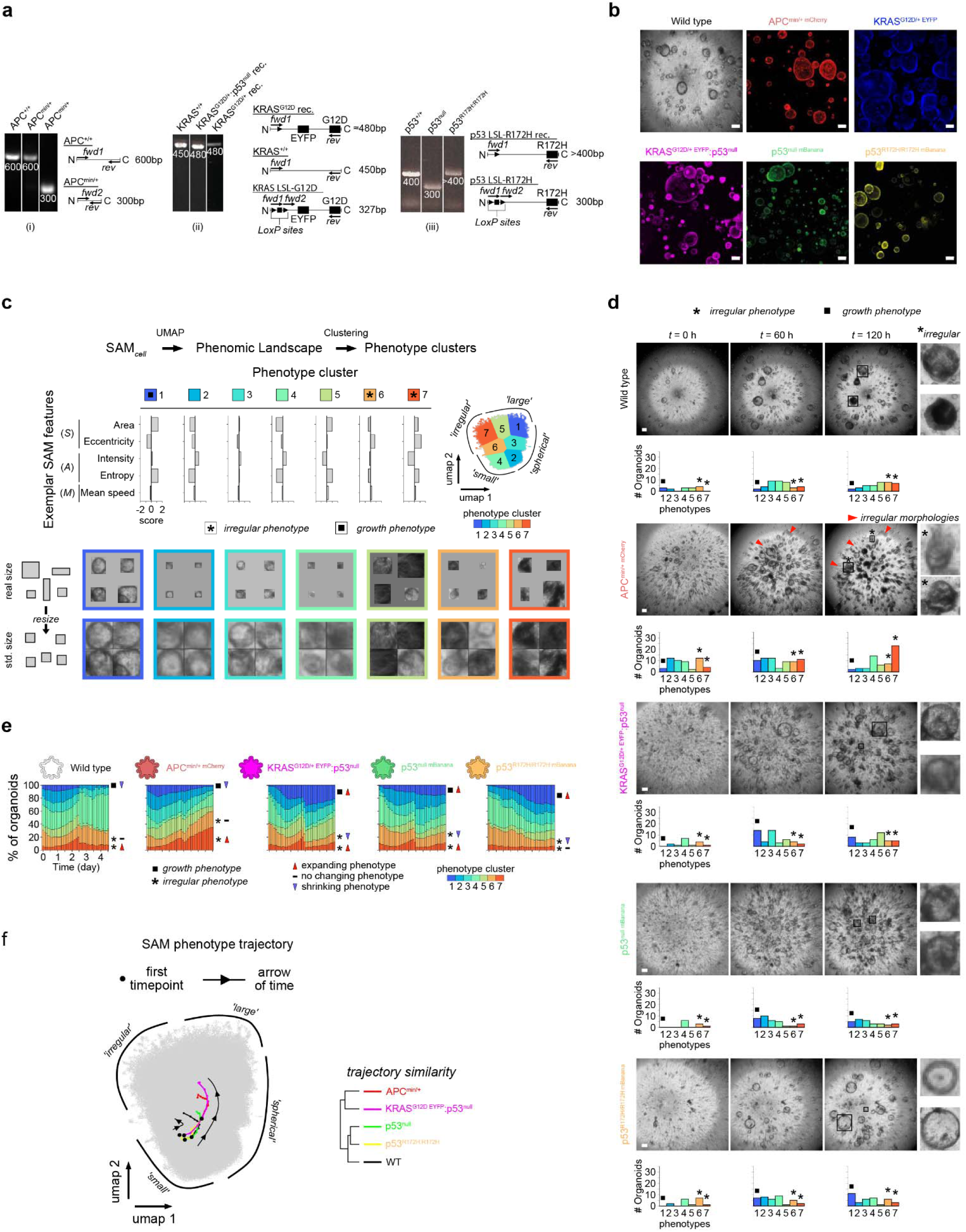
SPOT draws the same conclusions with label-free imaging of genotype defined murine colon organoids. **a)** PCR validation of mutant colon organoid construction for (i) APC^min/+^, (ii) KRAS^G12D/+^ and KRAS^G12D/+^:p53^null^ and (iii) p53^null^ and p53^R172H:R172H^. For each, the PCR gel with band base pair lengths labelled (left) and schematic of the primer design (right). See Suppl. Table 5 for primer sequences. **b)** Brightfield and confocal microscopy snapshots confirming fluorescent protein expression and correct organoid phenotype at day 7. Scalebars: 100 μm. **c)** K-means clustering of UMAP phenomic landscape of filtered SAM phenomes from 98 phase contrast videos, two independent experiments with organoids embedded in 10 μL Matrigel droplets in 96-well plates, total n=626,083 points. Each point corresponds to a segmented organoid instance at a single timepoint. Each cluster possesses distinct combinations of SAM features. Clusters are numbered in ascending order and are colored dark blue to dark orange with increasing shape eccentricity (top right). Barplots of mean SPOT score of representative, global SAM features for each k-means phenotype cluster (top left) and top four image exemplars below arranged as a 2x2 grid in both real and standardized (std.) sizes to illustrate the difference in scale and appearance of organoids across clusters (bottom). Irregular phenotypes are indicated by an asterisk and growth phenotypes are indicated by a black square. **d)** Example video snapshots of the phase contrast timelapse of wild type and mutant organoids. For each genotype, upper: the extended focus projection used for analysis, scalebar: 200µm; lower: number (#) of organoids classified as each of the seven k-means phenotype clusters of **c)**. APC^min/+^ organoids show elevated propensity to morphologically deform from the normal spherical shape of colon organoids into more irregular morphologies, as indicated by an asterisk and red triangle. Organoids outlined in images by a black square are digitally enlarged and displayed on the right. Growth and irregular phenotypes are indicated as in **c)**. **e)** Stacked barplot per genotype of the fraction of organoid instances in each phenotype cluster within consecutive 3 hour time intervals covering the 4.5 day acquisition. Beyond 2.5 days there is a discontinuous shift in cluster frequencies across most genotypes due to excess organoid death. **f)** UMAP SAM temporal phenotype trajectory of organoid growth for 0–2.5 days of **e)**. Each colored point denotes a 6 hour increment from the starting point (black point) on each trajectory. Black arrows show directionality of time. Average linkage hierarchical clustering of the depicted phenotype trajectories (right).

**Supplementary Figure 4b.**
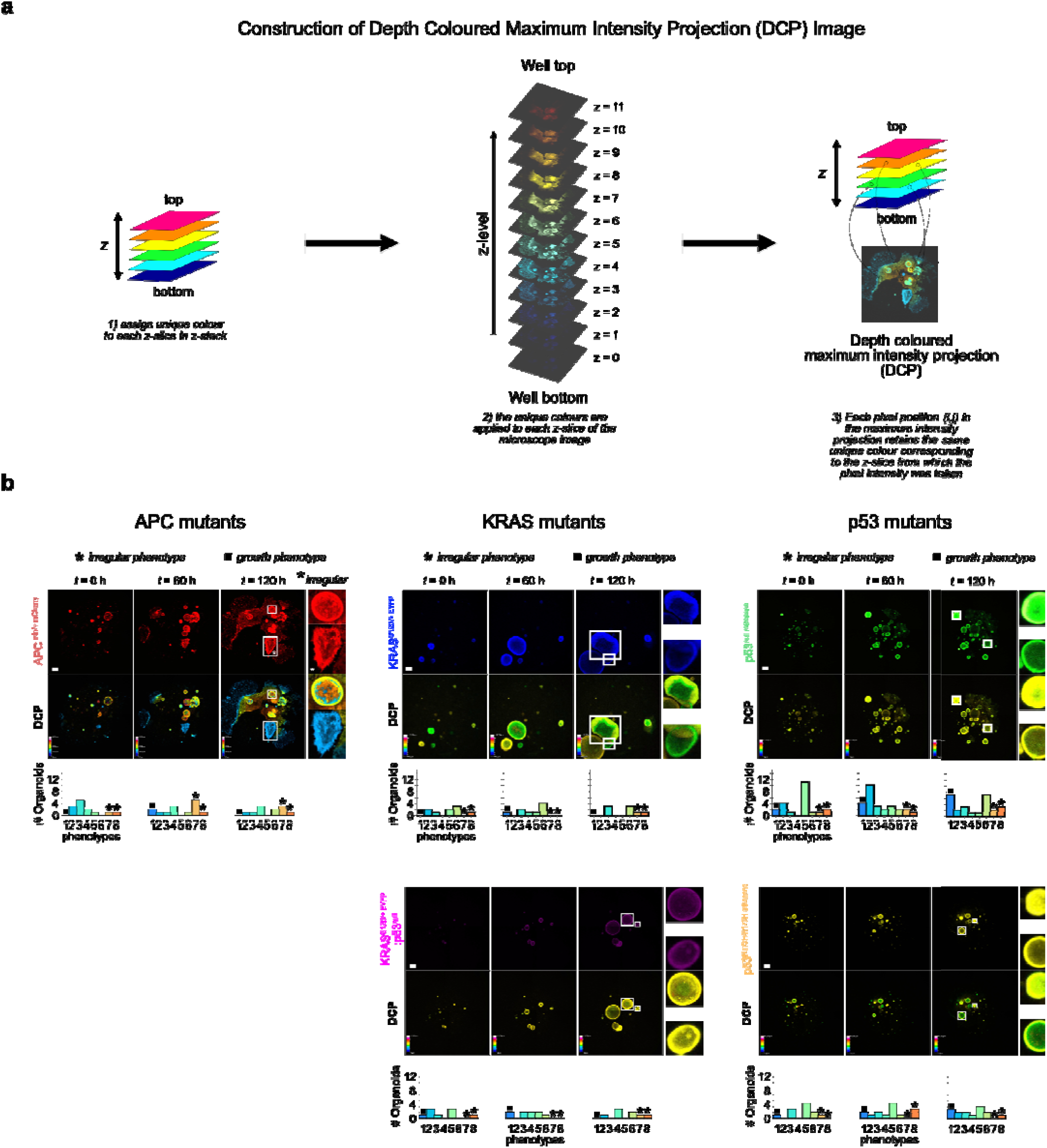
Murine colon organoids exhibit flattened morphologies preferentially at boundary of ECM dome. **a)** Schematic of computing a depth colored z-stack and its maximum intensity projection 2D image (DCP) to depict depth position of organoids in well. **b)** Representative DCP of mutant murine colon organoids at initial (*t*=0h), intermediate (*t*=60h) and final (*t*=120h) timepoints. For each mutant, top row: 2D projection image, middle row: corresponding DCP image, bottom row: phenotype cluster distribution of organoids in image. All scale bars: 200 μm.

**Supplementary Figure 5.**
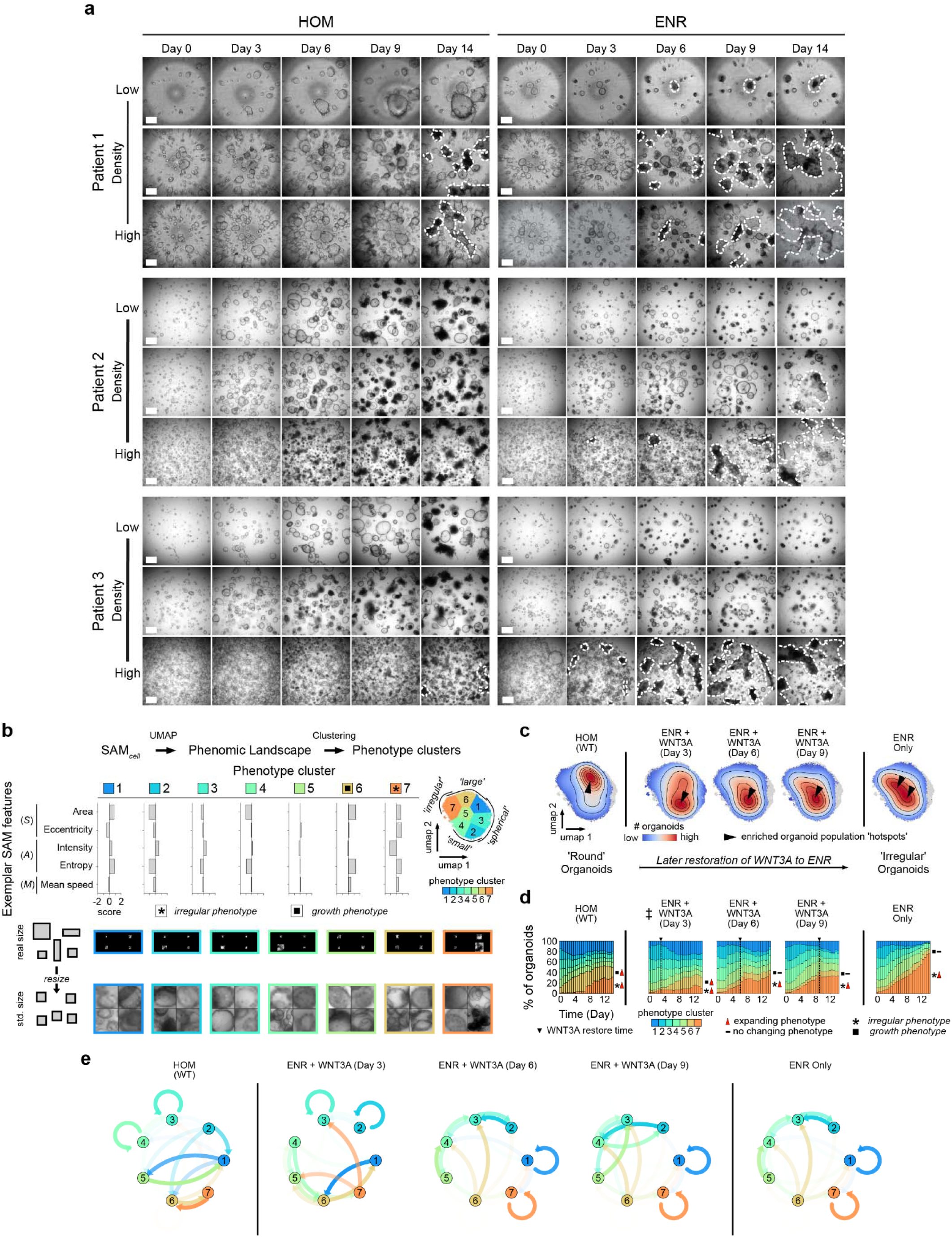
SPOT detects changes in WNT signaling-induced phenotypic dynamics in patient-derived duodenum organoids. **a)** Brightfield microscopy snapshots of patient-derived duodenum organoids for three patients; patient 1 (top), patient 2 (middle) and patient 3 (bottom). For each patient, snapshots for low (top panel), medium (middle panel) and high (bottom panel) organoid densities and cultured in HOM (left panels, day 0–14) and ENR (right panels, day 0–14) with different confluencies. Scale bar: 200 μm. White dashed lines outline the morphologically flattened organoids. The phase contrast filter was used for patient 1 and the brightfield filter was used for patient 2 and 3’s image collection. **b)** K-means clustering of UMAP phenomic landscape of filtered SAM phenomes from a total of 43 videos of organoids derived from three different patients, cultured and filmed in a 96-well plate, n=451,216 points (subsampled, every five organoid instances in the full compiled SAM table). Each point corresponds to a segmented organoid instance at a single timepoint. Corresponding barplots, representative image patches and UMAP phenomic landscape representations are constructed as in Fig. 4b. **c)** Smoothed UMAP local point density heatmaps, computed using all organoid instances in each respective culture condition. **d)** Stacked barplots showing the proportion of organoid instances in each phenotype cluster in **b)** over consecutive 12 hour intervals over the whole 0–14 day acquisition for each treatment condition. **e)** Graph showing the Hidden Markov Model (HMM) inferred transition probability (Methods) of an organoid transitioning to another phenotype cluster in the next timepoint given its phenotype cluster label in the current timepoint. Arrows are colored by the source cluster. The more transparent the arrow, the lower the probability of transition.

## SUPPLEMENTAL TABLES

**Supplementary Table 1.**
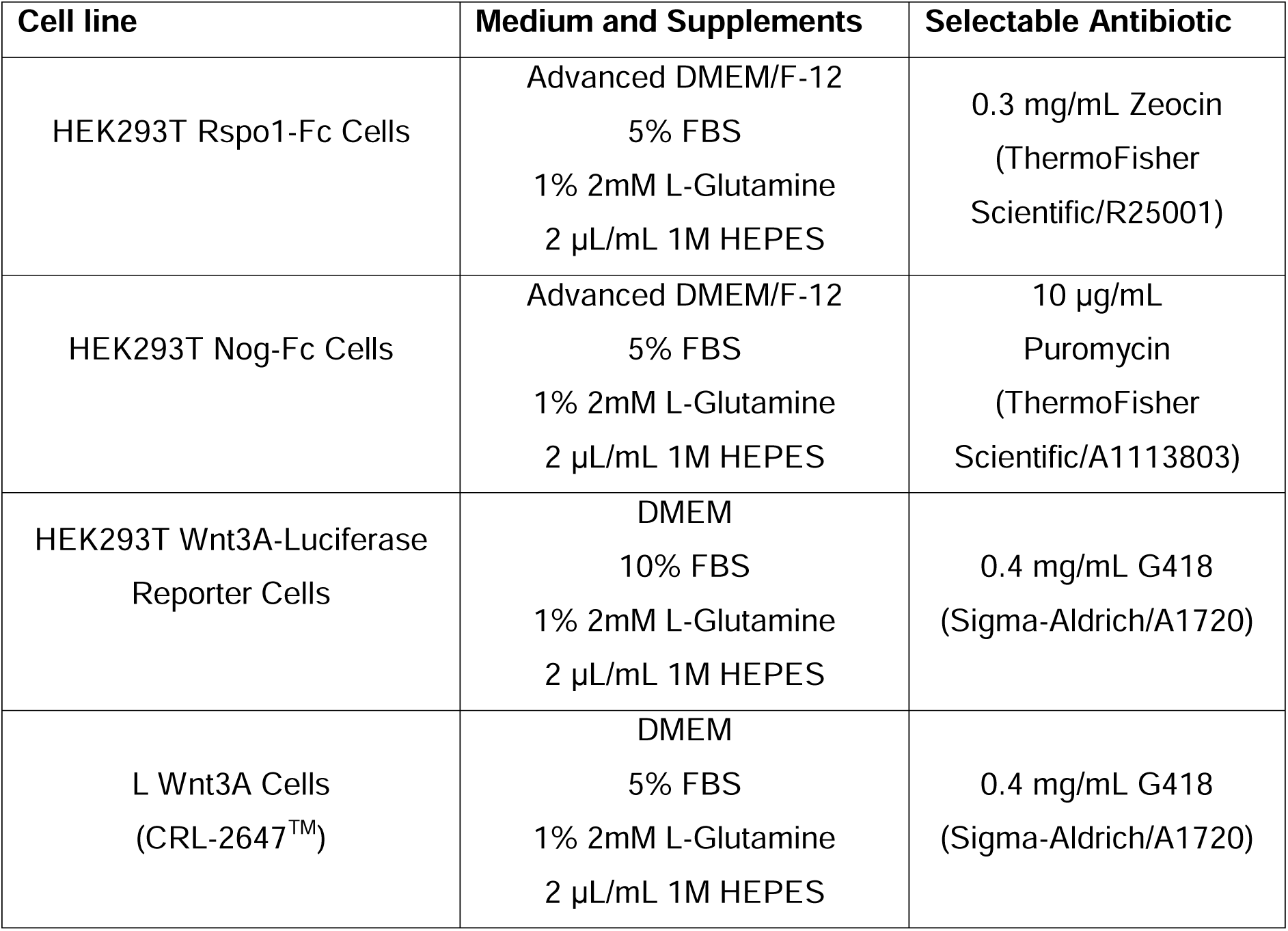
Cell lines and media requirements used for producing organoid conditioned media.

**Supplementary Table 2.**
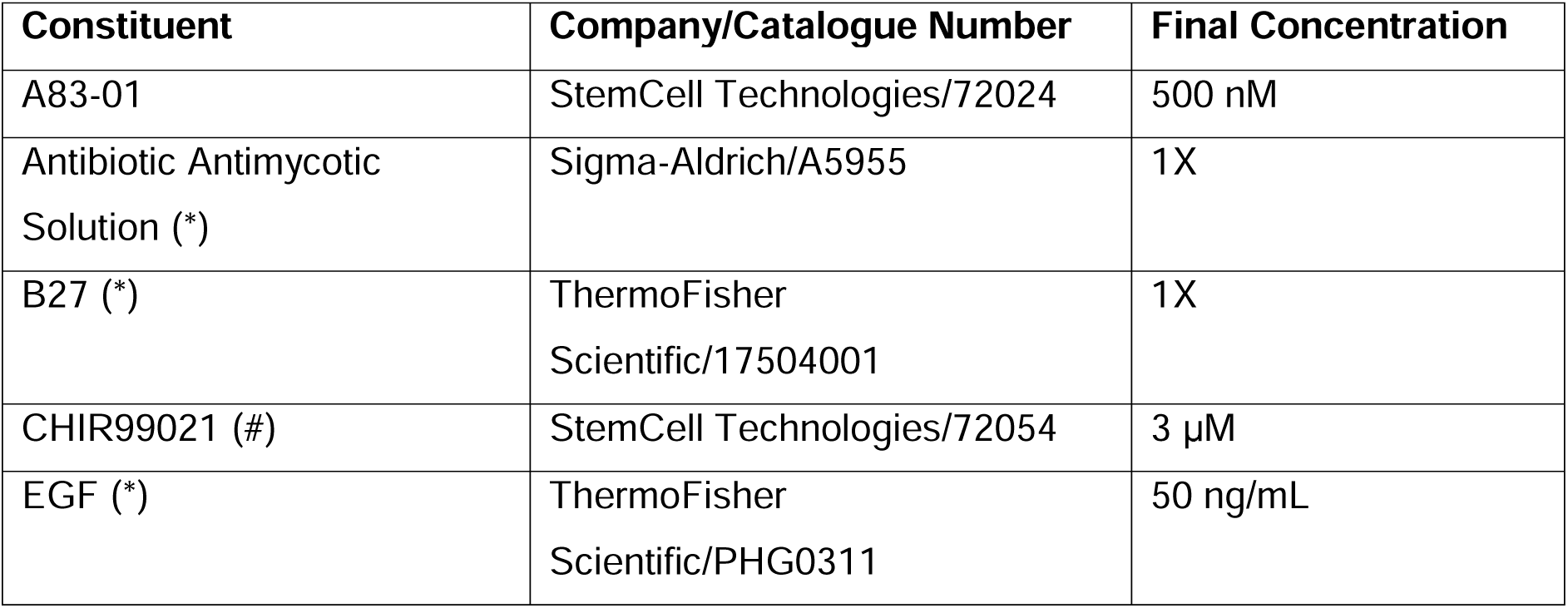
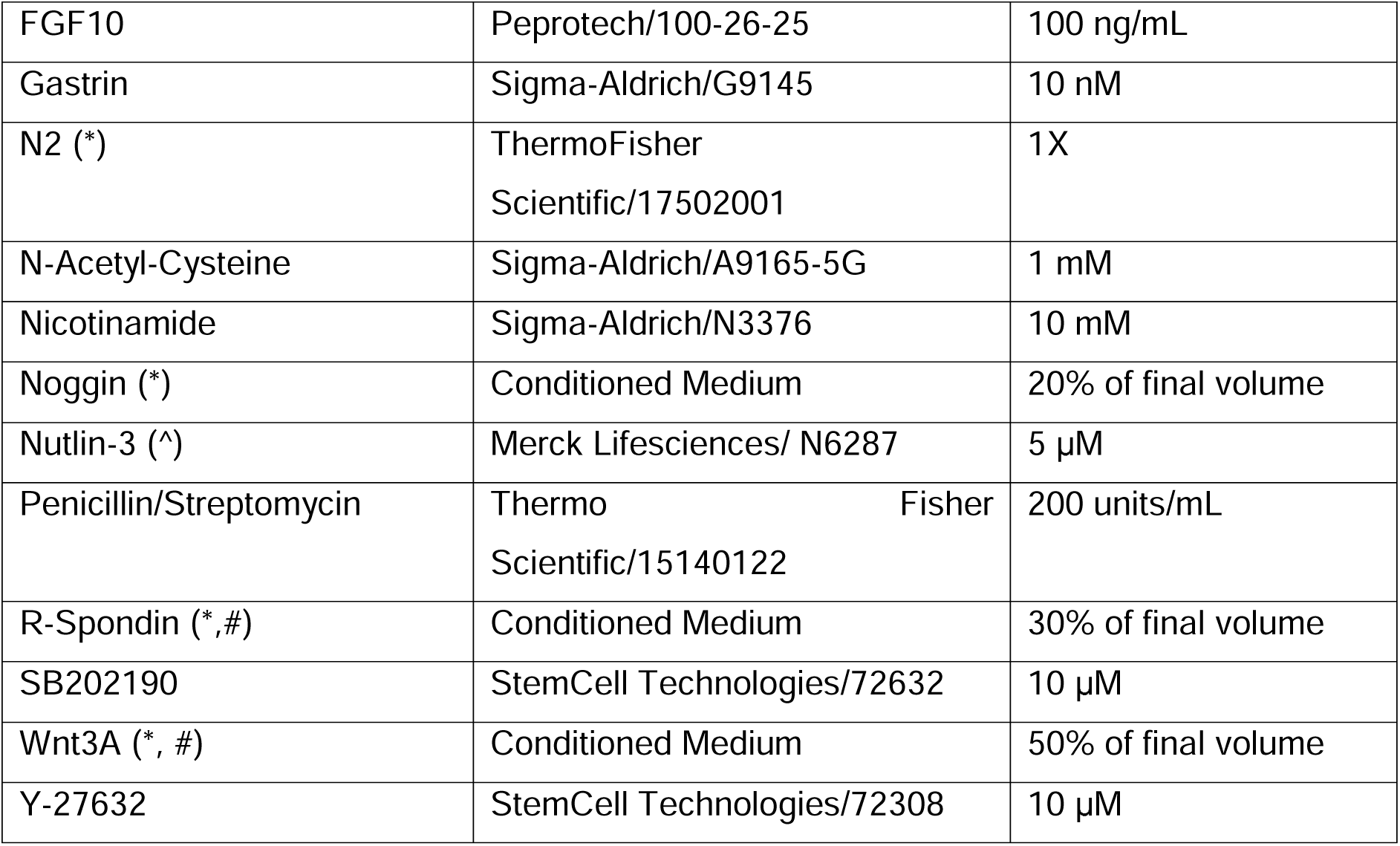
Constituents of the human and mouse organoid media and reagent concentrations. * denotes those constituents used in mouse organoid medium only, # denotes those constituents excluded from APC^min/+^ mouse organoid medium, ^ denotes constituents included in p53^R172H/R172H^ mouse organoid medium only.

**Supplementary Table 3.** Plate types and live cell imaging acquisition details for organoid experiments.

| Figure | Plate | Z-stack |
| --- | --- | --- |
| Figure 3 | Corning® Costar® TC-Treated Multiple Well Plates | Z-stack images 1000 µm in depth were acquired at 100 µm intervals (11 steps per acquisition) every hour for 5 days. |
| Figure 4 | 384-well plate, Flat bottom, rounded square wells | Z-stack images 1225 µm in depth were acquired at 35 µm intervals (35 steps per acquisition) every 2 hours for 5 days at each of the chosen positions. |
| Figure 5 | Nunc MicroWell 96-Well Optical-Bottom Plates with Polymer Base, Cell culture, black | Z-stack images 1000 µm in depth were acquired at 100 µm intervals (11 steps per acquisition) approximately every hour for 14 days. |
| Supp Fig 1a | Nunc MicroWell 96-Well Optical-Bottom Plates with Polymer Base, Cell culture, black | Z-stack images 1000 µm in depth were acquired at 100 µm intervals (11 steps per acquisition) approximately every 60 min for 14 days. |
| Supp Fig 1b | 384-well plate, Flat bottom, rounded square wells | Z-stack images 1225 µm in depth were acquired at 35 µm intervals (35 steps per acquisition) every 2 hours for 5 days at each of the chosen positions. |
| Supp Fig 3a.c | Nunc MicroWell 96-Well Optical-Bottom Plates with Polymer Base, Cell culture, black | Z-stack bright-field images covering 1000 µm total depth were acquired at 100 µm intervals (11 steps per acquisition) every 4 hours for 5 days. |
| Supp Fig 3b, Supp Fig 3c | “Plastic transparent” plates: Corning® Costar® TC-Treated Multiple Well Plates<br>“Optical-bottom” plates: Nunc MicroWell 96-Well Optical-Bottom Plates with Polymer Base, Cell culture, black | Z-stack bright-field images covering 1000 µm total depth were acquired at 100 µm intervals (11 steps per acquisition) every 4 hours for 5 days. |
| Supp Fig 3d | Corning® Costar® TC-Treated Multiple Well Plates | Z-stack images of 1000 µm in depth were acquired at 100 µm intervals (11 steps per acquisition) approximately every 15 minutes for 5 days. |
| Supp Fig 4a | Nunc MicroWell 96-Well Optical-Bottom Plates with Polymer Base, Cell culture, black | Z-stack bright-field images covering 1000 µm total depth were acquired at 100 µm intervals (11 steps per acquisition) every 4 hours for 5 days. |
| Supp Fig 5 | Nunc MicroWell 96-Well Optical-Bottom Plates with Polymer Base, Cell culture, black | Z-stack images 1000 µm in depth were acquired at 100 µm intervals (11 steps per acquisition) approximately every 60 min for 14 days. |

**Supplementary Table 4.**
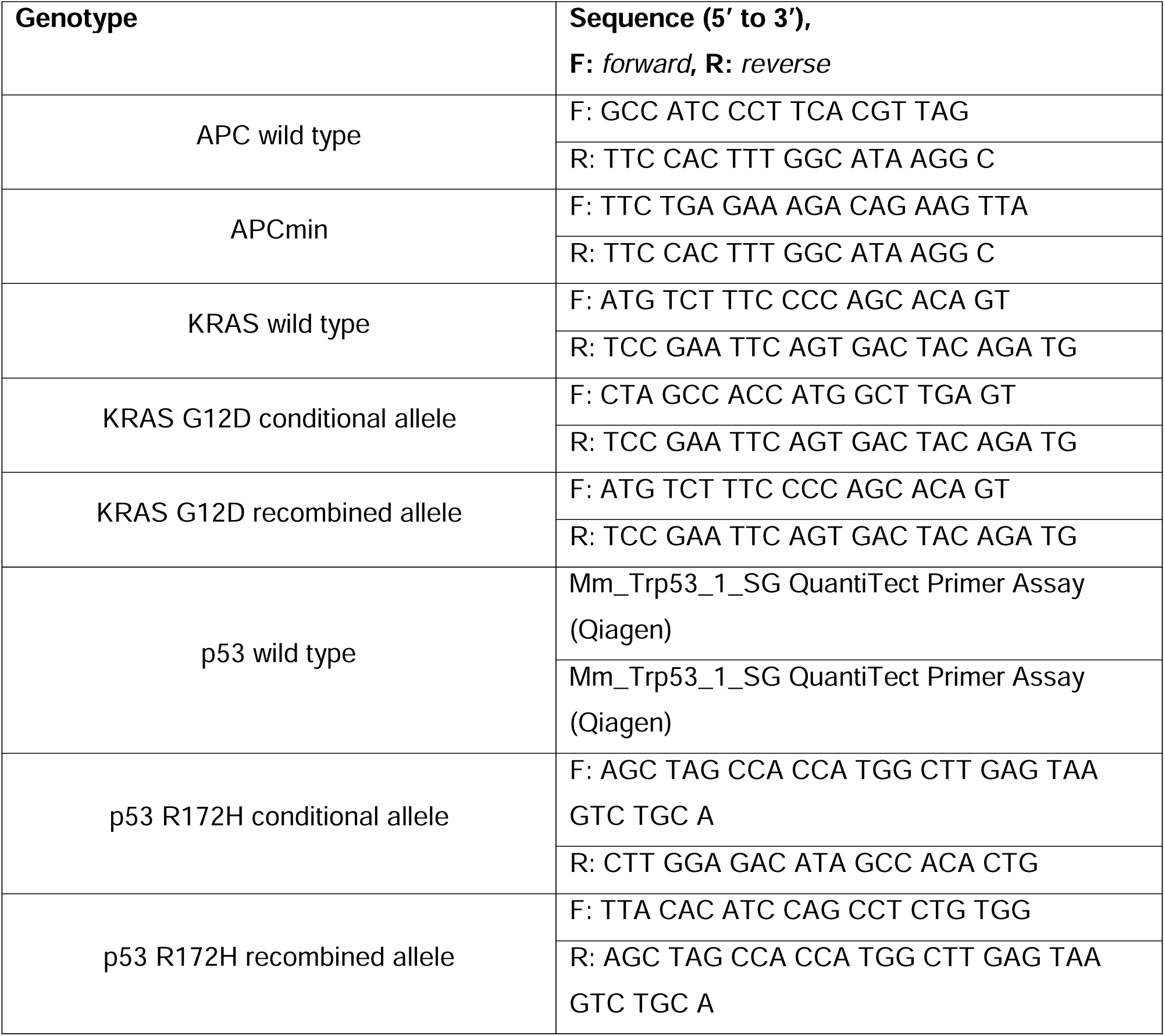
PCR primers used to validate the genotype of murine mutant colon organoids.

**Supplementary Table 5.** Metadata associated with simulated organoid datasets.

## SUPPLEMENTARY MOVIES

**Suppl. Movie 1. Tracking and segmentation of patient-derived duodenum organoids treated with different chemotherapy drugs.** One example video of organoids treated with each chemotherapy drug at the highest concentration and dimethyl sulfoxide (DMSO) control. For each treatment condition we firstly show the input spatiotemporally registered video; secondly, the bounding box detection and tracked individual organoids; and thirdly, the final filtered and segmented individual organoid boundaries. The current centroid of individual organoids is denoted by a white dot. Bounding box or contour color-matched dots denote historical centroid positions of the same organoid. The final boundaries remove all organoid instances near the border of the field of view to exclude partially segmented organoids from downstream phenotype analysis. Videos are organized such that the top and bottom row represent less toxic and more toxic drugs respectively, corresponding to the left and right clade of the hierarchical clustering in Fig. 2. Scalebar: 250 μm.

**Suppl. Movie 2. Tracking and segmentation of wild type and p53^-/-^ mouse small intestine organoids with and without V/C (valproic acid/CHIR99021) treatment.** One example video of wild type (p53^+/+^) and p53^-/-^ organoid cultures, with and without V/C treatment. For each treatment we firstly show the input spatiotemporally registered video; secondly, the bounding box detection and tracked individual organoids; and thirdly, the final filtered and segmented individual organoid boundaries. The current centroid of individual organoids is denoted by a white dot. Bounding box or contour color-matched dots denote historical centroid positions of the same organoid. The final boundaries remove all organoid instances near the border of the field of view to exclude partially segmented organoids from downstream phenotype analysis. Scalebar: 50 μm.

**Suppl. Movie 3. Tracking and segmentation of murine mutant colon organoids.** One example video of organoid culture for each mutant genotype; APC^min/+ mCherry^, KRAS^G12D/+ EYFP^:p53^null^, KRAS^G12D/+^, p53^null^ ^mBanana^, p53^R172H/R172H^ ^mBanana^. For each mutant we firstly show the input spatiotemporally registered video; secondly, the bounding box detection and tracked individual organoids; and thirdly, the final filtered and segmented individual organoid boundaries. The current centroid of individual organoids is denoted by a white dot. Bounding box or contour color-matched dots denote historical centroid positions of the same organoid. The final boundaries remove all organoid instances near the border of the field of view to exclude partially segmented organoids from downstream phenotype analysis. Scalebar: 200 μm.

**Suppl. Movie 4. Tracking and segmentation of patient-derived duodenum organoids in human organoid media (HOM) vs ENR media only and with WNT3A restoration, from three patients.** One example video of organoid culture in normal human organoid (HOM), ENR only media, and ENR with WNT3A restoration after the specified number of days from the start of the timelapse for each of the three analyzed patients. For each patient, for each condition, we firstly show the input spatiotemporally registered video; secondly, the bounding box detection and tracked individual organoids; and thirdly, the final filtered and segmented individual organoid boundaries. The current centroid of individual organoids is denoted by a white dot. Bounding box or contour color-matched dots denote historical centroid positions of the same organoid. The final boundaries remove all organoid instances near the border of the field of view to exclude partially segmented organoids from downstream phenotype analysis. Scalebar: 200 μm.

